# mTOR-regulated Mitochondrial Metabolism Limits Mycobacterium-induced Cytotoxicity

**DOI:** 10.1101/2022.01.30.478369

**Authors:** Antonio J. Pagán, Lauren J. Lee, Joy Edwards-Hicks, Cecilia B. Moens, David M. Tobin, Elisabeth M. Busch-Nentwich, Erika L. Pearce, Lalita Ramakrishnan

## Abstract

Necrosis of macrophages in the tuberculous granuloma represents a major pathogenic event in tuberculosis. Through a zebrafish forward genetic screen, we identified the mTOR kinase, a master regulator of metabolism, as an early host resistance factor in tuberculosis. We found that mTOR complex 1 protects macrophages from mycobacterium-induced death by enabling infection-induced increases in mitochondrial energy metabolism fueled by glycolysis. These metabolic adaptations are required to prevent mitochondrial damage and death caused by the secreted mycobacterial virulence determinant ESAT-6. Thus, the host can effectively counter this early critical mycobacterial virulence mechanism simply by regulating energy metabolism, thereby allowing pathogen-specific immune mechanisms time to develop. Our findings may explain why *Mycobacterium tuberculosis*, albeit humanity’s most lethal pathogen, is successful in only a minority of infected individuals.

## INTRODUCTION

*Mycobacterium tuberculosis* (Mtb) induces the formation of granulomas, organized structures comprised of macrophages within which mycobacteria reside, and accessory cells (Pagan and Ramakrishnan, 2014; 2018; Ramakrishnan, 2012). The granuloma represents a key host-pathogen battleground that determines the outcome of tuberculosis (TB) infection (Pagan and Ramakrishnan, 2014; Ramakrishnan, 2012). In most individuals, the granuloma successfully clears Mtb infection, often leaving a residual, sterile, fibrotic structure as a stamp of past infection (Behr et al., 2018; 2019; Behr et al., 2021; Canetti et al., 1972; Feldman and Baggenstoss, 1938; Opie and Aronson, 1927; Terplan, 1951). In contrast, in the minority of people who go on to develop TB, the granuloma often becomes a mycobacterium-beneficial structure that promotes bacterial expansion and dissemination (Pagan and Ramakrishnan, 2014; Ramakrishnan, 2012). Granuloma necrosis is a pivotal pathogenic event because it delivers macrophage-resident mycobacteria into the growth-enhancing extracellular milieu (Pagan and Ramakrishnan, 2014; Ramakrishnan, 2012; Russell, 2007). Necrosis of lung granulomas with their attendant rupture into the airways also facilitates transmission, sustaining the global TB burden and promoting Mtb’s evolutionary survival (Ong et al., 2014; Ramakrishnan, 2012). Mtb’s exploitation of the granuloma begins in the innate stage of the response, enabling Mtb to gain a foothold in the host (Cambier et al., 2014a; Pagan and Ramakrishnan, 2014; Ramakrishnan, 2012). Thus, innate immune dysfunctions that facilitate this exploitation can lead to susceptibility (Cambier *et al*., 2014a; Pagan and Ramakrishnan, 2014; Ramakrishnan, 2012).

Zebrafish infected with *Mycobacterium marinum* (Mm), a close relative of Mtb, develop TB-like disease with necrotic granulomas (Cosma et al., 2003; Swaim et al., 2006). Mm-infected zebrafish larvae, in which adaptive immunity has not yet developed, also form organized granulomas that undergo necrosis (Davis et al., 2002). Thus, the zebrafish larva offers the opportunity to dissect the contribution of innate immunity in tuberculous granuloma formation and necrosis (Davis *et al*., 2002; Davis and Ramakrishnan, 2009; Ramakrishnan, 2013; 2020). The larva’s optical transparency and genetic tractability enables the detailed, sequential monitoring by intravital microscopy of the steps of infection, the contribution of host and pathogen determinants to them, and the consequences to outcome (Ramakrishnan, 2013). The use of unbiased genetic screens and candidate gene approaches in zebrafish larvae has shed light on granuloma biology (Ramakrishnan, 2013). Hypersusceptible zebrafish mutants displaying accelerated granuloma necrosis have identified innate immune host determinants that protect against necrosis and that are relevant to human TB (Berg et al., 2016; Clay et al., 2008; Pagan et al., 2015; Roca and Ramakrishnan, 2013; Roca et al., 2022; Roca et al., 2019; Tobin et al., 2012; Tobin et al., 2010; Whitworth et al., 2021a; Whitworth et al., 2021b).

Here, we report on insights into TB pathogenesis and resistance gained from the genetic mapping and characterization of a loss-of-function mutant in mTOR kinase that exhibits rapid granuloma necrosis. The mTOR pathway integrates environmental signals emanating from diverse nutrient-sensing and growth factor receptor pathways to regulate biosynthetic and metabolic processes vital for cellular development, growth, survival, and function (Liu and Sabatini, 2020). Iterative experimental approaches in the zebrafish and in human macrophages uncover the mitotoxic function of mycobacterial ESAT-6 and show that mTOR-facilitated mitochondrial metabolism serves as a highly effective innate ‘counter virulence’ factor in TB by exerting a mitoprotective effect against this critical mycobacterial virulence factor.

## RESULTS

### mTORC1 Deficiency Confers Susceptibility to Mm Infection in Zebrafish

The zebrafish mutant *fh178*, identified in a forward genetic screen (Tobin *et al*., 2010), was hypersusceptible to Mm, with larvae exhibiting increased bacterial burdens relative to wild-type and heterozygote siblings following infection into the caudal vein (cv) (Figures 1A-1C). By four days post-infection (dpi), *fh178* mutant granulomas had depleted their macrophages and the released mycobacteria were growing extracellularly in rope-like cords (Pagan *et al*., 2015; Tobin *et al*., 2010)(Figure 1D and 1E). Mycobacterial cording, a sensitive and specific surrogate for macrophage depletion, is a readily quantifiable, binary phenotype for mapping mutants rendered hypersusceptible by granuloma necrosis (Berg *et al*., 2016; Tobin *et al*., 2010). Using polymorphic markers and the mycobacterial cording phenotype, we mapped the *fh178* mutation to a nonsense mutation in exon 24 of the *mtor* gene (See STAR Methods). We confirmed that genetic disruption of *mtor* was the cause of *fh178* hypersusceptibility by showing that animals with a nonsense mutation in exon 19 (*mtor*^*sa16755*^) (Kettleborough et al., 2013) also exhibited hypersusceptibility with cording as did compound *fh178/sa16755* heterozygotes (Figure 1F and 1G).

**Figure 1.**
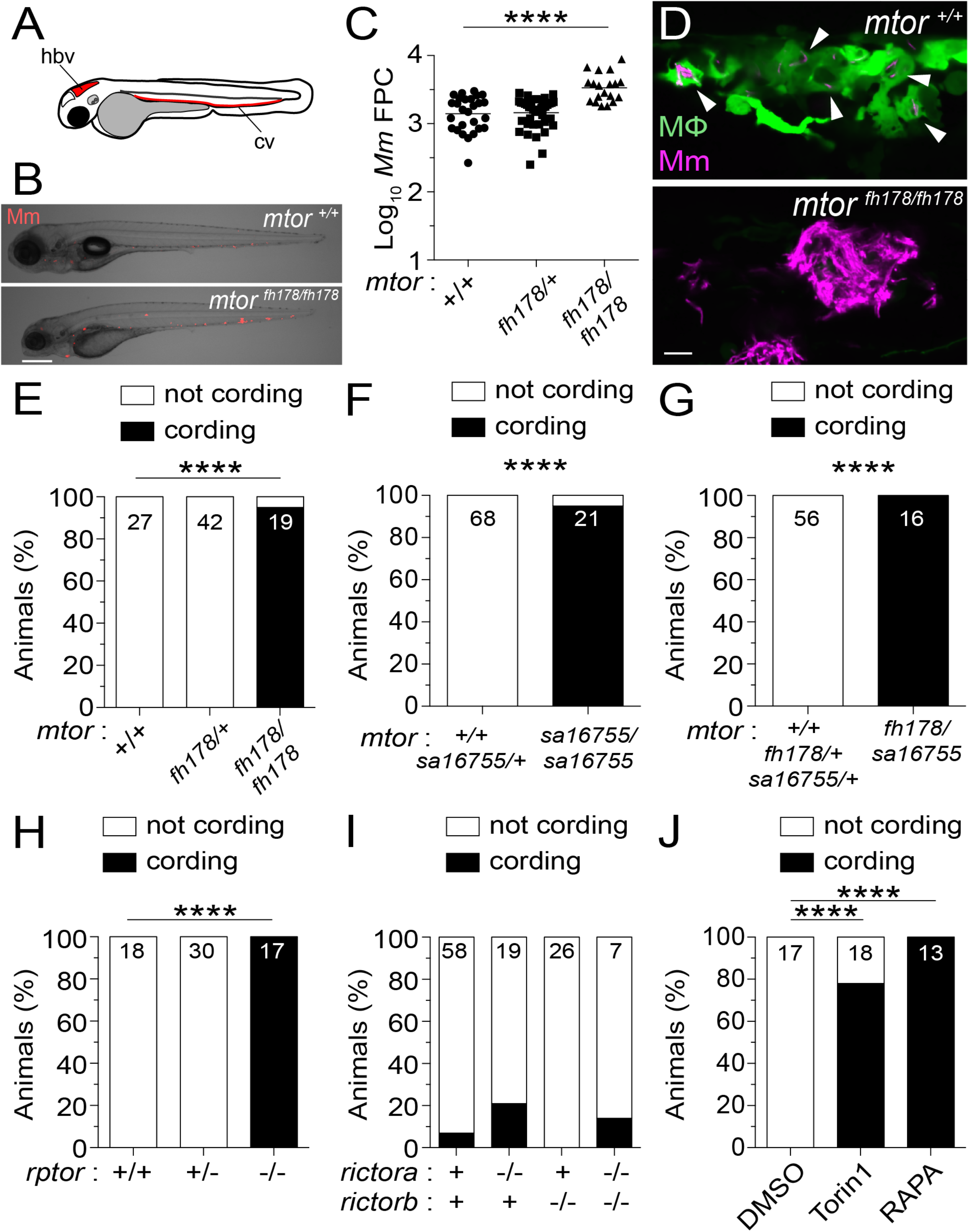
mTORC1-deficient Zebrafish Are Hypersusceptible to Mm Infection. (A) Hindbrain ventricle (hbv) and caudal vein (cv) injection routes used in this study. Larvae were infected with ∼150 Mm expressing tdTomato (B, C, E - J) or tdKatushka2 (D) fluorescent proteins via the caudal vein two days post-fertilization (dpf). (B) Overlaid micrographs of widefield mycobacterial fluorescence (Mm, red) and brightfield in *mtor*^*fh178/fh178*^ or wild-type siblings (*mtor*^*+/+*^) four days post infection (dpi). (C) Quantification of bacterial fluorescence (fluorescent pixel counts, FPC) in animals from *mtor*^*fh178/+*^ incross 4 dpi. Symbols represent individual animals. Horizontal lines indicate mean values. (D) Confocal micrograph optical sections of *mtor*^*fh178/fh178*^ and a wild-type sibling expressing *Tg(mpeg1:YFP)* 4 dpi showing a granuloma in the wild-type animal and mycobacterial cording in the *mtor*^*fh178/fh178*^ animal. Mm (magenta) and macrophages (green) are shown. Arrowheads indicate intracellular Mm. Mycobacterial cording in animals from (E) *mtor*^*fh178/+*^ incross, (F) *mtor*^*sa16755/+*^ incross, (G) *mtor*^*fh178/+*^ x *mtor*^*sa16755/+*^ cross, and (H) *rptor*^*sa11537/+*^ incross at 4 dpi, and (I) *rictora*^*sa15967/+*^; *rictorb*^*sa18403/+*^ double heterozygote incross and (J) wild-type animals treated with torin1 (400 nM), rapamycin (400 nM), or 0.5% DMSO (vehicle control) 5 dpi. (E - J) Numbers within columns indicate animals per group. Scale bars, (B) 300 µm and (D) 25 µm. Statistical analyses, (C) one-way ANOVA with Tukey’s post-test and (E - J) Fisher’s exact test. Data are representative of two or more independent experiments.

The mTOR kinase functions in two distinct complexes, mTORC1 and mTORC2, that require the adaptors Raptor and Rictor, respectively (Liu and Sabatini, 2020). Animals with nonsense alleles of *rptor*, the gene encoding Raptor, showed similar cording as the *mtor* mutants, whereas those with nonsense alleles of *Rictor* did not (Figure 1H and 1I). Inhibition of mTOR or mTORC1 with torin1 or rapamycin, respectively (Benjamin et al., 2011), recapitulated genetic mTOR/mTORC1 deficiency with increased cording (Figure 1J). Thus, mTORC1 deficiency confers susceptibility to mycobacterial infection, linked to early granuloma breakdown. Because zebrafish larvae have not yet developed adaptive immunity, this reflects innate resistance conferred by mTOR.

### mTOR Deficiency Accelerates Death of Mycobacterium-Infected Macrophages

At 4 dpi, wild-type animals had granulomas with sparse intracellular bacteria; in contrast mTOR-deficient animals had clusters of extracellular bacteria in shapes similar to wild-type granulomas, suggesting that they had been in granulomas that had broken down due to macrophage death (Figure 1D, compare top and bottom). To detail the kinetics of macrophage death, we infected zebrafish larvae in the hindbrain ventricle (hbv), an acellular compartment ideal for monitoring early granuloma formation (Davis and Ramakrishnan, 2009). Animals rendered mTOR-deficient by rapamycin treatment formed granulomas similar to wild-type by 2 dpi, but their macrophages died by 3 dpi, leaving clumps of extracellular bacteria (Figure 2A).

**Figure 2.**
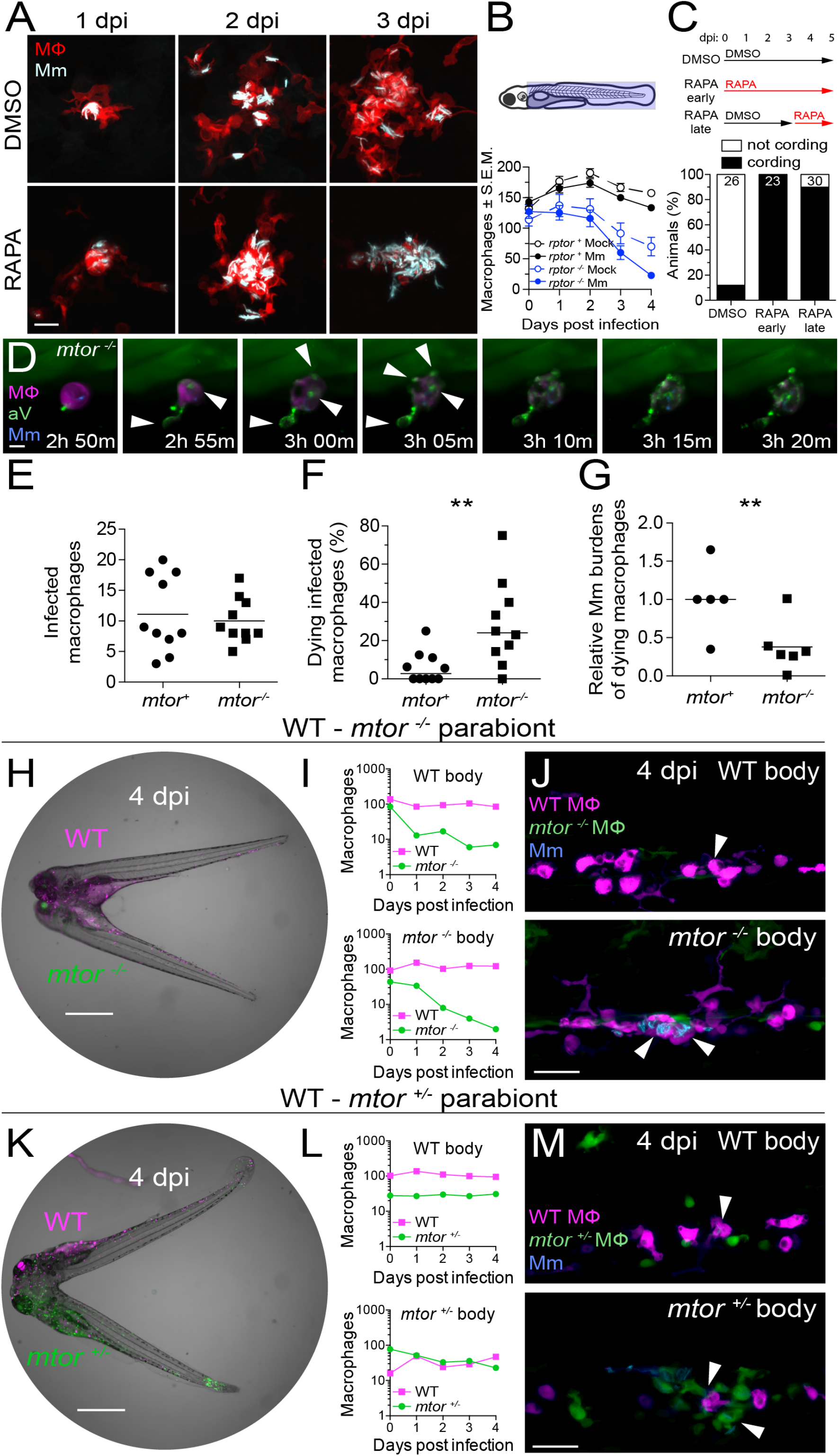
mTOR Deficiency Impairs Macrophage Development and Survival and Sensitizes Infected Macrophages to Mycobacterium-induced Cytotoxicity. Larvae were infected with Mm expressing BFP2 (A, E, I - N), mWasabi (C, D) or tdKatushka2 (F - H) fluorescent proteins via the hindbrain ventricle (A) or the caudal vein (B - N) 2 dpf. (A) Serial confocal micrographs of granulomas in *Tg(mfap4:tdTomato-CAAX)* zebrafish treated with rapamycin or DMSO. Mm (cyan), macrophages (red). (B) (Top) Macrophage counting region (shaded light blue). (Bottom) Numbers of macrophages in Mm*-* and mock-infected Raptor mutants and siblings expressing *Tg(mpeg1:tdTomato)*. Symbols indicate mean values for each group. Error bars show Standard Error of the Mean (S.E.M.). (C) (Top) Duration of rapamycin and DMSO treatments. (Bottom) Mycobacterial cording 5 dpi. (D) Time-lapse confocal micrographs of a dying infected macrophage in an *mtor*^*sa16755/sa16755*^; *Tg(mfap4:tdTomato-CAAX); Tg(ubb:secA5-YFP)* animal 2 dpi. Mm (blue), secreted annexin V-YFP (green), macrophage (magenta), annexin V^+^ blebs (arrowheads). See Movie S1. (E - G) 6-hr time-lapse confocal microscopy of *mtor*^*fh178/fh178*^ and mTOR-sufficient siblings expressing *Tg(mpeg1:YFP)* 2 dpi. See Movie S2. (E) Absolute numbers of infected macrophages per field. (F) Percentage of dying macrophages per field. (G) Relative mycobacterial burdens in dying macrophages of *mtor*^*-/-*^ and mTOR-sufficient fish. Bacterial volumes were normalized to values obtained from dying cells in mTOR-sufficient controls for each imaging run. (H) Widefield micrograph of parabiotic zebrafish comprised of conjoined wild-type *Tg(mpeg1:tdTomato)* and *mtor*^*fh178/fh178*^; *Tg(mpeg1:YFP)* embryos 4 dpi. (I) Absolute numbers of macrophages in the WT body (top) and *mtor*^*-/-*^ body (bottom) of WT - *mtor*^*-/-*^ parabiont. (J) Maximum intensity projections of infections in the WT body (top) and *mtor*^*-/-*^ body (bottom) of a WT - *mtor*^*-/-*^ parabiont 4 dpi. (K) Widefield micrograph of wild-type *Tg(mpeg1:tdTomato)* and *mtor*^*fh178/+*^; *Tg(mpeg1:YFP)* parabiont 4 dpi. (L) Absolute numbers of macrophages in the WT body (top) and *mtor*^*+/-*^ body (bottom) of WT - *mtor*^*+/-*^ parabiont. (M) Maximum intensity projections of infections in the WT body (top) and *mtor*^*+/-*^ body (bottom) of a WT - *mtor*^*+/-*^ parabiont 4 dpi. Scale bars, (A) 25 µm, (D) 10 µm, (H, K) 400 µm, and (J, M) 50 µm. Statistical analyses, (E - G) two-tailed, unpaired Student’s *t* test. Time lapse data were pooled from five (E and F) or three (G) independent experiments. See also Figure S1.

We hypothesized that mTORC1 deficiency might cause granuloma breakdown through its impairment of myelopoiesis (Karmaus et al., 2017; Lee et al., 2017). The paucity of available macrophages to replenish the growing granuloma could cause its breakdown, similar to the case of myeloid growth factor Colony Stimulating Factor-1 Receptor (CSF-1R) deficiency (Pagan *et al*., 2015). Supporting this possibility, mTORC1 promotes myelopoiesis through CSF-1R (Karmaus *et al*., 2017). However, even though mTORC1-deficient animals had more macrophages at baseline than CSF-1R-deficient animals, their granulomas broke down sooner (2-4 days versus 5-7 for CSF-1R mutants) (Figure 2B)(Pagan *et al*., 2015), suggesting that mTOR deficiency induces death of infected granuloma macrophages independently of reducing basal macrophage supply. In support of this, rapamycin treatment after formation of granulomas caused their rapid breakdown (Figure 2C). For further confirmation, we used time-lapse microscopy to capture in real-time the death of infected macrophages. If mTOR-deficient granulomas are breaking down due to reduced macrophage replenishment, then dying mTOR mutant macrophages should have bacterial burdens similar to or greater than wild-type. To assess bacterial burdens in dying macrophages, we used blue fluorescent Mm to infect mTOR-deficient animals and their wild-type siblings. that had red fluorescent macrophage membranes. All animals were also transgenic for a ubiquitously-expressed secreted annexin V tagged with yellow fluorescent protein (*Tg(ubb:secA5-YFP)*), which accumulates on the surface of cells undergoing apoptosis and other modes of regulated cell death (Bendall and Green, 2014). We monitored macrophage death by the appearance of annexin V-YFP labeling of the plasma membrane and membrane blebs followed by the loss of tdTomato fluorescence reflecting plasma membrane disintegration (Figure 2D and Movie S1). In 2 dpi animals, over a 4.5-hour imaging period that captured similar numbers of infected macrophages in mTOR-deficient and wild-type siblings, 6-fold more mTOR-deficient macrophages died (Figure 2E and 2F and Movie S2). Importantly, mTOR-deficient macrophages died with lower bacterial burdens than wild-type (Figure 2G). This finding ruled out numerical macrophage defects as the cause of macrophage death and showed that mTOR-deficient macrophages died because they were “intolerant” of mycobacterial infection.

To determine if mTOR’s protective effect was macrophage-intrinsic, we compared mTOR mutant and wild-type macrophages in the same environment by creating wild-type – mTOR mutant parabionts, with differentially labeled macrophages, red and yellow fluorescent in wild-type and mTOR mutants, respectively, and infecting both caudal veins (Figure 2H). By 4 dpi, mTOR mutant macrophages had been depleted by more than 90% on both wild-type and mTOR mutant sides of the parabionts, whereas wild-type macrophages persisted equally on both sides (Figure 2I). The depletion of mTOR mutant macrophages in these animals should not lead to Mm cording because the complement of WT macrophages should be able to engulf the infected mTOR mutant corpses and keep Mm intracellularly. By 4 dpi, as predicted, Mm was only found inside the WT macrophages (Figure 2J). In mTOR heterozygote – wild-type parabionts, mTOR heterozygote yellow fluorescent macrophages survived as well as wild-type red fluorescent macrophages, ruling out *Tg(mpeg1:YFP)* expression as an artifactual cause of macrophage depletion of mTOR mutant macrophages (Figure 2K - 2L).

Thus, mTOR confers early cell-intrinsic protection against *Mycobacterium*-induced death.

### Infected Macrophage Death in mTOR Deficiency is Associated with Reduced Mitochondrial Membrane Potential

Impairment in nutrient sensing pathways caused by mTOR-deficiency could sensitize cells to autophagic death or mitochondrial apoptosis (Gonzalez et al., 2020; Green and Levine, 2014; Hosoi et al., 1999; Muthukkumar et al., 1995). To assess autophagic death, we created zebrafish deficient in the essential autophagy protein ATG12 (Bento et al., 2016). ATG12-deficient animals had defective autophagosome formation as evidenced by reduced aggregation of LC3 (Figures S1A and S1B). However, ATG12 deficiency did not prevent death of mycobacterium-infected macrophages in rapamycin-treated animals (Figure S1C). To assess mitochondrial apoptosis, we used caspase-9 deficient zebrafish mutants (Galluzzi et al., 2016). Capase-9 mutants had the expected defect in developmental apoptosis as reflected by reduced dead cell debris in the brain at 3 dpf (S1D and S1E). However, rapamycin treatment induced death of their infected macrophages (Figure S1F). Thus, mTOR deficiency kills infected macrophages independent of inducing autophagy or mitochondrial apoptosis.

We ruled out two other modes of macrophage death commonly associated with mycobacterial infection – inflammasome-dependent mediated through the adapter ASC and Type 1 interferon-dependent mediated through the cytosolic DNA sensing adapter STING (Decout et al., 2021; Swanson et al., 2019). ASC or STING mutants had increased macrophage death when rendered mTOR-deficient (Figure S2G and S2H).

In contrast to mTOR deficiency, where macrophages died rapidly when bacterial burdens were still low, mycobacterium-induced macrophage death typically occurs with high intracellular bacterial burdens (Amaral et al., 2019; Beckwith et al., 2020; Clay *et al*., 2008; Lee et al., 2011), We had previously identified a case of macrophage death associated with low mycobacterial burdens, which was mediated through a pathway activated by dysregulated Tumor Necrosis Factor (TNF) (Roca and Ramakrishnan, 2013; Roca *et al*., 2019). mTOR-deficient macrophage death did not involve this pathway; inhibition of essential components, the kinase RIP1 and the L-type calcium channels, failed to rescue death (Figure S1I - S1K). Moreover, mTOR-deficient infected macrophages produced less mitochondrial reactive oxygen species (mROS) than wild-type ones, in contrast to macrophage death in high TNF animals, which is initiated by excess mROS (Roca and Ramakrishnan, 2013; Roca *et al*., 2019)(Figure 3A and 3B). mTOR-deficient macrophages had lower baseline mROS and a muted infection-induced increase, which only reached baseline wild-type levels (Figure 3B). mTOR regulates mitochondrial metabolism (Cunningham et al., 2007; Liu and Sabatini, 2020; Morita et al., 2013; Rambold and Pearce, 2018; Schieke et al., 2006). Our findings suggested that mTOR-dependent increases in mitochondrial metabolism in response to mycobacterial infection constitute a protective metabolic adaptation that prevents pathogenic macrophage death.

**Figure 3.**
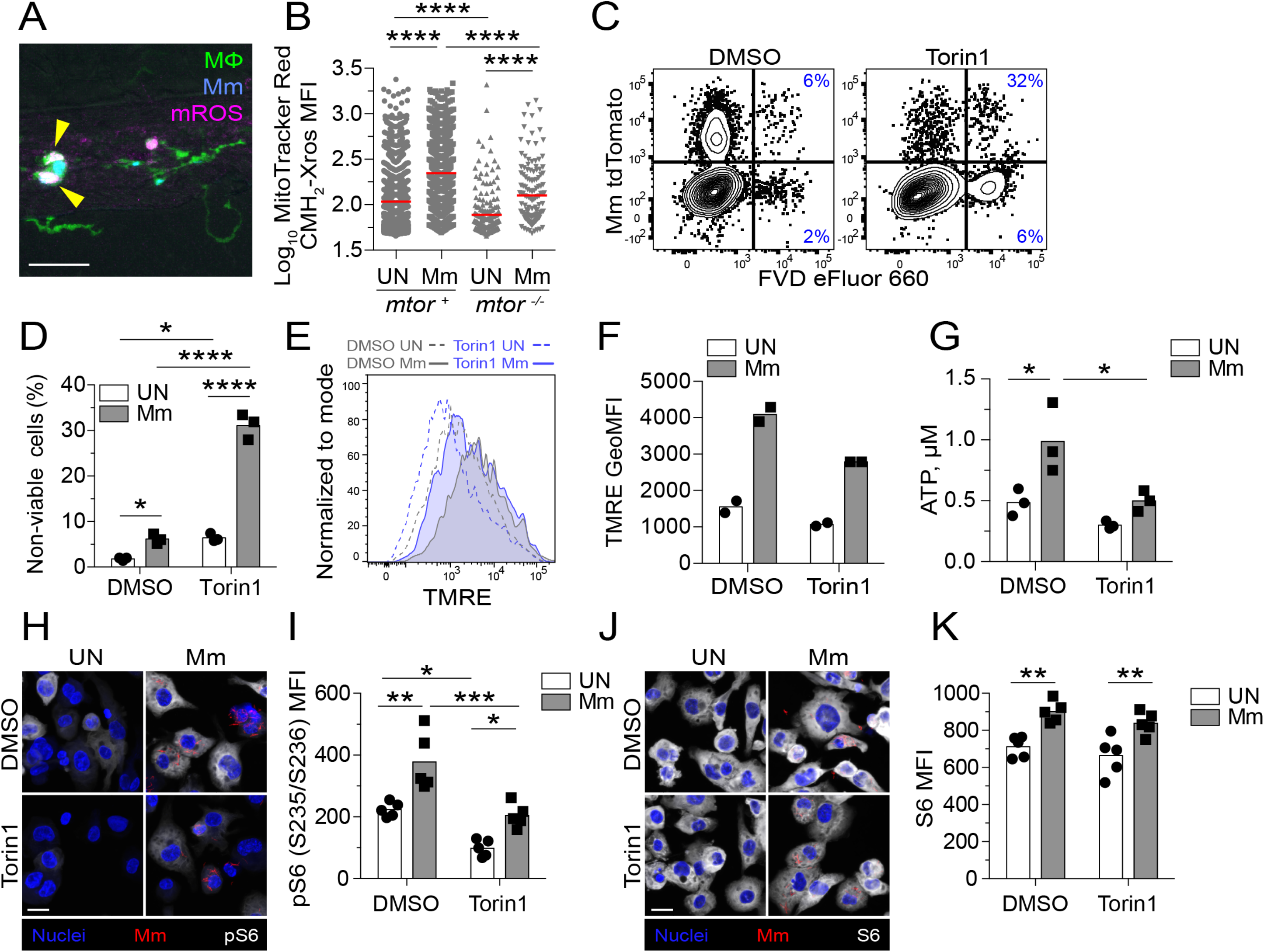
mTOR Deficiency Impairs Basal and Mycobacterium-stimulated Mitochondrial Metabolism in Macrophages. (A, B) *mtor*^*fh178/fh178*^ and mTOR-sufficient siblings expressing *Tg(mpeg1:YFP)* were infected intravenously with Mm expressing BFP2 on 2 dpf and injected intravenously with MitoTracker Red CMH_2_-Xros one day later. (A) Confocal micrograph illustrating mROS detection in an infected animal. Macrophages (green), Mm (blue), mROS (magenta), mROS-producing infected cells (arrowheads). Scale bar, 20 µm. (B) MitoTracker Red CMH_2_-Xros mean fluorescence intensity (MFI) in infected and uninfected macrophages of *mtor*^*-/-*^ animals and siblings at 1 dpi. Symbols represent individual macrophages. Horizontal lines indicate mean values. THP-1 macrophages were treated with torin1 or DMSO and infected with (C, D, G, H) tdTomato- (E, F) or mWasabi-expressing Mm at a multiplicity of infection (MOI) of 1 (C - F) or 3 (G, H). (C) Flow cytometry plots of cell viability 2 dpi. Percentages of non-viable cells (FVD eFluor 660^+^) in the infected and uninfected subpopulations are shown. (D) Quantification of non-viable cells. Symbols represent values from individual wells. Bars indicate mean values. (E) Flow cytometry histograms of TMRE fluorescence 1 dpi. (F) TMRE geometric mean fluorescence intensities (GeoMFI) 1 dpi. Symbols represent values from individual wells. Bars indicate mean values. (G) ATP concentration per well containing 10^6^ THP-1 macrophages 1 dpi. (H – K) 1 dpi THP-1 macrophage cultures infected with tdTomato-expressing Mm (MOI = 2) were treated with torin1 or DMSO for 4-hours in serum-free media. (H, J) Confocal micrographs depicting Hoechst-stained nuclei (blue), Mm (red), and (H) phospho-S6^S235/S236^ or (J) total S6 staining (white). Scale bars, 20 µm. (I and K) Mean fluorescent intensity (MFI) of (I) phospho-S6^S235/S236^ and (K) total S6 staining in uninfected and infected cells. Bars indicate group means. Symbols depict average MFI per field. Statistical analyses, (B) one-way or (D, G, I, K) two-way ANOVA with Tukey’s post-test. (A, B, H – K) Data are representative of two experiments.

To explore this further, we infected the THP-1 human macrophage cell line rendered mTOR-deficient by torin1 treatment. As in the zebrafish, mTOR deficiency increased macrophage death within one day post-infection as indicated by staining with the cell membrane-impermeant fixable viability dye eFluor 660 (Figure 3C and 3D). Moreover, Tetramethylrhodamine, ethyl ester (TMRE) staining showed that infection increased mitochondrial membrane potential in an mTOR-dependent manner (Murphy, 2009) (Figure 3E and 3F). Consistent with defective mitochondrial metabolism, mTOR-deficient macrophages had slightly lower baseline ATP production, again with a muted infection-induced increase that only reached baseline wild-type levels (Figure 3G).

We found that infection increased mTORC1 signaling as evidenced by increased phosphorylation of the ribosomal protein S6 (Battaglioni et al., 2022) (Figure 3H – 3K). Torin1-treated cells had a small increase in S6 phosphorylation, consistent with residual mTOR activity (Figure 3H and 3I). Notably, this increase only reached baseline wild-type levels, tracking with infection-induced increases in mitochondrial metabolism (mROS and TMRE) to baseline wild-type levels in mTOR-deficient conditions. Together, these findings suggest that baseline mTOR-facilitated mitochondrial metabolism is insufficient to protect macrophages from mycobacterium-induced death; adaptive infection-induced mTOR activity and corresponding increases in mitochondrial metabolism are required.

To determine if death of infected mTOR-deficient macrophages was due to mitochondrial damage, we assessed cytochrome c release by flow cytometry (Lienard et al., 2020). More mTOR-deficient infected cells had released cytochrome c (Figure 4A and 4B). In contrast, uninfected cells hardly released any cytochrome c (Figure 4B). To confirm that mitochondrial damage was the cause rather than the consequence of death, we checked if mitochondrial depolarization preceded death. Macrophages infected with blue fluorescent Mm for 32 hours were stained with TMRE, which is rapidly lost from the mitochondria upon loss of membrane potential, MitoTracker Deep Red (MTDR), a mitochondrial dye that is more resistant to changes in membrane potential, and Sytox Green, a cell membrane impermeant nucleic acid dye that labels dying cells. Widespread mitochondrial membrane depolarization (near total loss of TMRE) consistently preceded plasma membrane permeabilization and cell death (Sytox positivity) (Figure 4C and 4D and Movie S3). MTDR staining outlasted TMRE staining while preceding Sytox positivity. Moreover, its loss was often incomplete at the time of Sytox positivity, consistent with some retention of mitochondrial architecture at the commencement of the death process. Thus, infection induces widespread mitochondrial damage in mTOR-deficient macrophages causing cytochrome c release and death. Infection-induced increases in mTOR activity are adaptive, facilitating rapid increases in mitochondrial energy production, which appear to protect against this mycobacterium-mediated lethal mitochondrial damage.

**Figure 4.**
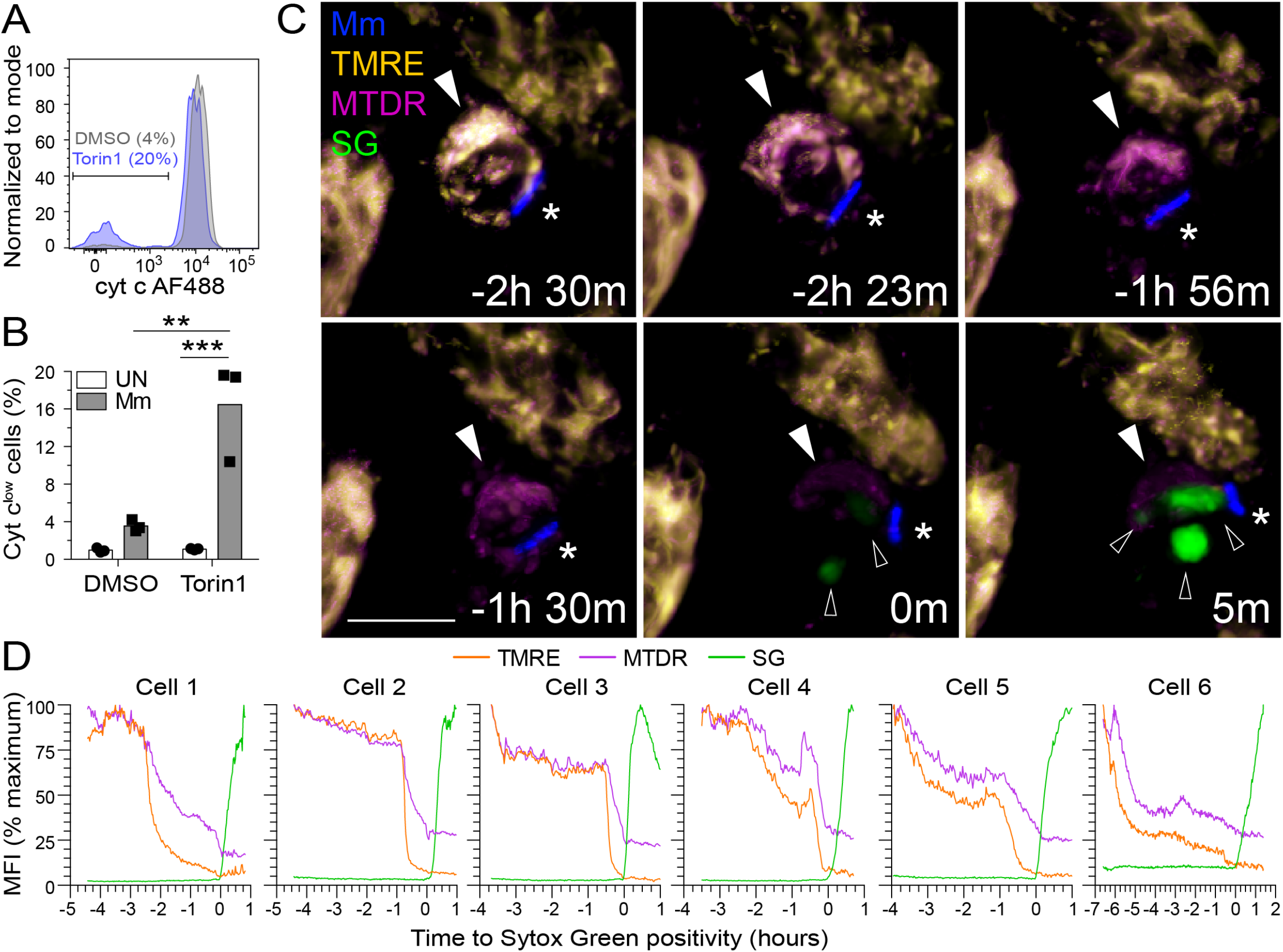
mTOR Deficiency Promotes Mycobacterium-induced, Mitochondrially-mediated Cell Death. THP-1 macrophages were infected with (A and B) tdTomato- or (C and D) BFP-expressing Mm at MOI = 3. (A) Flow cytometry histograms of cytochrome c (cyt c) fluorescence in infected viable cells (FVD eFluor 660^-^) 7 hours post-infection (7 hpi). Gate indicates cells that have released cyt c. (B) Quantification of cyt c^low^ cells 7 hpi. (C, D) Torin1-treated THP-1 macrophages were labeled with TMRE and MitoTracker Deep Red prior to imaging in the presence of Sytox Green 32 hpi. See Movie 3 and Figure S2. (C) Confocal micrographs of a dying infected macrophage (filled arrowhead) surrounded by surviving uninfected macrophages. Mm (asterisk), Sytox Green (open arrowheads). Scale bar, 10µm. (D) MFI of TMRE, MitoTracker Deep Red, and Sytox Green staining of dying infected macrophages over time. Key time-lapse frames for Cell 1 are shown in (C). Statistical analyses, (B) two-way ANOVA with Tukey’s post-test.

### mTOR Deficiency Sensitizes Macrophages to Mycobacterium-Induced Death by Impairing Glycolysis-dependent Oxidative Phosphorylation (OXPHOS)

Consistent with mTOR regulating a number of metabolic pathways that, in turn, regulate mitochondrial metabolism, mTOR inhibition of THP-1 cells caused broad reductions in glycolysis, pentose phosphate pathway and Krebs cycle metabolites (Duvel et al., 2010; Morita *et al*., 2013) (Figure S3A). Correspondingly, glycolytic and respiratory capacity were reduced in mTOR-deficient cells with lower basal and ATP-coupled respiration and spare respiratory capacity (Figure S3G − S3L). These deficits reflected compromised mitochondrial respiration and ATP production at baseline and a reduced ability to boost mitochondrial respiration in response to increased ATP demands. Moreover, mTOR deficiency blunted the small increase in mitochondrial metabolism that was apparent by 24 hours post-infection (Figure S3H and S3I). At this early stage, however, infection did not alter glucose levels, glycolytic capacity nor alter metabolite abundance in either wild-type or mTOR-deficient cells (Figure S3A − S3F and S3J − S3L). Consistent with mTOR deficiency impairing mitochondrial respiration by reducing glycolysis, selective inhibition of glycolysis with the glucose analog 2-deoxy-D-glucose (2DG), reduced Krebs cycle metabolites, mitochondrial respiration and ATP production (Figure S3A, S3F, S3M − S3O and Table S1).

Our findings were consistent with a model where mTOR-facilitated glycolytic fueling of the Krebs cycle drives the mitochondrial energy production required to protect infected macrophages from dying. If so, then inhibition of glycolysis should phenocopy mTOR deficiency, causing selective death of infected macrophages with impaired infection-induced increase in mitochondrial membrane potential. 2DG treatment induced both phenotypes in THP-1 cells, similar to torin1 treatment (Figure 5A and 5B). mTOR and glycolysis had a cytoprotective effect in Mtb infection also. To assess this, we used mc^2^ 6206, the isogenic leucine and pantothenate auxotrophic mutant of the virulent H37Rv Mtb strain, a biosafety 2 level pathogen that elicits similar inflammatory responses and triggers diverse cell death programs (Beckwith *et al*., 2020; Mouton et al., 2019; Roca *et al*., 2022; Roca *et al*., 2019; Sampson et al., 2004; Sampson et al., 2011). Mtb infection caused increased death of both mTOR-deficient and glycolysis-deficient THP-1 cells (Figure 5C). In the zebrafish, 2DG treatment depleted infected macrophages selectively and increased bacterial cording (Figure 5D - 5G and Movie S4). Thus, mTOR exerts its cytoprotective effect by supporting glycolysis both in Mm zebrafish infection and Mm- and Mtb-infected human macrophages.

**Figure 5.**
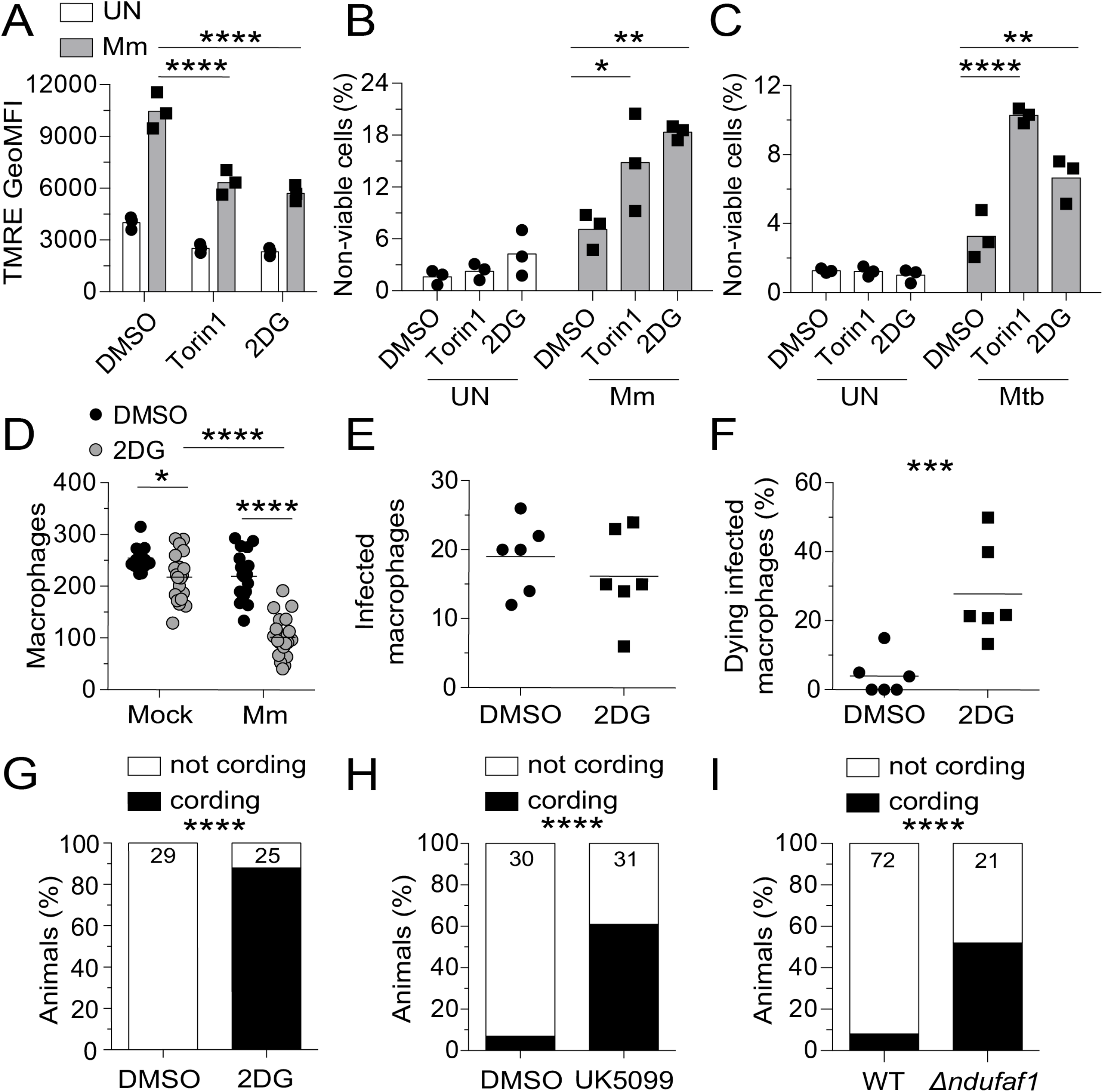
Glycolysis Inhibition Impairs Mitochondrial Metabolism and Sensitizes Infected Macrophages to Mycobacterium-induced Cytotoxicity. THP-1 macrophages treated with torin1 (400 nM), 2-deoxy-D-glucose (2DG, 5 mM) or DMSO were infected with Mm expressing (A) BFP2 or (B) tdTomato or (C) Mtb expressing tdTomato (MOI = 1). (A) TMRE GeoMFI 1 dpi. (B, C) Percentage of non-viable cells (FVD eFluor 660^+^) 1 dpi. (D – I) Zebrafish were infected with ∼150 fluorescent Mm via the caudal vein. (D) 5 dpi macrophage numbers in the body of mock- or Mm*-*infected *Tg(mpeg1:YFP)* zebrafish fish treated with 50 mM 2DG or 0.5% DMSO. (E, F) 6-hr time-lapse confocal microscopy of *Tg(mpeg1:YFP)* 3 dpi. (E) Absolute numbers of infected macrophages per field. (F) Percentage of dying infected macrophages per field. See Movie S3. (G) Cording in wild-type animals treated with 2DG or DMSO 5 dpi. (H) Cording in wildtype animals treated with UK5099 (10 µM) or 0.5% DMSO 5 dpi. (I) Cording in *ndufaf1* G0 crispants and wild-type (WT) siblings 5 dpi. Symbols represent values from individual (A-C, K) wells or (D - F) animals. (A - C) Bars and (D - F) horizontal lines indicate mean values. (G – I) Numbers within columns indicate animals per group. Statistical analyses, one-way ANOVA with (A - C) Sidak or (D) Tukey post-tests, (E, F) unpaired Student’s *t* test, or (G – I) Fisher’s exact test. (E, F) Time-lapse data were pooled from two independent experiments. Data are representative of (A, G, H) two independent experiments. See also Figure S2.

Since glycolysis contributes to mitochondrial ATP production by supplying pyruvate to the Krebs cycle (Ryan and O’Neill, 2020), intercepting this step should also produce bacterial cording. Inhibition of the mitochondrial pyruvate carrier with UK5099 similarly increased bacterial cording (Halestrap, 1975)(Figure 5H). mTOR-facilitated glycolysis also feeds the pentose phosphate pathway (Patra and Hay, 2014)(Figure S2A). Zebrafish mutants deficient in glucose-6-phosphate dehydrogenase (G6PD), the rate-limiting enzyme of the pentose phosphate pathway, did not have increased bacterial cording, ruling out the contribution of this pathway to mTOR-mediated resistance (Figure S3P and S3Q). Thus, mTOR deficiency sensitizes macrophages to Mm-induced mitochondrial damage and death by impairing glycolysis-dependent OXPHOS and thereby mitochondrial energy production. Consistent with this, genetic disruption of the electron transport chain complex 1 by targeting the assembly factor NDUFAF1 increased bacterial cording (Formosa et al., 2018)(Figure 5I).

### mTOR-dependent Glycolysis and OXPHOS Enable Macrophages to Resist Mycobacterium ESAT-6-mediated Death

The mTOR mutant had unmasked the potential of pathogenic mycobacteria to cause lethal mitochondrial damage in infected macrophages. We hypothesized that a specific mycobacterial determinant caused this damage. Our prime candidate was the ESX-1 secretion system because (1) it accelerates mycobacterium-induced macrophage death in wild-type (mTOR-sufficient) conditions, including in the zebrafish (Conrad et al., 2017; Davis and Ramakrishnan, 2009; Groschel et al., 2016; Simeone et al., 2012; Simeone et al., 2021; Volkman et al., 2004), and (2) it mediates mitochondrial damage in infected macrophages (Lienard *et al*., 2020; Wiens and Ernst, 2016). Whereas in wild-type macrophages, ESX-1-dependent death requires high intramacrophage bacterial burdens (Davis and Ramakrishnan, 2009; Volkman *et al*., 2004), we hypothesized that mTOR-deficiency sensitizes macrophages to ESX-1-mediated mitochondrial damage, causing death at low bacterial burdens. Consistent with our hypothesis, mTOR-deficient THP-1 macrophages infected with ESX-1-deficient Mm did not have increased cytochrome c release nor increased death (Figure 6A and 6B and Movie S5). In mTOR-deficient zebrafish infected with ESX-1-deficient Mm, infected macrophages did not die (Figure 6C − 6E and Movie S6). ESX-1-deficient infection also did not cause macrophage death in animals deficient in glycolysis or OXPHOS (Figure 6F − 6H). Thus, mycobacterium-induced, mTOR-dependent increase in mitochondrial metabolism specifically counters ESX-1-dependent mitochondrial damage and cell death.

**Figure 6.**
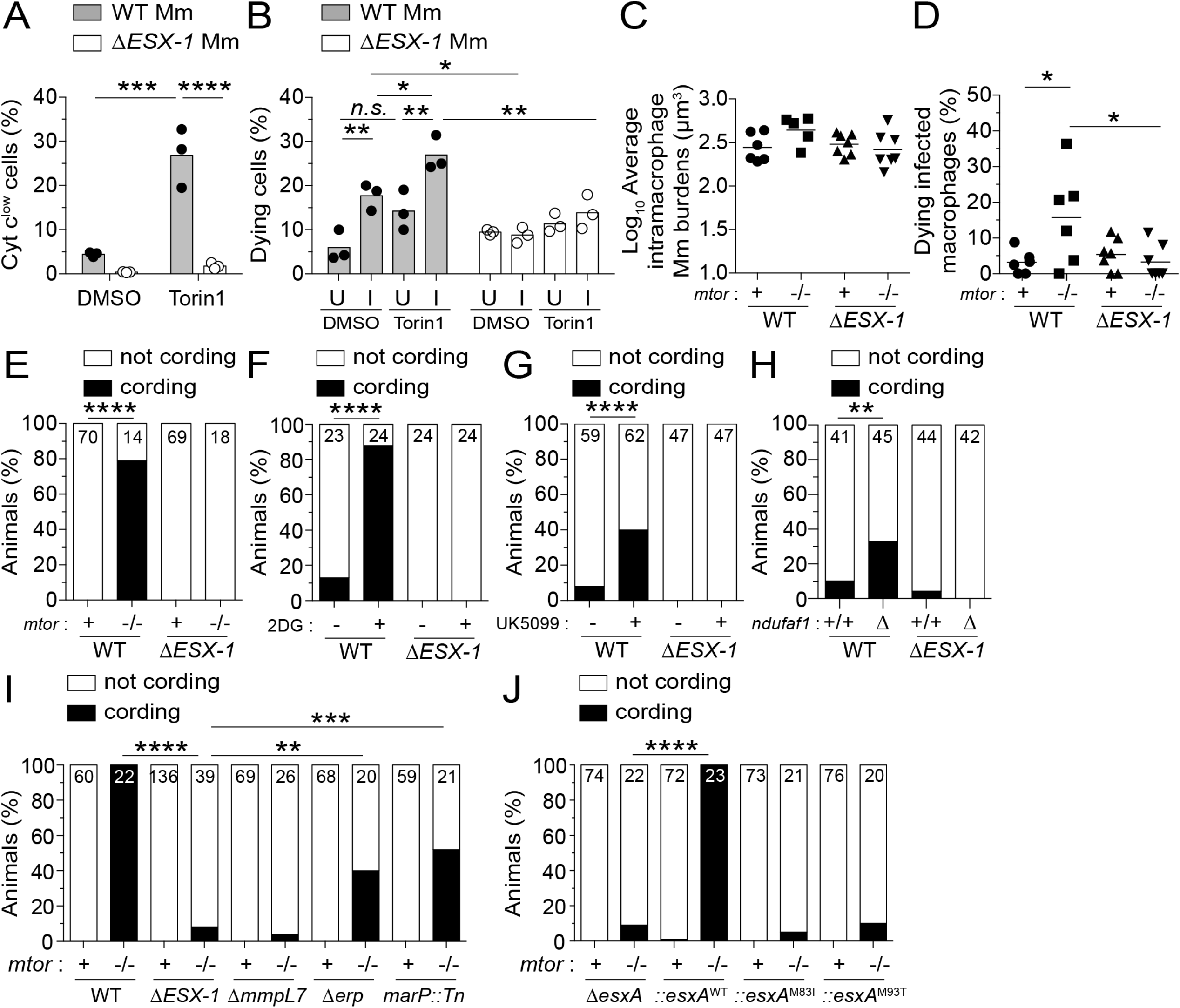
Deficiencies in mTOR, Glycolysis, and OXPHOS Sensitize Macrophages to Mycobacterial ESAT-6-dependent Cytotoxicity. (A) Cytochrome c release 7 hpi in THP-1 macrophages infected with BFP2-expressing WT or Δ*ESX-1* Mm at MOI = 3. (B) Percentage of dying cells (Sytox Green^+^) during 4-hr time-lapse at 1 dpi with tdTomato-expressing WT or Δ*ESX-1* Mm at MOI = 1. Values from uninfected (U) and infected (I) cells from the same fields are shown. See Movie S4. (C – J) Zebrafish were infected with dose-matched inocula of tdTomato-expressing Mm of the indicated strains via the caudal vein. (C) Intramacrophage Mm burdens at the beginning of 6-hr time-lapse confocal microscopy of *mtor*^*fh178/fh178*^ and *mtor-*sufficient siblings expressing *Tg(mpeg1:YFP)* 2 dpi. See Movie S5. (D) Percentage of dying infected macrophages in same experiment shown in (C). See Movie S5. (E) Cording in *mtor*^*fh178/fh178*^ animals and *mtor-*sufficient siblings 4 dpi. (F, G) C in wild-type zebrafish treated with 2DG, UK5099, or DMSO 5 dpi. (H) Cording in *ndufaf1* G0 crispants and WT siblings 5 dpi. (I) Cording in *mtor*^*sa16755/sa16755*^ animals and *mtor-*sufficient siblings 4 dpi. See also Figure S3. (J) Cording in *mtor*^*fh178/fh178*^ animals and *mtor-*sufficient siblings 4 dpi with *ΔesxA* Mm complemented with wild-type or point mutant Mtb *esxA*. Symbols represent values from individual (A) wells, (B) imaging fields, or (C) animals. (A, B) Bars and (C, D) horizontal lines indicate mean values. (E – J) Numbers within columns indicate animals per group. Statistical analyses, (A – D) one-way ANOVA with Sidak’s post-test or (E – J) Fisher’s exact test. (B, E, H) Data are representative of two experiments. Zebrafish time-lapse data were pooled from four experiments.

ESX-1-mediated damage of *Mycobacterium*-containing phagosomes is integral to its role in virulence (Lienard *et al*., 2020; Simeone *et al*., 2012; Simeone *et al*., 2021). This process is facilitated by the mycobacterial cell surface lipid phthiocerol dimycoceroserate (PDIM); Mm mutants in the PDIM transporter MmpL7 are also impaired in phagosomal damage as indicated by reduced galectin-8 recruitment (Augenstreich et al., 2017; Lerner et al., 2018; Osman et al., 2020; Quigley et al., 2017; Simeone *et al*., 2021) (Cambier et al., 2014b)(Figure S4A and S4B). To determine if ESX-1-mediated phagosomal damage was integral to its mitotoxicity in mTOR-deficient macrophages, we tested the Mm mmpL7 mutant. mmpL7 mutant Mm infection did not kill mTOR-deficient macrophages, as evidenced by the lack of cording in the animals (Figure 6I). In contrast, Mm mutants in the virulence determinants Erp and MarP, which do not mediate phagosomal damage and promote intramacrophage growth through distinct mechanisms from ESX-1, did accelerate macrophage death in mTOR-deficient animals (Berthet et al., 1998; Cosma et al., 2006; Levitte et al., 2016; Vandal et al., 2008)(Figure S4A and S4B and Figure 6I). Thus, ESX-1-dependent phagosomal permeabilization is a pre-requisite for its cytotoxicity to mTOR-deficient macrophages.

ESX-1’s membranolytic activity had been ascribed to its major secreted substrate ESAT-6 (6 kDa early secretory antigenic target); however, pinning down its role versus those of other ESX-1 substrates has been complicated by their co-dependency for secretion, as deletion of ESAT-6 causes loss of other ESX-1 substrates (Bao et al., 2021; Champion et al., 2014; Fortune et al., 2005). Recent work has pinpointed a specific role for ESAT-6 in phagosomal damage and virulence by identifying point mutations in ESAT-6 that allow substantial levels of secretion of ESAT-6 and other ESX-1 substrates yet cause loss of phagosomal membrane damage and/or virulence (Brodin et al., 2005; Osman et al., 2022; Zhang et al., 2016). To test the specific role of ESAT-6-induced phagosomal damage in mediating cell death in mTOR deficiency, we used *esxA* (ESAT-6) mutant Mm expressing either of two such ESAT-6 C-terminal point mutations, M83I and M93T (Brodin *et al*., 2005; Osman *et al*., 2022). Mm-ESAT-6^M83I^ and Mm-ESAT-6^M93T^ infections did not kill macrophages in mTOR-deficient zebrafish (Figure 6J). Thus, ESAT-6 causes the phagosomal damage required for the cell death induced by mTOR deficiency.

### mTOR Enables Infected Macrophages to Specifically Resist ESAT-6-mediated Mitochondrial Damage

mTOR-deficiency might simply sensitize phagosomes to ESAT-6-mediated damage. However, mTOR-deficient and wild-type macrophages had similar ESAT-6-dependent increases in phagosomal damage, showing that this was not the case and instead suggesting that mTOR-deficient macrophages were sensitized to ESAT-6-mediated mitochondrial damage (Figure 7A, upper panels and 7B, black symbols).

**Figure 7.**
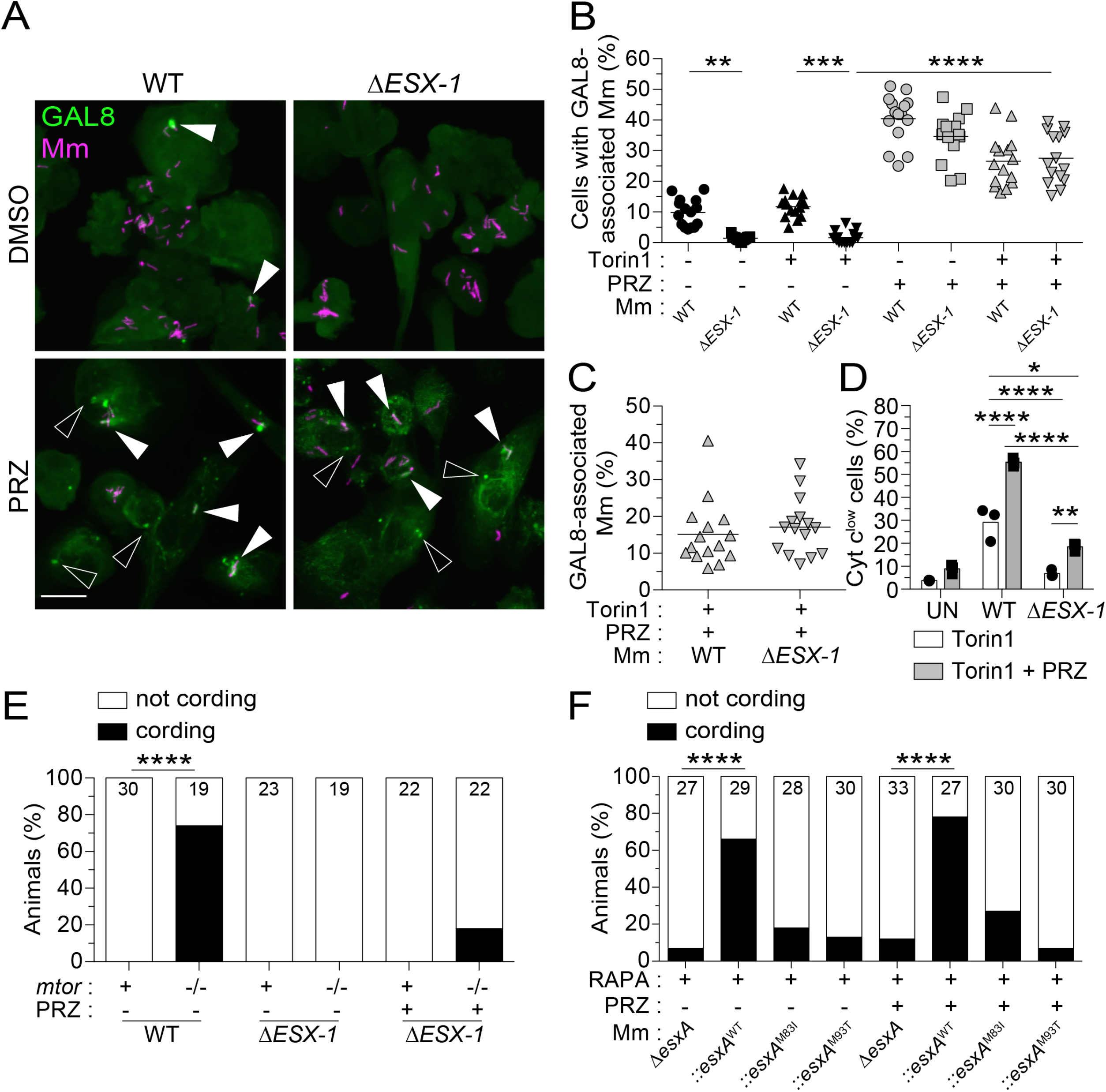
ESAT-6 Mediates Mitochondrial Damage in mTOR-deficient Macrophages Downstream of Its Involvement in Phagosomal Permeabilization. (A - E) Torin1- and DMSO-treated THP-1 macrophages were infected with tdTomato-expressing WT or Δ*ESX-1* Mm at MOI = 3 and treated with prazosin (PRZ, 20 µM) for 7 hrs. See also Figure S4. (A) Confocal micrographs of Galectin-8 (GAL8) immunofluorescence (green) and Mm fluorescence (magenta) in THP-1 macrophages 7 hpi. GAL8 foci associated with Mm (filled arrowheads) or not associated with Mm (open arrowheads) are shown. Scale bar, 20 µm. (B) Percentage of macrophages with GAL8-associated Mm foci. (C) Percentage of Mm volume associated with GAL8 foci 7 hpi. (D) Percentage of cells that have released cytochrome c 7 hpi. (E) *mtor*^*sa16755/sa16755*^ fish and *mtor-*sufficient siblings were infected with ∼90 fluorescent Mm via the hindbrain ventricle on 2 dpf. On 1 and 2 dpi, animals were injected with ∼3 nL of 300 µM PRZ or 1% DMSO into the hbv. Graph indicates the percentage of animals with cording 3 dpi. (F) Wild-type fish treated with 400 nM rapamycin were infected with ∼180 tdTomato-expressing *ΔesxA* Mm complemented with wild-type or point mutant Mtb *esxA* via the hbv on 2 dpf. Animals were injected with PRZ or DMSO as indicated on (E). Graph indicates the percentage of animals with cording 3 dpi. Symbols represent values from individual (B, C) imaging fields or (D) individual wells. (B, C) Horizontal lines and (D) bars indicate mean values. (E, F) Numbers within columns indicate animals per group. Statistical analyses, (B, D) one-way ANOVA with Sidak’s post-test or (E, F) Fisher’s exact test.

ESAT-6 might mediate mitochondrial damage in mTOR-deficient macrophages indirectly or directly. In the indirect case, ESAT-6 would only be required to permeabilize phagosomes, enabling host lysosomal factors or other mycobacterial determinant(s) to access and damage mitochondria. In the direct case, ESAT-6 would also be required after the phagosome has been permeabilized. To distinguish between the two, we treated THP-1 cells with prazosin, a drug that permeabilizes endo-lysosomal compartments in myeloid cells (Kozik et al., 2020). By 7 hours of treatment, prazosin had caused extensive endo-lysosomal damage in uninfected control and mTOR-deficient macrophages (Figure S4C). In infected macrophages, prazosin caused similar increases in damaged wild-type and ESX-1-deficient mycobacterial phagosomes (Figure 7A and 7B). Moreover, the proportion of wild-type and ESX-1-mutant mycobacteria associated with damaged phagosomes was similar, showing that the prazosin-induced phagosomal permeabilization could equalize access of wild-type and ESX-1-deficient mycobacteria to the mitochondria (Figure 7C). Consistent with increasing bacterial exposure to mitochondria, prazosin increased cytochrome c release from wild-type-infected macrophages (Figure 7D). Prazosin-treated ESX-1 mutant-infected macrophages had a much smaller increase in cytochrome c release, and significantly lower than in untreated wild-type macrophages (Figure 7D). The small increase in cytochrome c release in ESX-1-mutant infection suggests that additional mechanisms may also play a role, for instance PDIM. To corroborate these conclusions in vivo, we assessed bacterial cording in zebrafish with and without prazosin treatment. As before, mTOR deficiency caused cording of wild-type but not ESX-1-deficient bacteria (Figure 7E). Prazosin treatment of mTOR mutants increased ESX-1-mutant cording only slightly, and significantly less than wild-type bacteria without prazosin (Figure 7E).

Next, we tested if ESAT-6 was responsible for the mitochondrial damage by assessing bacterial cording of ESAT-6-mutant bacteria in zebrafish with and without prazosin treatment. As with ESX-1 mutant infections, prazosin treatment did not restore bacterial cording of ESAT-6 mutants (Figure 7F). These findings implicate ESAT-6 in mediating mitochondrial damage after first damaging the phagosome to enable access. ESAT-6 possesses both phagosomal and mitochondrial damaging activity and mTOR protects specifically against the latter. Our findings highlight the strength of mTOR as a “counter-virulence” factor against ESAT-6 and indicate that a small amount of ESAT-6 is sufficient to induce cytotoxicity in mTOR-deficient macrophages. mTOR-deficient macrophages tolerate infection with mycobacteria so long as they lack functional ESAT-6.

## DISCUSSION

Consistent with mTOR’s critical role in the development, homeostasis and function of myeloid cells and T cells (Lachmandas et al., 2016; Powell et al., 2012; Sinclair et al., 2017; Weichhart et al., 2015), therapeutic mTOR blockade in organ transplantation and cancer has been associated with increased risk of infections, including TB (Fijalkowska-Morawska et al., 2011; Garcia and Wu, 2016; Jeon et al., 2017; Ruiz-Camps and Aguilar-Company, 2021; Tsai et al., 2007). This susceptibility has generally been ascribed to compromised adaptive immunity (Weichhart *et al*., 2015). We show here that mTOR deficiency results in profound, innate susceptibility to mycobacteria resulting from the rapid death of infected macrophages. This death results from catastrophic mitochondrial damage caused by mycobacterial ESAT-6. Thus, mTOR-facilitated mitochondrial metabolism represents a formidable armor against a potent mycobacterial mitotoxin. This study adds to the appreciation that while intracellular microbes can exploit host cell metabolism for growth and pathogenesis, the cells’ metabolic capabilities can also avert microbial attack (Pernas, 2021).

Our findings provide insight into the adaptive metabolic changes that occur upon mycobacterial infection. Stimulation of cultured macrophages with lipopolysaccharide (LPS), a major Gram-negative bacterial virulence determinant, induces a metabolic switch from mitochondrial OXPHOS to glycolysis under normoxic conditions (O’Neill and Pearce, 2016; Weichhart *et al*., 2015). This switch, which occurs through mTORC1 activation, enables macrophages to elaborate antimicrobial responses that depend on cataplerosis of Krebs cycle intermediates (O’Neill and Pearce, 2016). Whether Mtb infection induces this glycolytic switch is unclear; studies variably find that it boosts or represses glycolysis (Braverman et al., 2016; Cumming et al., 2018; Gleeson et al., 2016; Huang et al., 2018; Lachmandas *et al*., 2016; Olson et al., 2021; Shi et al., 2015). We find here that early in infection, Mm infection of cultured human macrophages induces OXPHOS without altering glycolysis. This is consistent with our findings in the zebrafish where Mm- and Mtb-infected macrophages exhibit small increases in mitochondrial respiration early on (Roca *et al*., 2022). While adaptive glycolytic shifts may occur later in infection, mTORC1’s role in early resistance to mycobacteria stems not from a glycolytic shift but from the boost in OXPHOS from its stimulation of glycolysis (Weichhart *et al*., 2015). This is highlighted by our finding that directly inhibiting OXPHOS confers susceptibility.

How does ESAT-6 damage mitochondria? Mm and Mtb phagosomes frequently fuse to lysosomes, and contacts between lysosomes and mitochondria are proposed to facilitate metabolic exchanges between these organelles (Armstrong and Hart, 1971; Barker et al., 1997; Clemens and Horwitz, 1995; Harris et al., 2008; Levitte *et al*., 2016; Wong et al., 2018). The idea that mycobacterial phagolysosomes influence mitochondrial function is reinforced by the finding that mitochondria aggregate around these structures and undergo morphological changes (Jamwal et al., 2013; Mohareer et al., 2020). Given that direct contact is required for ESAT-6 permeabilization of host membranes (Conrad *et al*., 2017), ESAT-6 might specifically damage membranes of mitochondria that are in direct contact with permeabilized phagosomes. Indeed, ESX-1 is reported to cause some mitochondrial damage in mTOR-sufficient cells as evidenced by reductions in mitochondrial mass, membrane potential loss, and release of cytochrome c and mitochondrial DNA (Chen et al., 2008; Fine-Coulson et al., 2015; Lienard *et al*., 2020; Pajuelo et al., 2018; Wiens and Ernst, 2016). The finding that in mTOR deficiency, infection with very few ESX-1/ESAT-6-expressing mycobacteria, likely confined to a single phagosome/phagolysosome, causes global loss of mitochondrial membrane potential suggests that there is rapid propagation of the initial localized damage. The lack of rapid repair or replacement of damaged mitochondria in the absence of mTOR-regulated biosynthetic processes may account for this (Cunningham *et al*., 2007; Liu and Sabatini, 2020; Morita *et al*., 2013; Morita et al., 2017; Rambold and Pearce, 2018; Schieke *et al*., 2006). Alternatively, reduced mitochondrial membrane potential has been shown to cause structural changes in the mitochondria – matrix condensation and unfolding of cristae – that facilitate cytochrome c release upon exposure to outer membrane disrupters like BAX family proteins (Gottlieb et al., 2003). Similar changes from mTOR deficiency could cause global mitochondrial catastrophe when subjected to ESAT-6’s mitotoxic effects. As in that report, we too find that disruption of respiration at complex I phenocopies the ESAT-6-mediated death produced by mTOR deficiency.

ESX-1 increases macrophage death even in wild-type (mTOR competent) macrophages (Groschel *et al*., 2016; Ramakrishnan, 2012) (Augenstreich *et al*., 2017; Beckwith *et al*., 2020; Behar et al., 2010; Davis and Ramakrishnan, 2009; Srinivasan et al., 2014), with ESAT-6-mediated phagosomal damage being a pre-requisite step (Bao *et al*., 2021; Osman *et al*., 2022; Simeone *et al*., 2021; Srinivasan *et al*., 2014; Zhang *et al*., 2016). We show here that ESAT-6 induces phagosomal damage irrespective of whether the macrophage is mTOR-sufficient or -deficient. mTOR’s specific role is in resistance against ESAT-6-mediated catastrophic mitochondrial damage that rapidly kills the cell. We do not know if ESAT-6 is sufficient for mitochondrial damage in mTOR deficiency. ESX-1 is responsible for plasma membrane damage in mTOR-sufficient cells and there too it is not clear exactly how damage is mediated — studies have variably implicated ESAT-6, other ESX-1 substrates, non-ESX-1 mycobacterial products, and excess lipoxins (Beckwith *et al*., 2020; Divangahi et al., 2009; Pagan and Ramakrishnan, 2018; Pajuelo *et al*., 2018). For the mitochondrial damage in mTOR deficiency, PDIM may facilitate ESAT-6’s role, in addition to facilitating ESAT-6-mediated phagosomal damage.

TB is notorious for killing far more people over the millennia than any other infectious agent (Paulson, 2013). Yet, our recent epidemiological analyses show that most individuals clear Mtb infection through a combination of innate and adaptive immunity (Behr *et al*., 2018; 2019). Our work finds that tapping into mTOR-regulated homeostatic metabolic pathways constitutes a major host defense strategy. Curiously, these pathways protect against a specific mycobacterial virulence determinant, identifying mTOR as a “counter-virulence” factor against ESAT-6, where mTOR averts its catastrophic mitotoxicity to buy the host time to call in other “classical” immune defenses that can clear infection much of the time. In the minority of individuals, ESAT-6 and other mycobacterial virulence factors win out, allowing the evolutionary survival of the pathogen as well (Cambier *et al*., 2014a). Indeed, the modes of cell death that ESAT-6 produces in wild-type hosts through phagosomal or plasma membrane damage likely represent “workarounds” in mycobacterium’s pathogenic strategy (Beckwith *et al*., 2020; Chen *et al*., 2008; Divangahi *et al*., 2009; Pagan and Ramakrishnan, 2018; Pajuelo *et al*., 2018; Simeone *et al*., 2021; Zhang et al., 2021).

Therapies targeting mTOR are being explored for a number of conditions, including aging. Genetic and pharmacological mTOR inhibition can increase lifespan in yeast, worms, flies, and mice (Papadopoli et al., 2019; Saxton and Sabatini, 2017). Pilot studies of a short course of pharmacological mTOR inhibition in older human volunteers report increased responses to influenza vaccines (suggesting decreased immune senescence) and a reduction in self-reported viral respiratory infections (Mannick et al., 2014; Mannick et al., 2018). Similarly, in lung TB patients receiving adjunctive mTOR inhibition therapy together with appropriate antimicrobial treatment had possible, transient improvement in lung function (Wallis et al., 2021). In a mouse model of severe TB, mTOR inhibition therapy induced host-beneficial or -detrimental effects depending on the treatment regimen; mTOR inhibition therapy reduced lung immunopathology in established infections when given in conjunction with an antimicrobial drug, but exacerbated lung damage and morbidity when administered alone in the early infection (Bhatt et al., 2021). Our finding that mTOR inhibitors dramatically increase susceptibility to pathogenic mycobacteria warrants caution in their use as anti-aging or immune boosting therapies in the many areas of the world with a high burden of TB.

### Limitations of the Study

How ESAT-6 causes the mitochondrial damage which ultimately triggers cell death in mTOR deficiency remains to be determined. ESAT-6 may directly permeabilize the outer membrane of mitochondria in close proximity to permeabilized phagosomes or induce mitochondrial damage by cooperating with host or mycobacterial molecules with intrinsic mitotoxic activity. Furthermore, how reductions in mitochondrial membrane potential specifically increase sensitivity to ESAT-6-mediated mitotoxicity remains unclear. Identifying the molecular players in the cell death cascade and the alterations in mitochondrial structure and function in mTOR deficiency, and specifically in response to ESAT-6-mediated damage, should help to clarify these processes.

## ACKNOWLEDGEMENTS

We thank M. Behr, P. Edelstein, M. Murphy and F. Roca for discussions; M. Eisenberg-Bord and N. Yamaguchi for manuscript review; K. Takaki for parabiosis instruction; C. Hale for providing access to Seahorse XFp analyzer and advice on experimental design; J. Ray for assistance mapping *mtor*^*fh178*^; R. Berg, L. Hernández, S. Levitte, and J. Zimmermann for assistance with early characterization of *mtor*^*fh178*^; H. Boyd, B. Brockley, C. Dooley, S. McGuinan, R. Santhakumar, I. Smallwood, and R. White for technical and logistical assistance with genetic screens; J. Cameron, N. Goodwin, R. Foster, and the University of Cambridge aquatics facility staff for zebrafish husbandry; the LMB’s flow cytometry core facility and the NIHR Cambridge BRC Cell Phenotyping Hub for equipment access, and the LMB’s media service for preparation of bacterial and tissue culture reagents. This work was supported by the National Institutes of Health (T32-AI055396, A.J.P. and MERIT award R37-AI054503 and Director’s Pioneer Award DP1MH099901, L.R.), a Wellcome Trust Principal Research Fellowship (223103/Z/21/Z; L.R.), the Max Planck Society (E.L.P.), and the Liebnietz Prize (E.L.P).

## AUTHOR CONTRIBUTIONS

A.J.P., C.B.M., D.M.T., and L.R. conceived the project. A.J.P., L.J.L., J.E.-H., C.B.M., D.M.T., E.M.B.-N., E.L.P. and L.R. designed experiments and analyzed data. A.J.P. and L.J.L. performed zebrafish experiments. D.M.T. isolated and mapped *mtor*^*fh178*^. A.J.P. conducted cell culture experiments. J.E.-H. performed and analyzed LC/MS. C.B.M. and E.M.B.-N. provided chemically-mutagenized zebrafish lines. E.L.P. suggested and supervised metabolomics experiments and advised on the experimental design and interpretation of metabolic experiments. A.J.P. and L.R. wrote the manuscript and got input from the other authors.

## SUPPLEMENTARY FIGURES, TABLES, AND MOVIES

**Figure S1. mTOR Deficiency Impairs Hematopoiesis in Zebrafish**.

(A) Overlaid widefield fluorescence and brightfield micrographs of an *mtor*^*fh178/fh178*^ animal and wild-type sibling expressing the neutrophil-specific fluorescent reporter *Tg(lysC:EGFP)* 6 dpf (Hall et al., 2007).

(B) Numbers of neutrophils in the caudal hematopoietic tissue (CHT) of animals from *mtor*^*fh178/+*^; *Tg(lysC:EGFP)* incross 2 and 6 dpf.

(C – F) Zebrafish embryos were manually dechorionated and treated with 400nM rapamycin or 0.5% DMSO on 1 dpf to block primitive and intermediate waves of hematopoiesis (Clements and Traver, 2013).

(C) Confocal micrographs of the CHT of *Tg(cd41:GFP)* zebrafish at 2 dpf. Hematopoietic stem cells (HSCs, open arrowheads) and thrombocytes, nucleated counterparts of platelets in non-mammalian vertebrates, (filled arrowheads) were identified by low vs. high level expression of *cd41:GFP*, respectively (Lin et al., 2005; Ma et al., 2011).

(D) Numbers of HSCs in the CHT of 2 dpf animals.

(E) Numbers of thrombocytes in the CHT of 2 dpf animals.

(F) Numbers of macrophages in the midbrain and CHT of *Tg(mpeg1:YFP)* zebrafish 2 dpf.

Scale bars, (A) 300 µm and (C) 100 µm. (B, D – F) Symbols represent individual animals. Horizontal lines indicate means. Statistical analyses, (B) two-way ANOVA with Tukey’s post-test and (D – F) unpaired Student’s *t* test.

**Figure S2. Inhibition of Autophagic Cell Death, Mitochondrial Apoptosis, or TNF-associated Necrosis Does Not Prevent Mycobacterium-induced Macrophage Death in mTOR-deficient Animals, Related to Figures 2 and 3**.

(A) Confocal micrographs of LC3 clustering in neuromasts, clusters of mechanosensory cells of the fish lateral line, from *atg12*^*sa42684*^ incross fish expressing *Tg(CMV:lc3b-GFP)* 5 dpf. GFP fluorescence (top) and surface-rendered puncta (bottom) are shown. Scale bar, 10 µm.

(B) Number of LC3 puncta per neuromast.

(C) Cording in rapamycin- or DMSO-treated *atg12*^*sa42684*^ incross fish 5 dpi.

(D) Confocal micrographs of acridine orange (AO) staining (green) and surface-rendered puncta (magenta) in the midbrain of *casp9sa* incross fish 3 dpf. Scale bar, 50 µm.

(E) Number of AO puncta in the midbrain.

(F) Cording in rapamycin- or DMSO-treated *casp9*^*sa11164*^ incross fish 5 dpi.

(G) Cording in rapamycin- and DMSO-treated *pycard*^*w216/w216*^ (Asc-deficient) animals and siblings 4 dpi.

(H) Cording in rapamycin- and DMSO-treated *sting1*^*sa35634*^ (Sting-deficient) animals and WT siblings 5 dpi.

(I) Necrosis pathway induced by mycobacterial infection plus excess TNF and pharmacological interventions tested.

(J, K) Mycobacterial cording in *mtor*^*fh178/+*^ incross fish treated with (H) necrostatin-1, (I) nifedipine (5 µM), diltiazem (10 µM), or 0.5% DMSO 4 dpi.

Symbols represent individual (B) neuromasts or (E) animals. (B, E) Horizontal lines indicate mean values. (C, F, G, H, J, K) Numbers within columns indicate animals per group. (B, E) one-way ANOVA with Tukey’s post-test.

**Figure S3. mTOR Inhibition Impairs Glycolysis and Mitochondrial Metabolism, Related to Figure 5**.

(A) Metabolite profiles of uninfected and Mm-infected THP-1 macrophages 1 dpi (MOI = 1). Cell were treated with torin1 (400nM), 2DG (5mM), or 0.5% DMSO for 1.5 days prior to harvest. Heat map scale indicates relative log_2_ expression levels. See also Table S1.

(B – F) Volcano plots of differences in metabolite abundances induced by the indicated treatments. Dashed lines indicate fold-change and p value cut-offs.

(G) Diagram of Mitochondrial Oxygen Consumption Rate (OCR) assay.

(H) OCR kinetics of torin1 or DMSO-treated THP-1 macrophages 1 dpi (Mm, MOI = 4).

(I) Modular analysis of mitochondrial OCR.

(J) Diagram of Glycolytic Proton Efflux Rate (PER) assay.

(K) PER kinetics of torin1 or DMSO-treated THP-1 macrophages 1 dpi (Mm, MOI = 4).

(L) Basal and compensatory glycolytic PER.

(M) OCR kinetics of uninfected THP-1 macrophages treated with 2DG or DMSO for 1.5 days.

(N) Modular analysis of mitochondrial OCR.

(O) Relative ATP levels in THP-1 macrophage cultures 1.5 days after treatment.

(P) Glucose-6 phosphate dehydrogenase (G6PD) activity in 5 dpf animals from *g6pd*^*sa24272/+*^ incross.

(Q) Cording in animals from *g6pd*^*sa24272/+*^ incross 5 dpi.

Symbols represent (B – F) individual metabolites, (H, K, M) mean values, (O) individual wells, or (P) individual animals. (I, L, N – P) Bars indicate mean values. (H, I, K, L, M, N) Error bars depict standard deviation. (G, J) Arrows indicate the time of compound injection. Abbreviations: rotenone plus antimycin A (Rot + AA), 2DG, oligomycin (Oligo), carbonyl cyanide 4-(trifluoromethoxy)phenylhydrazone (FCCP), compensatory glycolysis (Comp), spare respiratory capacity (SRC). Statistical significance, (B – F) unpaired Students’ *t* test, one-way ANOVA with (I, L, N) Sidak or (P) Tukey’s post-tests.

**Figure S4. Damage of Phagosomal/Lysosomal Compartments by ESX-1-competent Mycobacteria and the Drug Prazosin, Related to Figures 6 and 7**.

(A) Confocal micrographs of Galectin-8 (GAL8) immunofluorescence (green) and Mm fluorescence (magenta) in THP-1 macrophages infected with the indicated Mm strains (MOI = 1) 1 dpi. The *ΔmmpL7* Mm strain is defective in PDIM transport to the myco-membrane. Bottom panels show area enclosed in dashed squares on top panels. Arrowheads indicate foci of GAL8-associated Mm. Scale bar, 25 µm.

(B) Percentage of macrophages with foci of GAL8-associated Mm. Symbols represent values from individual imaging fields. Horizontal bars indicate mean values. One-way ANOVA with Tukey’s post-test.

(C) Confocal micrographs of GAL8 immunofluorescence 7 hours after treatment with prazosin (PRZ, 20 µM) or 0.5% DMSO. Arrowheads indicate GAL8 puncta. Scale bar, 20 µm.

**Table S1. Metabolite Profiles of Mm-infected and Uninfected THP-1 Macrophages Treated with Torin1, 2DG, or DMSO, Related to Figure S3**.

**Movie S1. Annexin V Labeling of a Dying, Mm-infected Macrophage in an mTOR Mutant Animal, Related to Figure 2**. Time-lapse confocal microscopy of a dying, infected macrophage in an *mtor*^*sa16755/sa16755*^; *Tg(mfap4:tdTomato-CAAX); Tg(ubib:secA5-YFP)* animal 2 dpi. Mm (blue), secreted annexin V-YFP (green), and macrophage (magenta) are shown.

**Movie S2. Death of Mm-infected Macrophages in mTOR Mutant Animals, Related to Figure 2**. Time-lapse confocal microscopy of *mtor*^*fh178/fh178*^ animals and mTOR-sufficient siblings expressing *Tg(mpeg1:YFP)*, 2 dpi. Mm (magenta) and macrophages (green) are shown. Arrows indicate dying infected macrophages.

**Movie S3. Mitochondrial Damage Precedes Mycobacterium-induced Cell Death in mTOR-deficient Macrophages, Related to Figure 4**. 8-hr time-lapse confocal microscopy of a torin1-treated THP-1 macrophage experiencing mitochondrial damage and subsequently dying (arrow). Note the sequential loss of the mitochondrial dyes TMRE (gold) and MitoTracker Deep Red (magenta) prior to labeling with the cell-membrane impermeant nucleic acid dye Sytox Green. The adjacent viable uninfected cells retained the mitochondrial dyes for the duration of the experiment.

**Movie S4. Death of Mm-infected Macrophages in 2DG-treated Animals, Related to Figure 5**. Time-lapse confocal microscopy of *Tg(mpeg1:YFP)* animals treated with 2DG or DMSO, 3 dpi. Mm (magenta) and macrophages (green) are shown. Arrows indicate dying infected macrophages.

**Movie S5. Torin1-treated THP-1 Macrophages Tolerate** *Δ****ESX-1* Infection, Related to Figure 6**. Time-lapse confocal microscopy of THP-1 macrophage treated with torin1 or DMSO and infected with WT or Δ*ESX-1* Mm, 1 dpi. Mm (magenta) and Sytox Green-labeled dying cells (green). Arrows indicate dying infected macrophages.

**Movie S6. Macrophages of mTOR Mutant Animals Tolerate** *Δ****ESX-1* Infection, Related to Figure 6**. Time-lapse confocal microscopy of *mtor*^*fh178/fh178*^ animals and mTOR-sufficient siblings expressing *Tg(mpeg1:YFP)* infected with WT or *ΔESX-1* Mm, 2 dpi. Mm (magenta) and macrophages (green) are shown. Arrows indicate dying infected macrophages.

## MATERIALS AND METHODS

### RESOURCE AVAILABILITY

Additional information and requests for resources and reagents should be directed to and will be fulfilled by the Lead Contact, Lalita Ramakrishnan.

## EXPERIMENTAL MODEL AND SUBJECT DETAILS

### Ethics Statement

Zebrafish husbandry and experiments were carried out in compliance with guidelines from the UK Home Office and the US Public Health Service Policy on Human Care and Use of Laboratory Larvae using protocols approved by the Animal Welfare and Ethical Review Body of the University of Cambridge and the Institutional Animal Care and Use Committee of the University of Washington, respectively.

### Zebrafish Husbandry and Infections

Zebrafish of the wild-type AB strain (Zebrafish International Resource Center, ZIRC), TL strain (ZIRC), or of mixed AB/TL backgrounds were used in experiments (Table S2). The *Tg(mpeg1:YFP)*^*w200*^, *Tg(mpeg1:Brainbow)*^*w201*^ (described as *mpeg1:tdTomato*), *Tg(mfap4:tdTomato-CAAX)*^*xt6*^, *Tg(CMV:EGFP-map1lc3b)*^*zf155*^, *Tg(lysC:EGFP)*^*nz117*^, *Tg(cd41:GFP)*, and *Tg(ubb:secA5-YFP)*^*cu34*^ (this work) fluorescent reporter lines were maintained in the AB strain (Hall *et al*., 2007; He et al., 2009; Lin *et al*., 2005; Pagan *et al*., 2015; Roca and Ramakrishnan, 2013; Walton et al., 2015) (Table S2). The *pycard*^*w216*^ mutant line (Matty et al., 2019) and *mtor*^*fh178*^ mutant line (this work) were generated and maintained in the AB strain (Table S2). The *mtor*^*sa16755*^, *rptor*^*sa11537*^, *rictora*^*sa15967*^, *rictorb*^*sa18403*^, *atg12*^*sa42684*^, *casp9*^*sa11164*^, *sting1*^*sa35634*^, and *g6pd*^*sa24272*^ mutant lines (Wellcome Trust Sanger Institute) (Kettleborough *et al*., 2013) were generated in the TL strain and maintained in either the TL strain (*rictora*^*sa15967*^, *rictorb*^*sa18403*^, and *g6pd*^*sa24272*^) or mixed AB/TL backgrounds (*mtor*^*sa16755*^, *rptor*^*sa11537*^, and *atg12*^*sa42684*^, *casp9*^*sa11164*^, and *sting1*^*sa35634*^) (Table S2). Zebrafish of the WIK strain (ZIRC) were used to map *mtor*^*fh178*^. Zebrafish were maintained in buffered reverse osmotic water systems under a 14-hr light/10-hr dark cycle. Zebrafish larvae were fed paramecia twice daily, while juvenile and adult zebrafish were fed at least twice a day with dry food and brine shrimp. Zebrafish embryos were collected and cultured in reverse osmosis water containing 0.18g/L Instant Ocean Salt supplemented with 0.25µg/mL methylene blue at 28.5°C. On 1 dpf, embryos to be used in experiments were transferred to 0.5x E2 medium (7.5mM NaCl, 0.25mM KCl, 0.5mM MgSO_4_, 0.075mM KH_2_PO_4_, 0.025mM Na_2_HPO_4_, 0.5mM CaCl_2_, and 0.35mM NaHCO_3_) supplemented with 0.003% PTU (1-phenyl-2-thiourea, Sigma) to inhibit melanin synthesis.

For infections, 2 dpf larvae of undetermined sex (due to early developmental stage) were dechorionated manually or with ≤0.5mg/mL pronase (Sigma-Aldrich) and then anesthetized with fish water containing 0.025% tricaine (Sigma). Larvae were injected via the caudal vein or the hindbrain ventricle using single-cell suspensions of Mm of known titer to deliver ∼100 - 250 bacteria per 3 – 5 nL injection as previously described (Takaki et al., 2013). Phenol red sodium salt (≤ 1% w/v diluted in PBS, Sigma-Aldrich) was used as an injection tracer. Inoculums were confirmed by injecting onto Middlebrook 7H10 agar plates (supplemented with oleic acid, albumin, dextrose, and Tween-80 plus hygromycin B or kanamycin, as appropriate. For experiments with mutant lines, wild-type and heterozygous siblings were used as comparators, animals were genotyped after data acquisition. In experiments involving drug treatments, infected animals were randomly assigned to treatment groups. Unless indicated, treatments (Table S2) were initiated immediately after infection via soaking, and drug exposure was maintained until the experimental endpoint. UK5099 (Cambridge Bioscience) was changed daily.

### THP-1 Macrophage Culture and Infections

Monocytic human THP-1 cells (ATCC) were grown at 37°C, 5% CO_2_ in RPMI medium (Sigma-Aldrich) supplemented with 10% FBS (Gibco), L-glutamine (Gibco), Penicillin, and Streptomycin (complete RPMI) (Table S2). To differentiate THP-1 cells into macrophages, cells were seeded at 5 × 10^5^ cells/mL in flat-bottomed tissue culture plates (2.5 × 10^5^ cells/well of 24-well plate) and stimulated with 100nM PMA Phorbol 12-myristate 13-acetate (Sigma-Aldrich) for two days. The resulting adherent cells were then washed with complete RPMI and rested for two days prior to infection. For microscopy experiments, cells were plated on optical bottom plates (Perkin Elmer or MatTek). Cells were pre-incubated overnight with pharmacological compounds or matching concentrations of vehicle (≤0.5% DMSO). On the infection day, cells were washed with antibiotic-free complete RPMI and infected with single-cell suspensions of Mm or Mtb for five hours in a 33°C, 5% CO_2_ incubator (for Mm) and 37°C, 5% CO_2_ incubator (for Mtb). After infection, cells were washed with antibiotic-free complete RPMI supplemented with the corresponding pharmacological compounds and returned to the appropriate incubators. For experiments with the leucine and pantothenate double auxotroph *ΔleuDΔpanCD* Mtb, complete RMPI was also supplemented with 0.05 mg/mL L-leucine and 0.024 mg/mL calcium pantothenate (Sigma-Aldrich). Multiplicity of infection was determined by calculating the number of mycobacterial colony forming units (CFU) per cells plated in each well.

## METHODS DETAILS

### Bacterial Strains

*M. marinum* M strain (ATCC #BAA-535) and its mutant derivatives *ΔESX-1, ΔmmpL7, Δerp, marP::tn*, and *ΔesxA* expressing BFP2, mWasabi, tdTomato, or tdKatushka2 under the control of the *msp12* promoter (Cambier *et al*., 2014b; Cosma *et al*., 2006; Levitte *et al*., 2016; Osman *et al*., 2022; Takaki *et al*., 2013) were grown at 33°C under hygromycin B (Cambridge Bioscience) or kanamycin (Sigma-Aldrich) selection in Middlebrook 7H9 medium (BD Difco) supplemented with oleic acid, albumin, dextrose, and Tween-80 (Sigma-Aldrich) (Takaki *et al*., 2013) (Table S2). *M. tuberculosis ΔleuDΔpanCD* mc^2^ 6206 expressing *msp12:tdTomato* was grown at 37°C under hygromycin B and kanamycin selection in Middlebrook 7H9 medium supplemented with oleic acid, albumin, dextrose, Tween-80, catalase, and 0.05 mg/mL L-leucine and 0.024 mg/mL calcium pantothenate (Sigma-Aldrich) (Roca *et al*., 2019; Sampson *et al*., 2011).

### Zebrafish Genotyping

DNA from adult fin clips or whole larvae was extracted using the HotSHOT method (Truett et al., 2000). Animals were genotyped by High Resolution Melt Analysis (HRMA) of PCR products (Garritano et al., 2009) or by Kompetitive Allele-Specific PCR (KASP) assay (LGC Biosearch) (He et al., 2014) in a CFX Connect thermocycler (BioRad) (Table S2). DNA from 2DG-treated animals was purified with AMPure XP beads (Beckman Coulter) according to the manufacturer’s instructions prior to PCR.

### Mapping of *mtor*^*fh178*^

Zebrafish carrying the *fh178* allele were outcrossed to the WIK strain for mapping as previously described (Tobin *et al*., 2010). *fh178* was mapped to chromosome 8 initially between the markers z7370 and z14670. Further mapping defined a critical two-gene interval that included *mtor* and *qars1* to the right of a single nucleotide polymorphism in the 3’ end of *angptl7* (1 recombination in 302 meioses) and to the left of a polymorphism in intron 2 of *ogg1* (2 recombinations in 300 meioses). Sequencing of cDNAs isolated from mutants and wild-type animals in the critical region identified a stop codon in exon 24 of *mtor* in the mutant but not wild-type animals. This mutation segregated absolutely with the *fh178* mutants (no recombinants).

### Creation of *Tg(ubb:secA5-YFP)*^*cu34*^

The Tol2 *ubb:secA5-YFP* plasmid was assembled from PCR-amplified fragments of pDestTol2pA2_ubi:EGFP (RRID: Addgene 27323) (Mosimann et al., 2011) and pBH-UAS-secA5-YFP (RRID: Addgene 32359) (van Ham et al., 2010) the Gibson Cloning kit (New England Biolabs) (Table S2). Correct plasmid assembly was confirmed by diagnostic PCR of joined segments and by Sanger sequencing. Tol2 transposase (RRID: Addgene 51818) (Khattak et al., 2014) was in vitro transcribed with the T7 mMessage/mMachine kit (ThermoFisher) according to the manufacturer’s instructions (Table S2). Tol2 *ubb:secA5-YFP* plasmid and transposase mRNA were co-injected into one-cell embryos of the wild-type AB strain as previously described (Suster et al., 2011). G0 larvae expressing the transgene were identified by fluorescence microscopy and raised to adulthood. Potential founders were identified through pairwise crosses of G0 adults and non-transgenic AB fish. A single F1 transgenic animal was used to establish the line.

### CRISPR-Cas9 Mutagenesis

The mutagenesis procedure was adapted from published methods (Burger et al., 2016; Wu et al., 2018). Guide RNAs (60 µM) were generated by complexing equimolar amounts of Alt-R tracrRNA and individual *ndufaf1*-specific Alt-R crRNAs (Dr.Cas9.NDUFAF1.1.AA, AB, AC, and AD; IDT) (Table S2) in nuclease-free Duplex Buffer (IDT) at 95°C for 5 minutes. Aliquots of complexed RNA were stored at -20°C. To produce ribonucleoprotein complexes (RNPs) (10 µM), guide RNA pools were combined with Alt-R Sp Cas9 Nuclease V3 (IDT) (Table S2) at an equimolar ratio of total RNA to Cas9 in working buffer (20 mM HEPES, 150 mM KCl, pH 7.5) and incubated at 37°C for 10 minutes. Wild-type zebrafish of the AB strain were injected at the 1-cell stage with ∼2 nL of RNP to create G0 crispants or ∼2nl of working buffer to produce unmutagenized experimental control animals. Mutagenesis was assessed in individual animals by HRMA of PCR amplicons (Table S2) spanning each of the CRISPR targets.

### Parabiosis

Parabiosis of zebrafish embryos was performed as previously described (Demy et al., 2013). Briefly, blastulae derived from synchronized *mtor*^*fh178/+*^; *Tg(mpeg1:YFP)*^*w200*^ and *Tg(mpeg1:tdTomato)* incrosses were manually dechorionated, paired, scratched with an aluminum silicate microinjection needle, and allowed to fuse in high-calcium Ringer’s solution (116 mM NaCl, 2.9 mM KCl, 10 mM CaCl_2_, 5 mM HEPES, pH 7.2) supplemented with 50 U/mL penicillin and 50 U/mL streptomycin. Only parabiotic pairs fused at the head, devoid of gross morphological abnormalities, and sharing blood circulation were used in experiments. On 2 dpf, each parabiotic animal was infected via the caudal vein to ensure similar distribution of mycobacteria between the pair’s caudal hematopoietic tissues. The bodies of each parabiotic pair were micro-dissected and individually genotyped after completion of the experiment.

### Microscopy and Image Analyses

Widefield fluorescence microscopy was performed as previously described (Takaki *et al*., 2013). Bacterial burdens were determined with a Nikon Eclipse Ti-E inverted microscope using a 4x objective and running a fluorescent pixel counts macro in ImageJ (National Institutes of Health) (Takaki *et al*., 2013). Macrophages were counted manually with a Nikon Eclipse E600 upright microscope using a 20x objective. For serial imaging of individual zebrafish, larvae were housed individually in 96-well plates under standard husbandry conditions.

Laser scanning confocal microscopy was performed as described (Pagan *et al*., 2015). Larvae were anesthetized in PTU-supplemented 0.5x E2 medium containing 0.025% tricaine and embedded in 2% low melt agarose (TopVision) on optical-bottom plates (MatTek). Imaging was performed with a Nikon A1R confocal microscope using 20x or 40x objectives and the galvano or resonant scanners. 0.9 - 2 µm optical sections were combined to generate 15 - 60 µm z stacks. For time-lapse microscopy of zebrafish larvae, imaging was done at 27°C using a microscope incubator (Okolab), using acquisition intervals of 2 - 5 minutes for 6 hours. For serial imaging of individual zebrafish, larvae were removed from the agarose using jeweler’s forceps and housed individually under standard husbandry conditions. For time-lapse microscopy of Mm-infected THP-1 cells, samples were imaged at 33°C, 5% CO_2_ in antibiotic-free, phenol-free complete RPMI containing 50 nM SYTOX Green nucleic acid stain (Invitrogen) for 4 hours.

The acquired images were processed using the denoising feature in the Elements software (Nikon). The surface rendering feature of Imaris (Bitplane) was used to quantify intramacrophage bacterial burdens and mROS levels; the number of acridine orange foci in the midbrain and LC3 puncta in neuromasts, clusters of mechanosensory hair cells of the lateral line in fish; phospho-S6 and total S6 MFIs, and the frequencies of galectin-8-associated Mm per imaging field (Thomas et al., 2015). The numbers of dying macrophages and macrophages with galectin-8-decorated Mm foci were counted manually.

### Acridine Orange Staining

Zebrafish larvae were transferred to 1.5 mL microcentrifuge tubes and stained with 1 mL of 2.4µg/mL acridine orange in 0.5x E2 medium with PTU for 30 minutes with rotation and protected from light (Roca *et al*., 2019). Stained larvae were then rinsed in 0.5x E2 medium with PTU and imaged by confocal microscopy.

### Mitochondrial Vital Dye Staining

Zebrafish larvae were injected with ∼1 nL of 10 mM MitoTracker Red CMXH_2_-Xros via the caudal vein immediately prior to imaging by confocal microscopy (Roca and Ramakrishnan, 2013).

THP-1 macrophages were incubated with 40nM MitoTracker Deep Red (ThermoFisher) and/or 100 nM tetramethylrhodamine, ethyl ester (TMRE) (Abcam) for 20 minutes in a 37°C, 5% CO_2_ incubator according to the manufacturers’ instructions. For flow cytometric analysis of TMRE, cells were then washed with complete RPMI media, stained with FVD eF660, and run without chemical fixation.

### Immunostaining

The staining procedure was performed as previously described (Osman *et al*., 2020). THP-1 cells were fixed in PFA solution (4% w/v paraformaldehyde in PBS) at room temperature (RT) for at least 30 minutes. Cells were then washed twice with PBS and incubated in permeabilization/block (PB) solution (0.1% Triton-X 100, 1% bovine serum albumin in PBS) for 30 minutes at RT, and subsequently stained with Alexa Fluor 647-conjugated rabbit anti-phospho-S6^S235/S236^ or total S6 (Cell Signaling Technology) or goat anti-human galectin-8 antibody (R&D Systems) diluted in PB solution overnight at 4°C. The following day, cells stained with the galectin-8 antibody were washed three times with PBS and stained with Alexa Fluor 488-conjugated donkey anti-goat IgG (ThermoFisher) in PB solution for one hour at RT. Cells were then washed three times with PBS and maintained in PBS for imaging by confocal microscopy. In some experiments, cells nuclei were stained with Hoechst 33342 (10µg/ml) for 20 minutes at RT prior to imaging.

### Flow Cytometry

THP-1 macrophages were stained with Fixable Viability Dye (FVD) eFluor 660 (eBioscience), detached with Accutase (Sigma-Aldrich), and transferred to polystyrene FACS tubes (Sarstedt). Single-cell suspensions were washed with MACS buffer (0.5% bovine serum albumin, 2mM EDTA in PBS, ph7.2) at 290 x g for 5 minutes at 4°C, and in most cases, fixed in 4% PFA solution overnight at 4°C. The following day, samples were washed with and resuspended in MACS buffer. Data were acquired in LSRII or Fortessa flow cytometers (BD Biosciences) and analyzed in FlowJo 10 (TreeStar), gating on single-cell events.

### Cytochrome c Release Assay

The cytochrome c release assay was adapted from Lienard, et al. (Lienard *et al*., 2020). 7 hours after infection with tdTomato-expressing Mm, cells were stained with Fixable Viability Dye (FVD) eFluor 660 (eBioscience), detached with Accutase (Sigma-Aldrich), transferred to polystyrene FACS tubes (Sarstedt), and washed first with ice-cold MACS buffer (0.5% bovine serum albumin, 2mM EDTA in PBS, ph7.2) and then ice-cold PBS by centrifugation at 290 x g for 5 minutes at 4°C. Cells were incubated in ice-cold permeabilization buffer (50 μg/mL digitonin, 100 mM KCl) for 5 minutes and immediately fixed in 4% PFA solution to stop the permeabilization reaction. Cells were then centrifuged as above and fixed overnight in 4% PFA solution at 4°C. Samples were subsequently permeabilized with eBio Permeabilization Buffer (ThermoFisher) and stained with AlexaFluor 488 anti-cytochrome c antibody (BioLegend) (Table S2) for 2 hours at room temperature. Samples were analyzed by flow cytometry by gating on the viable, infected (FVD eFluor 660^-^ tdTomato^+^) single-cell events.

### ATP Quantification

ATP levels were quantified by fluorimetry in a CLARIOStar Plus (BMG Labtech) microplate reader using the ATP Assay Kit (Abcam) according to the manufacturer’s instructions. 10^6^ THP-1 macrophages were analyzed per replicate.

### Measurement of G6PD Activity

Individual 5 dpf larvae from an *g6pd*^*sa24272/+*^ incross were lysed using Mammalian Cell Lysis Buffer (Abcam). A small amount of each lysate was used to genotype the larvae. G6PD activity in individual larvae was determined by fluorimetry in a CLARIOStar Plus (BMG Labtech) microplate reader using the G6PD Activity Assay Kit (Abcam) according to the manufacturer’s instructions.

### Metabolic Analyses

For metabolite quantification, THP-1 cells (2 × 10^6^ cells/sample, four replicates/condition) were washed in ice-cold PBS, and metabolites were extracted with 150 µL extraction buffer (50:30:20, methanol:acetonitrile:water) cooled on dry ice for 30 minutes beforehand. Samples were then centrifuged at maximum speed for 10 minutes at °C to remove protein debris, and supernatants were stored at -80°C until acquisition. Liquid chromatography-mass spectrometry (LC-MS) was carried out using an Agilent 1290 Infinity II UHPLC in line with a Bruker Impact II QTOF operating in negative ion mode, as previously described (Edwards-Hicks et al., 2020). LC separation was performed on a Phenomenex Luna propylamine column (50 × 2 mm, 3 µm particles) using a solvent gradient of 100% buffer B (5 mM ammonium carbonate in 90% acetonitrile) to 90% buffer A (10 mM NH_4_ in water). Flow rate was from 1000 to 750 µL/min. Autosampler temperature was 5°C, and injection volume was 2 µL. Metabolites were quantified using AssayR (Wills et al., 2017). The hierarchical clustering heatmap was generated in MetaboAnalyst 5.0 using Euclidean distance measure, the Ward clustering algorithm, and autoscale feature standarization (Pang et al., 2021).

Metabolic flux measurements were carried out with the Seahorse XFp Glycolytic Rate Assay and Cell Mito Stress Test kits (Agilent) according to the manufacturer’s instructions. On the day of the assay, 4 × 10^4^ differentiated THP-1 macrophages were seeded per well of the XFp cartridge, allowed to adhere for 2 hours in a 37°C, 5% CO_2_ incubator, and then incubated with assay medium (XF DMEM medium with 10 mM glucose and 2 mM glutamine but without sodium pyruvate (Agilent)) for 45 - 60 minutes in a non-CO_2_ 37°C incubator prior to data acquisition in a Seahorse XFp analyzer (Agilent).

### Statistical Analyses

Statistical analyses were performed on Prism (GraphPad) with each statistical test and number of samples per group indicated in the corresponding figure legend. Not significant, *P* > 0.05; \**P < 0*.*05*; \*\**P* < 0.01; *\*\*\*P <* 0.001; *\*\*\*P <* 0.0001.

## REFERENCES

Amaral, E.P., Costa, D.L., Namasivayam, S., Riteau, N., Kamenyeva, O., Mittereder, L., Mayer-Barber, K.D., Andrade, B.B., and Sher, A. (2019). A major role for ferroptosis in Mycobacterium tuberculosis-induced cell death and tissue necrosis. J Exp Med 216, 556–570. 10.1084/jem.20181776.

Armstrong, J.A., and Hart, P.D. (1971). Response of cultured macrophages to Mycobacterium tuberculosis, with observations on fusion of lysosomes with phagosomes. J Exp Med 134, 713–740. 10.1084/jem.134.3.713.

Augenstreich, J., Arbues, A., Simeone, R., Haanappel, E., Wegener, A., Sayes, F., Le Chevalier, F., Chalut, C., Malaga, W., Guilhot, C., et al. (2017). ESX-1 and phthiocerol dimycocerosates of Mycobacterium tuberculosis act in concert to cause phagosomal rupture and host cell apoptosis. Cell Microbiol 19. 10.1111/cmi.12726.

Bao, Y., Wang, L., and Sun, J. (2021). A Small Protein but with Diverse Roles: A Review of EsxA in Mycobacterium-Host Interaction. Cells 10. 10.3390/cells10071645.

Barker, L.P., George, K.M., Falkow, S., and Small, P.L. (1997). Differential trafficking of live and dead Mycobacterium marinum organisms in macrophages. Infect Immun 65, 1497–1504. 10.1128/iai.65.4.1497-1504.1997.

Battaglioni, S., Benjamin, D., Walchli, M., Maier, T., and Hall, M.N. (2022). mTOR substrate phosphorylation in growth control. Cell 185, 1814–1836. 10.1016/j.cell.2022.04.013.

Beckwith, K.S., Beckwith, M.S., Ullmann, S., Saetra, R.S., Kim, H., Marstad, A., Asberg, S.E., Strand, T.A., Haug, M., Niederweis, M., et al. (2020). Plasma membrane damage causes NLRP3 activation and pyroptosis during Mycobacterium tuberculosis infection. Nature communications 11, 2270. 10.1038/s41467-020-16143-6.

Behar, S.M., Divangahi, M., and Remold, H.G. (2010). Evasion of innate immunity by Mycobacterium tuberculosis: is death an exit strategy? Nat Rev Microbiol 8, 668–674. 10.1038/nrmicro2387.

Behr, M.A., Edelstein, P.H., and Ramakrishnan, L. (2018). Revisiting the timetable of tuberculosis. BMJ 362, k2738. 10.1136/bmj.k2738.

Behr, M.A., Edelstein, P.H., and Ramakrishnan, L. (2019). Is Mycobacterium tuberculosis infection life long? BMJ 367, l5770. 10.1136/bmj.l5770.

Behr, M.A., Kaufmann, E., Duffin, J., Edelstein, P.H., and Ramakrishnan, L. (2021). Latent Tuberculosis: Two Centuries of Confusion. Am J Respir Crit Care Med 204, 142–148. 10.1164/rccm.202011-4239PP.

Bendall, L.J., and Green, D.R. (2014). Autopsy of a cell. Leukemia 28, 1341–1343. 10.1038/leu.2014.17.

Benjamin, D., Colombi, M., Moroni, C., and Hall, M.N. (2011). Rapamycin passes the torch: a new generation of mTOR inhibitors. Nat Rev Drug Discov 10, 868–880. 10.1038/nrd3531.

Bento, C.F., Renna, M., Ghislat, G., Puri, C., Ashkenazi, A., Vicinanza, M., Menzies, F.M., and Rubinsztein, D.C. (2016). Mammalian Autophagy: How Does It Work? Annu Rev Biochem 85, 685–713. 10.1146/annurev-biochem-060815-014556.

Berg, R.D., Levitte, S., O’Sullivan, M.P., O’Leary, S.M., Cambier, C.J., Cameron, J., Takaki, K.K., Moens, C.B., Tobin, D.M., Keane, J., and Ramakrishnan, L. (2016). Lysosomal Disorders Drive Susceptibility to Tuberculosis by Compromising Macrophage Migration. Cell 165, 139–152. 10.1016/j.cell.2016.02.034.

Berthet, F.X., Lagranderie, M., Gounon, P., Laurent-Winter, C., Ensergueix, D., Chavarot, P., Thouron, F., Maranghi, E., Pelicic, V., Portnoi, D., et al. (1998). Attenuation of virulence by disruption of the Mycobacterium tuberculosis erp gene. Science 282, 759–762. 10.1126/science.282.5389.759.

Bhatt, K., Bhagavathula, M., Verma, S., Timmins, G.S., Deretic, V.P., Ellner, J.J., and Salgame, P. (2021). Rapamycin modulates pulmonary pathology in a murine model of Mycobacterium tuberculosis infection. Disease models & mechanisms 14. 10.1242/dmm.049018.

Braverman, J., Sogi, K.M., Benjamin, D., Nomura, D.K., and Stanley, S.A. (2016). HIF-1alpha Is an Essential Mediator of IFN-gamma-Dependent Immunity to Mycobacterium tuberculosis. J Immunol 197, 1287–1297. 10.4049/jimmunol.1600266.

Brodin, P., de Jonge, M.I., Majlessi, L., Leclerc, C., Nilges, M., Cole, S.T., and Brosch, R. (2005). Functional analysis of early secreted antigenic target-6, the dominant T-cell antigen of Mycobacterium tuberculosis, reveals key residues involved in secretion, complex formation, virulence, and immunogenicity. J Biol Chem 280, 33953–33959. 10.1074/jbc.M503515200.

Burger, A., Lindsay, H., Felker, A., Hess, C., Anders, C., Chiavacci, E., Zaugg, J., Weber, L.M., Catena, R., Jinek, M., et al. (2016). Maximizing mutagenesis with solubilized CRISPR-Cas9 ribonucleoprotein complexes. Development 143, 2025–2037. 10.1242/dev.134809.

Cambier, C.J., Falkow, S., and Ramakrishnan, L. (2014a). Host evasion and exploitation schemes of Mycobacterium tuberculosis. Cell 159, 1497–1509. 10.1016/j.cell.2014.11.024.

Cambier, C.J., Takaki, K.K., Larson, R.P., Hernandez, R.E., Tobin, D.M., Urdahl, K.B., Cosma, C.L., and Ramakrishnan, L. (2014b). Mycobacteria manipulate macrophage recruitment through coordinated use of membrane lipids. Nature 505, 218–222. 10.1038/nature12799.

Canetti, G., Sutherland, I., and Svandova, E. (1972). Endogenous reactivation and exogenous reinfection: their relative importance with regard to the development of non-primary tuberculosis. Bull Int Union Tuberc 47, 116–134.

Champion, M.M., Williams, E.A., Pinapati, R.S., and Champion, P.A. (2014). Correlation of phenotypic profiles using targeted proteomics identifies mycobacterial esx-1 substrates. J Proteome Res 13, 5151–5164. 10.1021/pr500484w.

Chen, M., Divangahi, M., Gan, H., Shin, D.S., Hong, S., Lee, D.M., Serhan, C.N., Behar, S.M., and Remold, H.G. (2008). Lipid mediators in innate immunity against tuberculosis: opposing roles of PGE2 and LXA4 in the induction of macrophage death. J Exp Med 205, 2791–2801. jem.20080767 [pii] 10.1084/jem.20080767.

Clay, H., Volkman, H.E., and Ramakrishnan, L. (2008). Tumor necrosis factor signaling mediates resistance to mycobacteria by inhibiting bacterial growth and macrophage death. Immunity 29, 283–294.

Clemens, D.L., and Horwitz, M.A. (1995). Characterization of the Mycobacterium tuberculosis phagosome and evidence that phagosomal maturation is inhibited. J Exp Med 181, 257–270. 10.1084/jem.181.1.257.

Clements, W.K., and Traver, D. (2013). Signalling pathways that control vertebrate haematopoietic stem cell specification. Nat Rev Immunol 13, 336–348. 10.1038/nri3443.

Conrad, W.H., Osman, M.M., Shanahan, J.K., Chu, F., Takaki, K.K., Cameron, J., Hopkinson-Woolley, D., Brosch, R., and Ramakrishnan, L. (2017). Mycobacterial ESX-1 secretion system mediates host cell lysis through bacterium contact-dependent gross membrane disruptions. Proc Natl Acad Sci U S A 114, 1371–1376. 10.1073/pnas.1620133114.

Cosma, C.L., Klein, K., Kim, R., Beery, D., and Ramakrishnan, L. (2006). Mycobacterium marinum Erp is a virulence determinant required for cell wall integrity and intracellular survival. Infect Immun 74, 3125–3133. 74/6/3125 [pii] 10.1128/IAI.02061-05.

Cosma, C.L., Sherman, D.R., and Ramakrishnan, L. (2003). The secret lives of the pathogenic mycobacteria. Annu Rev Microbiol 57, 641–676.

Cumming, B.M., Addicott, K.W., Adamson, J.H., and Steyn, A.J. (2018). Mycobacterium tuberculosis induces decelerated bioenergetic metabolism in human macrophages. Elife 7. 10.7554/eLife.39169.

Cunningham, J.T., Rodgers, J.T., Arlow, D.H., Vazquez, F., Mootha, V.K., and Puigserver, P. (2007). mTOR controls mitochondrial oxidative function through a YY1-PGC-1alpha transcriptional complex. Nature 450, 736–740. 10.1038/nature06322.

Davis, J.M., Clay, H., Lewis, J.L., Ghori, N., Herbomel, P., and Ramakrishnan, L. (2002). Real-time visualization of mycobacterium-macrophage interactions leading to initiation of granuloma formation in zebrafish embryos. Immunity 17, 693–702.

Davis, J.M., and Ramakrishnan, L. (2009). The role of the granuloma in expansion and dissemination of early tuberculous infection. Cell 136, 37–49.

Decout, A., Katz, J.D., Venkatraman, S., and Ablasser, A. (2021). The cGAS-STING pathway as a therapeutic target in inflammatory diseases. Nat Rev Immunol 21, 548–569. 10.1038/s41577-021-00524-z.

Demy, D.L., Ranta, Z., Giorgi, J.M., Gonzalez, M., Herbomel, P., and Kissa, K. (2013). Generating parabiotic zebrafish embryos for cell migration and homing studies. Nature methods 10, 256–258. 10.1038/nmeth.2362.

Divangahi, M., Chen, M., Gan, H., Desjardins, D., Hickman, T.T., Lee, D.M., Fortune, S., Behar, S.M., and Remold, H.G. (2009). Mycobacterium tuberculosis evades macrophage defenses by inhibiting plasma membrane repair. Nat Immunol 10, 899–906. 10.1038/ni.1758.

Duvel, K., Yecies, J.L., Menon, S., Raman, P., Lipovsky, A.I., Souza, A.L., Triantafellow, E., Ma, Q., Gorski, R., Cleaver, S., et al. (2010). Activation of a metabolic gene regulatory network downstream of mTOR complex 1. Mol Cell 39, 171–183. 10.1016/j.molcel.2010.06.022.

Edwards-Hicks, J., Mitterer, M., Pearce, E.L., and Buescher, J.M. (2020). Metabolic Dynamics of In Vitro CD8+ T Cell Activation. Metabolites 11. 10.3390/metabo11010012.

Feldman, W.H., and Baggenstoss, A.H. (1938). The reisdual infectivity of the primary complex of tuberculosis. Am. J. Pathol. 14, 473–490.

Fijalkowska-Morawska, J.B., Jagodzinska, M., and Nowicki, M. (2011). Pulmonary embolism and reactivation of tuberculosis during everolimus therapy in a kidney transplant recipient. Ann Transplant 16, 107–110. 10.12659/aot.882226.

Fine-Coulson, K., Giguere, S., Quinn, F.D., and Reaves, B.J. (2015). Infection of A549 human type II epithelial cells with Mycobacterium tuberculosis induces changes in mitochondrial morphology, distribution and mass that are dependent on the early secreted antigen, ESAT-6. Microbes Infect 17, 689–697. 10.1016/j.micinf.2015.06.003.

Formosa, L.E., Dibley, M.G., Stroud, D.A., and Ryan, M.T. (2018). Building a complex complex: Assembly of mitochondrial respiratory chain complex I. Semin Cell Dev Biol 76, 154–162. 10.1016/j.semcdb.2017.08.011.

Fortune, S.M., Jaeger, A., Sarracino, D.A., Chase, M.R., Sassetti, C.M., Sherman, D.R., Bloom, B.R., and Rubin, E.J. (2005). Mutually dependent secretion of proteins required for mycobacterial virulence. Proc Natl Acad Sci U S A 102, 10676–10681. 10.1073/pnas.0504922102.

Galluzzi, L., Bravo-San Pedro, J.M., Kepp, O., and Kroemer, G. (2016). Regulated cell death and adaptive stress responses. Cell Mol Life Sci 73, 2405–2410. 10.1007/s00018-016-2209-y.

Garcia, C.A., and Wu, S. (2016). Attributable Risk of Infection to mTOR Inhibitors Everolimus and Temsirolimus in the Treatment of Cancer. Cancer Invest 34, 521–530. 10.1080/07357907.2016.1242009.

Garritano, S., Gemignani, F., Voegele, C., Nguyen-Dumont, T., Le Calvez-Kelm, F., De Silva, D., Lesueur, F., Landi, S., and Tavtigian, S.V. (2009). Determining the effectiveness of High Resolution Melting analysis for SNP genotyping and mutation scanning at the TP53 locus. BMC Genet 10, 5. 10.1186/1471-2156-10-5.

Gleeson, L.E., Sheedy, F.J., Palsson-McDermott, E.M., Triglia, D., O’Leary, S.M., O’Sullivan, M.P., O’Neill, L.A., and Keane, J. (2016). Cutting Edge: Mycobacterium tuberculosis Induces Aerobic Glycolysis in Human Alveolar Macrophages That Is Required for Control of Intracellular Bacillary Replication. J Immunol 196, 2444–2449. 10.4049/jimmunol.1501612.

Gonzalez, A., Hall, M.N., Lin, S.C., and Hardie, D.G. (2020). AMPK and TOR: The Yin and Yang of Cellular Nutrient Sensing and Growth Control. Cell Metab 31, 472–492. 10.1016/j.cmet.2020.01.015.

Gottlieb, E., Armour, S.M., Harris, M.H., and Thompson, C.B. (2003). Mitochondrial membrane potential regulates matrix configuration and cytochrome c release during apoptosis. Cell Death Differ 10, 709–717. 10.1038/sj.cdd.4401231.

Green, D.R., and Levine, B. (2014). To be or not to be? How selective autophagy and cell death govern cell fate. Cell 157, 65–75. 10.1016/j.cell.2014.02.049.

Groschel, M.I., Sayes, F., Simeone, R., Majlessi, L., and Brosch, R. (2016). ESX secretion systems: mycobacterial evolution to counter host immunity. Nat Rev Microbiol 14, 677–691. 10.1038/nrmicro.2016.131.

Halestrap, A.P. (1975). The mitochondrial pyruvate carrier. Kinetics and specificity for substrates and inhibitors. The Biochemical journal 148, 85–96. 10.1042/bj1480085.

Hall, C., Flores, M.V., Storm, T., Crosier, K., and Crosier, P. (2007). The zebrafish lysozyme C promoter drives myeloid-specific expression in transgenic fish. BMC Dev Biol 7, 42. 10.1186/1471-213X-7-42.

Harris, J., Hope, J.C., and Keane, J. (2008). Tumor necrosis factor blockers influence macrophage responses to Mycobacterium tuberculosis. J Infect Dis 198, 1842–1850. 10.1086/593174.

He, C., Bartholomew, C.R., Zhou, W., and Klionsky, D.J. (2009). Assaying autophagic activity in transgenic GFP-Lc3 and GFP-Gabarap zebrafish embryos. Autophagy 5, 520–526. 10.4161/auto.5.4.7768.

He, C., Holme, J., and Anthony, J. (2014). SNP genotyping: the KASP assay. Methods Mol Biol 1145, 75–86. 10.1007/978-1-4939-0446-4_7.

Hosoi, H., Dilling, M.B., Shikata, T., Liu, L.N., Shu, L., Ashmun, R.A., Germain, G.S., Abraham, R.T., and Houghton, P.J. (1999). Rapamycin causes poorly reversible inhibition of mTOR and induces p53-independent apoptosis in human rhabdomyosarcoma cells. Cancer Res 59, 886–894.

Huang, L., Nazarova, E.V., Tan, S., Liu, Y., and Russell, D.G. (2018). Growth of Mycobacterium tuberculosis in vivo segregates with host macrophage metabolism and ontogeny. J Exp Med 215, 1135–1152. 10.1084/jem.20172020.

Jamwal, S., Midha, M.K., Verma, H.N., Basu, A., Rao, K.V., and Manivel, V. (2013). Characterizing virulence-specific perturbations in the mitochondrial function of macrophages infected with Mycobacterium tuberculosis. Scientific reports 3, 1328. 10.1038/srep01328.

Jeon, S.Y., Yhim, H.Y., Lee, N.R., Song, E.K., Kwak, J.Y., and Yim, C.Y. (2017). Everolimus-induced activation of latent Mycobacterium tuberculosis infection in a patient with metastatic renal cell carcinoma. Korean J Intern Med 32, 365–368. 10.3904/kjim.2015.121.

Karmaus, P.W.F., Herrada, A.A., Guy, C., Neale, G., Dhungana, Y., Long, L., Vogel, P., Avila, J., Clish, C.B., and Chi, H. (2017). Critical roles of mTORC1 signaling and metabolic reprogramming for M-CSF-mediated myelopoiesis. J Exp Med 214, 2629–2647. 10.1084/jem.20161855.

Kettleborough, R.N., Busch-Nentwich, E.M., Harvey, S.A., Dooley, C.M., de Bruijn, E., van Eeden, F., Sealy, I., White, R.J., Herd, C., Nijman, I.J., et al. (2013). A systematic genome-wide analysis of zebrafish protein-coding gene function. Nature 496, 494–497. 10.1038/nature11992.

Khattak, S., Murawala, P., Andreas, H., Kappert, V., Schuez, M., Sandoval-Guzman, T., Crawford, K., and Tanaka, E.M. (2014). Optimized axolotl (Ambystoma mexicanum) husbandry, breeding, metamorphosis, transgenesis and tamoxifen-mediated recombination. Nature protocols 9, 529–540. 10.1038/nprot.2014.040.

Kozik, P., Gros, M., Itzhak, D.N., Joannas, L., Heurtebise-Chretien, S., Krawczyk, P.A., Rodriguez-Silvestre, P., Alloatti, A., Magalhaes, J.G., Del Nery, E., et al. (2020). Small Molecule Enhancers of Endosome-to-Cytosol Import Augment Anti-tumor Immunity. Cell reports 32, 107905. 10.1016/j.celrep.2020.107905.

Lachmandas, E., Beigier-Bompadre, M., Cheng, S.C., Kumar, V., van Laarhoven, A., Wang, X., Ammerdorffer, A., Boutens, L., de Jong, D., Kanneganti, T.D., et al. (2016). Rewiring cellular metabolism via the AKT/mTOR pathway contributes to host defence against Mycobacterium tuberculosis in human and murine cells. Eur J Immunol 46, 2574–2586. 10.1002/eji.201546259.

Lee, J., Repasy, T., Papavinasasundaram, K., Sassetti, C., and Kornfeld, H. (2011). Mycobacterium tuberculosis induces an atypical cell death mode to escape from infected macrophages. PLoS One 6, e18367. 10.1371/journal.pone.0018367.

Lee, P.Y., Sykes, D.B., Ameri, S., Kalaitzidis, D., Charles, J.F., Nelson-Maney, N., Wei, K., Cunin, P., Morris, A., Cardona, A.E., et al. (2017). The metabolic regulator mTORC1 controls terminal myeloid differentiation. Science immunology 2. 10.1126/sciimmunol.aam6641.

Lerner, T.R., Queval, C.J., Fearns, A., Repnik, U., Griffiths, G., and Gutierrez, M.G. (2018). Phthiocerol dimycocerosates promote access to the cytosol and intracellular burden of Mycobacterium tuberculosis in lymphatic endothelial cells. BMC Biol 16, 1. 10.1186/s12915-017-0471-6.

Levitte, S., Adams, K.N., Berg, R.D., Cosma, C.L., Urdahl, K.B., and Ramakrishnan, L. (2016). Mycobacterial Acid Tolerance Enables Phagolysosomal Survival and Establishment of Tuberculous Infection In Vivo. Cell Host Microbe 20, 250–258. 10.1016/j.chom.2016.07.007.

Lienard, J., Nobs, E., Lovins, V., Movert, E., Valfridsson, C., and Carlsson, F. (2020). The Mycobacterium marinum ESX-1 system mediates phagosomal permeabilization and type I interferon production via separable mechanisms. Proc Natl Acad Sci U S A 117, 1160–1166. 10.1073/pnas.1911646117.

Lin, H.F., Traver, D., Zhu, H., Dooley, K., Paw, B.H., Zon, L.I., and Handin, R.I. (2005). Analysis of thrombocyte development in CD41-GFP transgenic zebrafish. Blood 106, 3803–3810. 10.1182/blood-2005-01-0179.

Liu, G.Y., and Sabatini, D.M. (2020). mTOR at the nexus of nutrition, growth, ageing and disease. Nat Rev Mol Cell Biol 21, 183–203. 10.1038/s41580-019-0199-y.

Ma, D., Zhang, J., Lin, H.F., Italiano, J., and Handin, R.I. (2011). The identification and characterization of zebrafish hematopoietic stem cells. Blood 118, 289–297. 10.1182/blood-2010-12-327403.

Mannick, J.B., Del Giudice, G., Lattanzi, M., Valiante, N.M., Praestgaard, J., Huang, B., Lonetto, M.A., Maecker, H.T., Kovarik, J., Carson, S., et al. (2014). mTOR inhibition improves immune function in the elderly. Science translational medicine 6, 268ra179. 10.1126/scitranslmed.3009892.

Mannick, J.B., Morris, M., Hockey, H.P., Roma, G., Beibel, M., Kulmatycki, K., Watkins, M., Shavlakadze, T., Zhou, W., Quinn, D., et al. (2018). TORC1 inhibition enhances immune function and reduces infections in the elderly. Science translational medicine 10. 10.1126/scitranslmed.aaq1564.

Matty, M.A., Knudsen, D.R., Walton, E.M., Beerman, R.W., Cronan, M.R., Pyle, C.J., Hernandez, R.E., and Tobin, D.M. (2019). Potentiation of P2RX7 as a host-directed strategy for control of mycobacterial infection. Elife 8. 10.7554/eLife.39123.

Mohareer, K., Medikonda, J., Vadankula, G.R., and Banerjee, S. (2020). Mycobacterial Control of Host Mitochondria: Bioenergetic and Metabolic Changes Shaping Cell Fate and Infection Outcome. Front Cell Infect Microbiol 10, 457. 10.3389/fcimb.2020.00457.

Morita, M., Gravel, S.P., Chenard, V., Sikstrom, K., Zheng, L., Alain, T., Gandin, V., Avizonis, D., Arguello, M., Zakaria, C., et al. (2013). mTORC1 controls mitochondrial activity and biogenesis through 4E-BP-dependent translational regulation. Cell Metab 18, 698–711. 10.1016/j.cmet.2013.10.001.

Morita, M., Prudent, J., Basu, K., Goyon, V., Katsumura, S., Hulea, L., Pearl, D., Siddiqui, N., Strack, S., McGuirk, S., et al. (2017). mTOR Controls Mitochondrial Dynamics and Cell Survival via MTFP1. Mol Cell 67, 922–935 e925. 10.1016/j.molcel.2017.08.013.

Mosimann, C., Kaufman, C.K., Li, P., Pugach, E.K., Tamplin, O.J., and Zon, L.I. (2011). Ubiquitous transgene expression and Cre-based recombination driven by the ubiquitin promoter in zebrafish. Development 138, 169–177. 10.1242/dev.059345.

Mouton, J.M., Heunis, T., Dippenaar, A., Gallant, J.L., Kleynhans, L., and Sampson, S.L. (2019). Comprehensive Characterization of the Attenuated Double Auxotroph Mycobacterium tuberculosisDeltaleuDDeltapanCD as an Alternative to H37Rv. Front Microbiol 10, 1922. 10.3389/fmicb.2019.01922.

Murphy, M.P. (2009). How mitochondria produce reactive oxygen species. The Biochemical journal 417, 1–13. 10.1042/BJ20081386.

Muthukkumar, S., Ramesh, T.M., and Bondada, S. (1995). Rapamycin, a potent immunosuppressive drug, causes programmed cell death in B lymphoma cells. Transplantation 60, 264–270. 10.1097/00007890-199508000-00010.

O’Neill, L.A., and Pearce, E.J. (2016). Immunometabolism governs dendritic cell and macrophage function. J Exp Med 213, 15–23. 10.1084/jem.20151570.

Olson, G.S., Murray, T.A., Jahn, A.N., Mai, D., Diercks, A.H., Gold, E.S., and Aderem, A. (2021). Type I interferon decreases macrophage energy metabolism during mycobacterial infection. Cell reports 35, 109195. 10.1016/j.celrep.2021.109195.

Ong, C.W., Elkington, P.T., and Friedland, J.S. (2014). Tuberculosis, pulmonary cavitation, and matrix metalloproteinases. Am J Respir Crit Care Med 190, 9–18. 10.1164/rccm.201311-2106PP.

Opie, E.L., and Aronson, J.D. (1927). Tubercle bacilli in latent tuberculous lesions and in lung tissue without tuberculous lesions. Arch Pathol Lab Med 4, 1–21.

Osman, M.M., Pagan, A.J., Shanahan, J.K., and Ramakrishnan, L. (2020). Mycobacterium marinum phthiocerol dimycocerosates enhance macrophage phagosomal permeabilization and membrane damage. PLoS One 15, e0233252. 10.1371/journal.pone.0233252.

Osman, M.M., Shanahan, J.K., Chu, F., Takaki, K.K., Pinckert, M.L., Pagan, A.J., Brosch, R., Conrad, W.H., and Ramakrishnan, L. (2022). The C terminus of the mycobacterium ESX-1 secretion system substrate ESAT-6 is required for phagosomal membrane damage and virulence. Proc Natl Acad Sci U S A 119, e2122161119. 10.1073/pnas.2122161119.

Pagan, A.J., and Ramakrishnan, L. (2014). Immunity and Immunopathology in the Tuberculous Granuloma. Cold Spring Harbor perspectives in medicine 5. 10.1101/cshperspect.a018499.

Pagan, A.J., and Ramakrishnan, L. (2018). The Formation and Function of Granulomas. Annu Rev Immunol 36, 639–665. 10.1146/annurev-immunol-032712-100022.

Pagan, A.J., Yang, C.T., Cameron, J., Swaim, L.E., Ellett, F., Lieschke, G.J., and Ramakrishnan, L. (2015). Myeloid Growth Factors Promote Resistance to Mycobacterial Infection by Curtailing Granuloma Necrosis through Macrophage Replenishment. Cell Host Microbe 18, 15–26. 10.1016/j.chom.2015.06.008.

Pajuelo, D., Gonzalez-Juarbe, N., Tak, U., Sun, J., Orihuela, C.J., and Niederweis, M. (2018). NAD(+) Depletion Triggers Macrophage Necroptosis, a Cell Death Pathway Exploited by Mycobacterium tuberculosis. Cell reports 24, 429–440. 10.1016/j.celrep.2018.06.042.

Pang, Z., Chong, J., Zhou, G., de Lima Morais, D.A., Chang, L., Barrette, M., Gauthier, C., Jacques, P.E., Li, S., and Xia, J. (2021). MetaboAnalyst 5.0: narrowing the gap between raw spectra and functional insights. Nucleic Acids Res 49, W388–W396. 10.1093/nar/gkab382.

Papadopoli, D., Boulay, K., Kazak, L., Pollak, M., Mallette, F., Topisirovic, I., and Hulea, L. (2019). mTOR as a central regulator of lifespan and aging. F1000Res 8. 10.12688/f1000research.17196.1.

Patra, K.C., and Hay, N. (2014). The pentose phosphate pathway and cancer. Trends Biochem Sci 39, 347–354. 10.1016/j.tibs.2014.06.005.

Paulson, T. (2013). Epidemiology: A mortal foe. Nature 502, S2–3. 10.1038/502S2a.

Pernas, L. (2021). Cellular metabolism in the defense against microbes. J Cell Sci 134. 10.1242/jcs.252023.

Powell, J.D., Pollizzi, K.N., Heikamp, E.B., and Horton, M.R. (2012). Regulation of immune responses by mTOR. Annu Rev Immunol 30, 39–68. 10.1146/annurev-immunol-020711-075024.

Quigley, J., Hughitt, V.K., Velikovsky, C.A., Mariuzza, R.A., El-Sayed, N.M., and Briken, V. (2017). The Cell Wall Lipid PDIM Contributes to Phagosomal Escape and Host Cell Exit of Mycobacterium tuberculosis. mBio 8. 10.1128/mBio.00148-17.

Ramakrishnan, L. (2012). Revisiting the role of the granuloma in tuberculosis. Nat Rev Immunol 12, 352–366. 10.1038/nri3211.

Ramakrishnan, L. (2013). The zebrafish guide to tuberculosis immunity and treatment. Cold Spring Harb Symp Quant Biol 78, 179–192. 10.1101/sqb.2013.78.023283.

Ramakrishnan, L. (2020). Mycobacterium tuberculosis pathogenicity viewed through the lens of molecular Koch’s postulates. Curr Opin Microbiol 54, 103–110. 10.1016/j.mib.2020.01.011.

Rambold, A.S., and Pearce, E.L. (2018). Mitochondrial Dynamics at the Interface of Immune Cell Metabolism and Function. Trends Immunol 39, 6–18. 10.1016/j.it.2017.08.006.

Roca, F.J., and Ramakrishnan, L. (2013). TNF dually mediates resistance and susceptibility to mycobacteria via mitochondrial reactive oxygen species. Cell 153, 521–534. 10.1016/j.cell.2013.03.022.

Roca, F.J., Whitworth, L.J., Prag, H.A., Murphy, M.P., and Ramakrishnan, L. (2022). Tumor necrosis factor induces pathogenic mitochondrial ROS in tuberculosis through reverse electron transport. Science 376, eabh2841. 10.1126/science.abh2841.

Roca, F.J., Whitworth, L.J., Redmond, S., Jones, A.A., and Ramakrishnan, L. (2019). TNF Induces Pathogenic Programmed Macrophage Necrosis in Tuberculosis through a Mitochondrial-Lysosomal-Endoplasmic Reticulum Circuit. Cell 178, 1344–1361 e1311. 10.1016/j.cell.2019.08.004.

Ruiz-Camps, I., and Aguilar-Company, J. (2021). Risk of infection associated with targeted therapies for solid organ and hematological malignancies. Ther Adv Infect Dis 8, 2049936121989548. 10.1177/2049936121989548.

Russell, D.G. (2007). Who puts the tubercle in tuberculosis? Nat Rev Microbiol 5, 39–47.

Ryan, D.G., and O’Neill, L.A.J. (2020). Krebs Cycle Reborn in Macrophage Immunometabolism. Annu Rev Immunol 38, 289–313. 10.1146/annurev-immunol-081619-104850.

Sampson, S.L., Dascher, C.C., Sambandamurthy, V.K., Russell, R.G., Jacobs, W.R., Jr., Bloom, B.R., and Hondalus, M.K. (2004). Protection elicited by a double leucine and pantothenate auxotroph of Mycobacterium tuberculosis in guinea pigs. Infect Immun 72, 3031–3037. 10.1128/IAI.72.5.3031-3037.2004.

Sampson, S.L., Mansfield, K.G., Carville, A., Magee, D.M., Quitugua, T., Howerth, E.W., Bloom, B.R., and Hondalus, M.K. (2011). Extended safety and efficacy studies of a live attenuated double leucine and pantothenate auxotroph of Mycobacterium tuberculosis as a vaccine candidate. Vaccine 29, 4839–4847. 10.1016/j.vaccine.2011.04.066.

Saxton, R.A., and Sabatini, D.M. (2017). mTOR Signaling in Growth, Metabolism, and Disease. Cell 168, 960–976. 10.1016/j.cell.2017.02.004.

Schieke, S.M., Phillips, D., McCoy, J.P., Jr., Aponte, A.M., Shen, R.F., Balaban, R.S., and Finkel, T. (2006). The mammalian target of rapamycin (mTOR) pathway regulates mitochondrial oxygen consumption and oxidative capacity. J Biol Chem 281, 27643–27652. 10.1074/jbc.M603536200.

Shi, L., Salamon, H., Eugenin, E.A., Pine, R., Cooper, A., and Gennaro, M.L. (2015). Infection with Mycobacterium tuberculosis induces the Warburg effect in mouse lungs. Scientific reports 5, 18176. 10.1038/srep18176.

Simeone, R., Bobard, A., Lippmann, J., Bitter, W., Majlessi, L., Brosch, R., and Enninga, J. (2012). Phagosomal rupture by Mycobacterium tuberculosis results in toxicity and host cell death. PLoS Pathog 8, e1002507. 10.1371/journal.ppat.1002507.

Simeone, R., Sayes, F., Lawaree, E., and Brosch, R. (2021). Breaching the phagosome, the case of the tuberculosis agent. Cell Microbiol 23, e13344. 10.1111/cmi.13344.

Sinclair, C., Bommakanti, G., Gardinassi, L., Loebbermann, J., Johnson, M.J., Hakimpour, P., Hagan, T., Benitez, L., Todor, A., Machiah, D., et al. (2017). mTOR regulates metabolic adaptation of APCs in the lung and controls the outcome of allergic inflammation. Science 357, 1014–1021. 10.1126/science.aaj2155.

Srinivasan, L., Ahlbrand, S., and Briken, V. (2014). Interaction of Mycobacterium tuberculosis with host cell death pathways. Cold Spring Harbor perspectives in medicine 4. 10.1101/cshperspect.a022459.

Suster, M.L., Abe, G., Schouw, A., and Kawakami, K. (2011). Transposon-mediated BAC transgenesis in zebrafish. Nature protocols 6, 1998–2021. 10.1038/nprot.2011.416.

Swaim, L.E., Connolly, L.E., Volkman, H.E., Humbert, O., Born, D.E., and Ramakrishnan, L. (2006). Mycobacterium marinum infection of adult zebrafish causes caseating granulomatous tuberculosis and is moderated by adaptive immunity. Infect Immun 74, 6108–6117.

Swanson, K.V., Deng, M., and Ting, J.P. (2019). The NLRP3 inflammasome: molecular activation and regulation to therapeutics. Nat Rev Immunol 19, 477–489. 10.1038/s41577-019-0165-0.

Takaki, K., Davis, J.M., Winglee, K., and Ramakrishnan, L. (2013). Evaluation of the pathogenesis and treatment of Mycobacterium marinum infection in zebrafish. Nature protocols 8, 1114–1124. 10.1038/nprot.2013.068.

Terplan, K. (1951). [Pathogenesis of postprimary tuberculosis, in relation to chronic pulmonary tuberculosis (phtisis)]. Bibl Tuberc 5, 186–219.

Thomas, E.D., Cruz, I.A., Hailey, D.W., and Raible, D.W. (2015). There and back again: development and regeneration of the zebrafish lateral line system. Wiley Interdiscip Rev Dev Biol 4, 1–16. 10.1002/wdev.160.

Tobin, D.M., Roca, F.J., Oh, S.F., McFarland, R., Vickery, T.W., Ray, J.P., Ko, D.C., Zou, Y., Bang, N.D., Chau, T.T., et al. (2012). Host genotype-specific therapies can optimize the inflammatory response to mycobacterial infections. Cell 148, 434–446. 10.1016/j.cell.2011.12.023.

Tobin, D.M., Vary, J.C., Jr., Ray, J.P., Walsh, G.S., Dunstan, S.J., Bang, N.D., Hagge, D.A., Khadge, S., King, M.C., Hawn, T.R., et al. (2010). The lta4h locus modulates susceptibility to mycobacterial infection in zebrafish and humans. Cell 140, 717–730. 10.1016/j.cell.2010.02.013.

Truett, G.E., Heeger, P., Mynatt, R.L., Truett, A.A., Walker, J.A., and Warman, M.L. (2000). Preparation of PCR-quality mouse genomic DNA with hot sodium hydroxide and tris (HotSHOT). BioTechniques 29, 52, 54. 10.2144/00291bm09.

Tsai, M.K., Lee, C.Y., Hu, R.H., and Lee, P.H. (2007). Conversion to combined therapy with sirolimus and mycophenolate mofetil improved renal function in stable renal transplant recipients. J Formos Med Assoc 106, 372–379. 10.1016/S0929-6646(09)60322-3.

van Ham, T.J., Mapes, J., Kokel, D., and Peterson, R.T. (2010). Live imaging of apoptotic cells in zebrafish. FASEB J 24, 4336–4342. 10.1096/fj.10-161018.

Vandal, O.H., Pierini, L.M., Schnappinger, D., Nathan, C.F., and Ehrt, S. (2008). A membrane protein preserves intrabacterial pH in intraphagosomal Mycobacterium tuberculosis. Nat Med 14, 849–854. 10.1038/nm.1795.

Volkman, H.E., Clay, H., Beery, D., Chang, J.C., Sherman, D.R., and Ramakrishnan, L. (2004). Tuberculous granuloma formation is enhanced by a mycobacterium virulence determinant. PLoS Biol 2, e367.

Wallis, R.S., Ginindza, S., Beattie, T., Arjun, N., Likoti, M., Edward, V.A., Rassool, M., Ahmed, K., Fielding, K., Ahidjo, B.A., et al. (2021). Adjunctive host-directed therapies for pulmonary tuberculosis: a prospective, open-label, phase 2, randomised controlled trial. Lancet Respir Med 9, 897–908. 10.1016/S2213-2600(20)30448-3.

Walton, E.M., Cronan, M.R., Beerman, R.W., and Tobin, D.M. (2015). The Macrophage-Specific Promoter mfap4 Allows Live, Long-Term Analysis of Macrophage Behavior during Mycobacterial Infection in Zebrafish. PLoS One 10, e0138949. 10.1371/journal.pone.0138949.

Weichhart, T., Hengstschlager, M., and Linke, M. (2015). Regulation of innate immune cell function by mTOR. Nat Rev Immunol 15, 599–614. 10.1038/nri3901.

Whitworth, L., Coxon, J., van Laarhoven, A., Thuong, N.T.T., Dian, S., Alisjahbana, B., Ganiem, A.R., van Crevel, R., Thwaites, G.E., Troll, M., et al. (2021a). A Bayesian analysis of the association between Leukotriene A4 Hydrolase genotype and survival in tuberculous meningitis. Elife 10. 10.7554/eLife.61722.

Whitworth, L.J., Troll, R., Pagan, A.J., Roca, F.J., Edelstein, P.H., Troll, M., Tobin, D.M., Phu, N.H., Bang, N.D., Thwaites, G.E., et al. (2021b). Elevated cerebrospinal fluid cytokine levels in tuberculous meningitis predict survival in response to dexamethasone. Proc Natl Acad Sci U S A 118. 10.1073/pnas.2024852118.

Wiens, K.E., and Ernst, J.D. (2016). The Mechanism for Type I Interferon Induction by Mycobacterium tuberculosis is Bacterial Strain-Dependent. PLoS Pathog 12, e1005809. 10.1371/journal.ppat.1005809.

Wills, J., Edwards-Hicks, J., and Finch, A.J. (2017). AssayR: A Simple Mass Spectrometry Software Tool for Targeted Metabolic and Stable Isotope Tracer Analyses. Anal Chem 89, 9616–9619. 10.1021/acs.analchem.7b02401.

Wong, Y.C., Ysselstein, D., and Krainc, D. (2018). Mitochondria-lysosome contacts regulate mitochondrial fission via RAB7 GTP hydrolysis. Nature 554, 382–386. 10.1038/nature25486.

Wu, R.S., Lam, II, Clay, H., Duong, D.N., Deo, R.C., and Coughlin, S.R. (2018). A Rapid Method for Directed Gene Knockout for Screening in G0 Zebrafish. Dev Cell 46, 112–125 e114. 10.1016/j.devcel.2018.06.003.

Zhang, L., Jiang, X., Pfau, D., Ling, Y., and Nathan, C.F. (2021). Type I interferon signaling mediates Mycobacterium tuberculosis-induced macrophage death. J Exp Med 218. 10.1084/jem.20200887.

Zhang, Q., Wang, D., Jiang, G., Liu, W., Deng, Q., Li, X., Qian, W., Ouellet, H., and Sun, J. (2016). EsxA membrane-permeabilizing activity plays a key role in mycobacterial cytosolic translocation and virulence: effects of single-residue mutations at glutamine 5. Scientific reports 6, 32618. 10.1038/srep32618.

