## Supplementary figures and images for "mTOR-regulated Mitochondrial Metabolism Limits Mycobacterium-induced Cytotoxicity"

### Figure S1

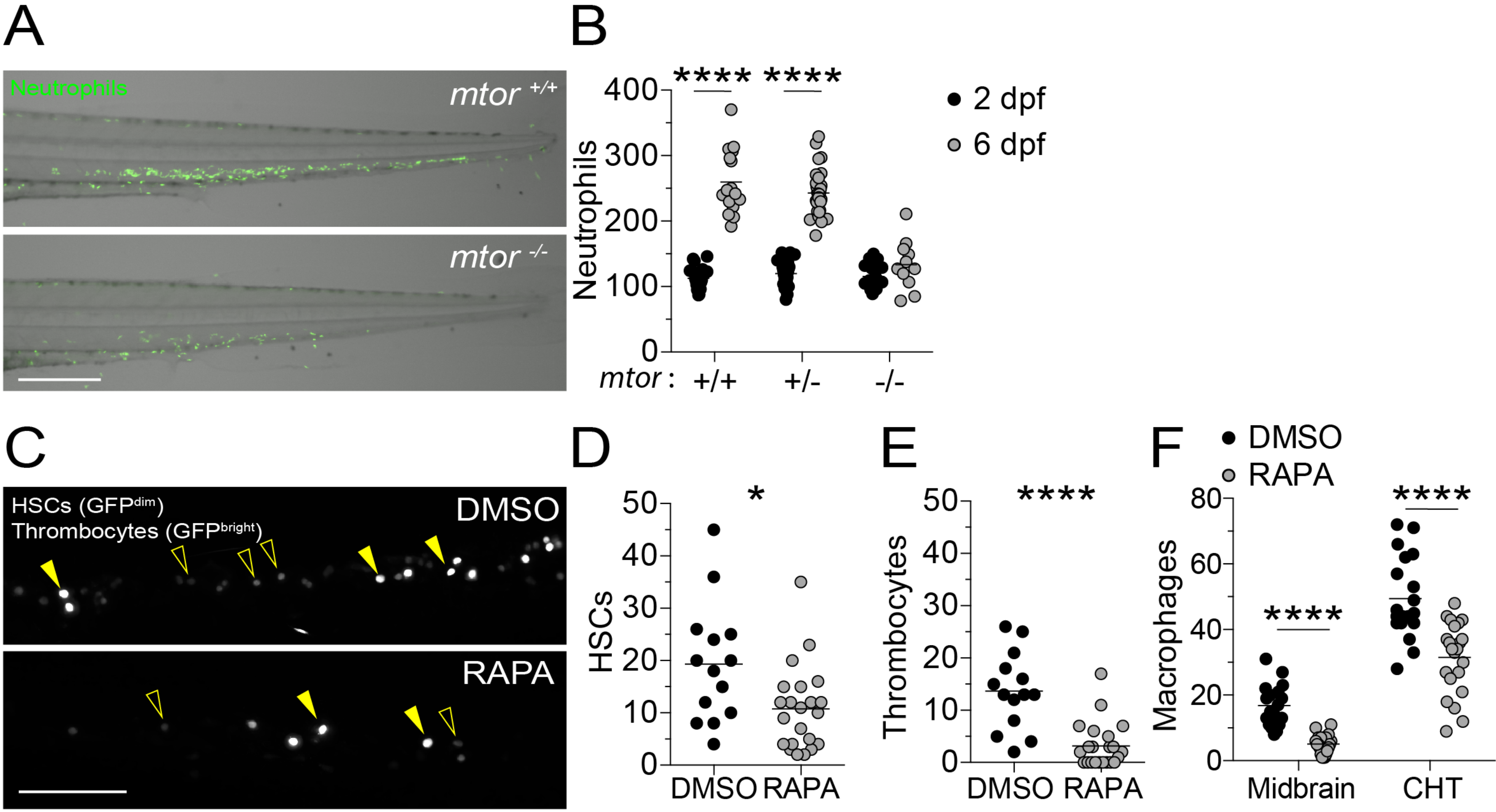

### Figure S2

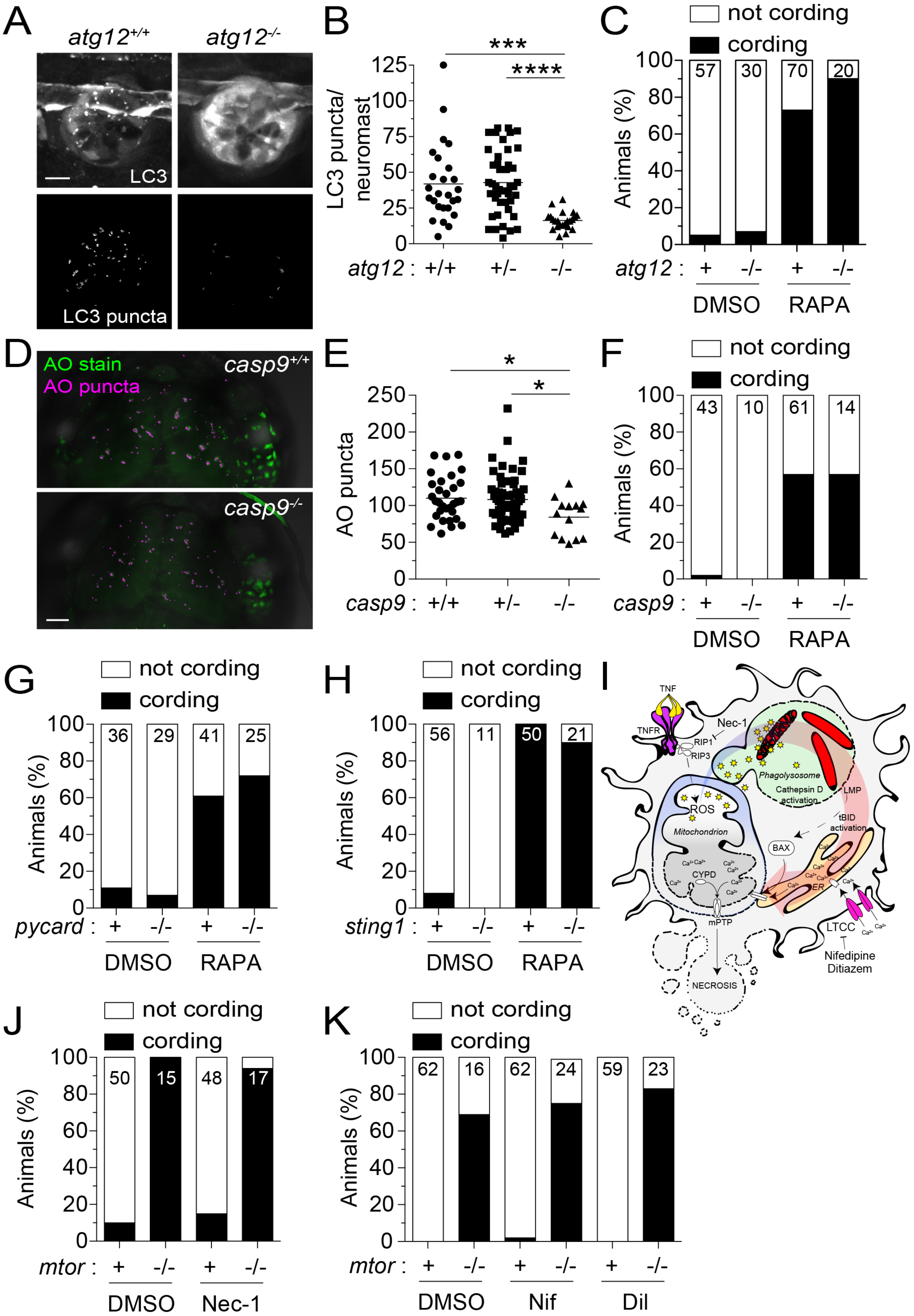

### Figure S3

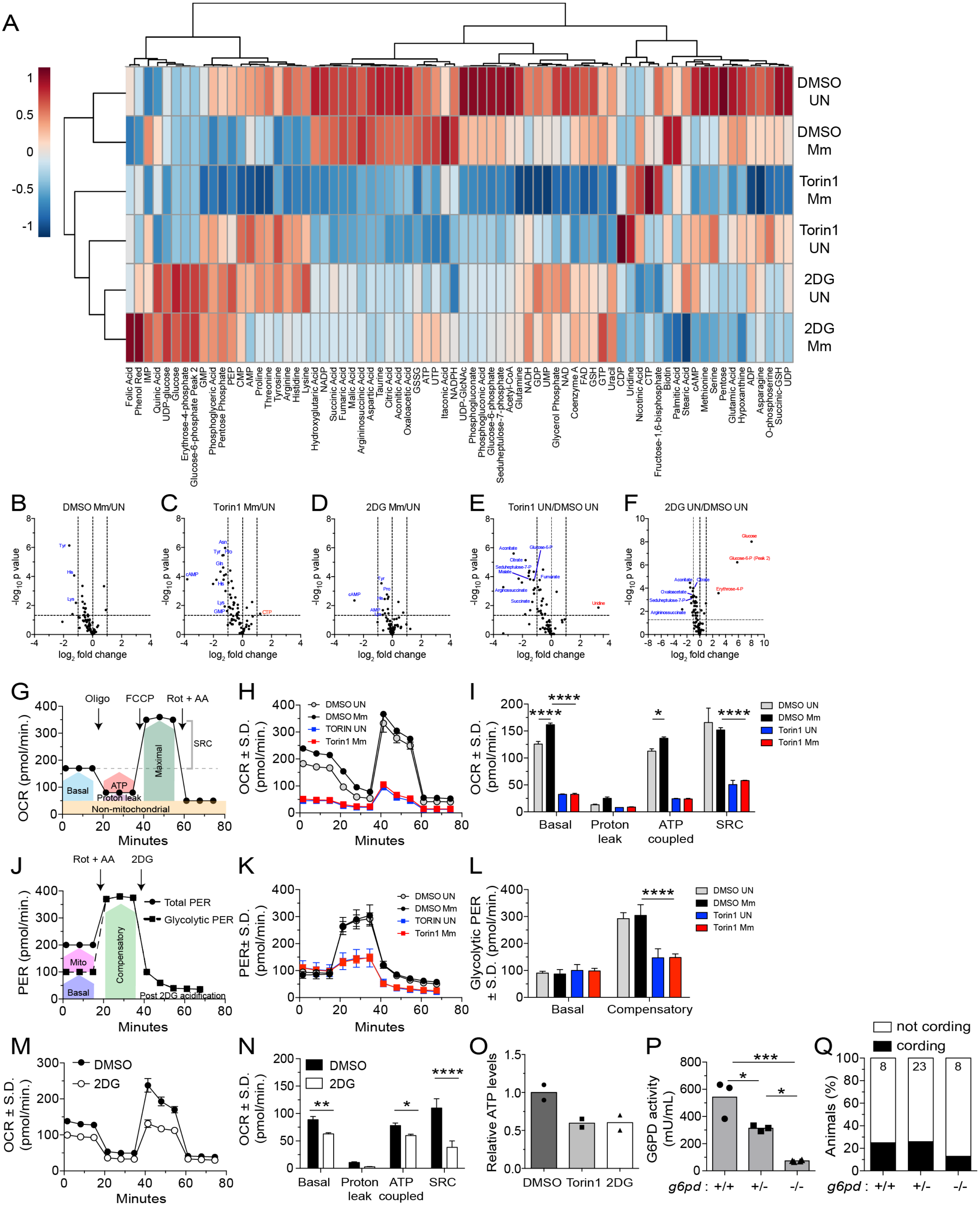

### Figure S4

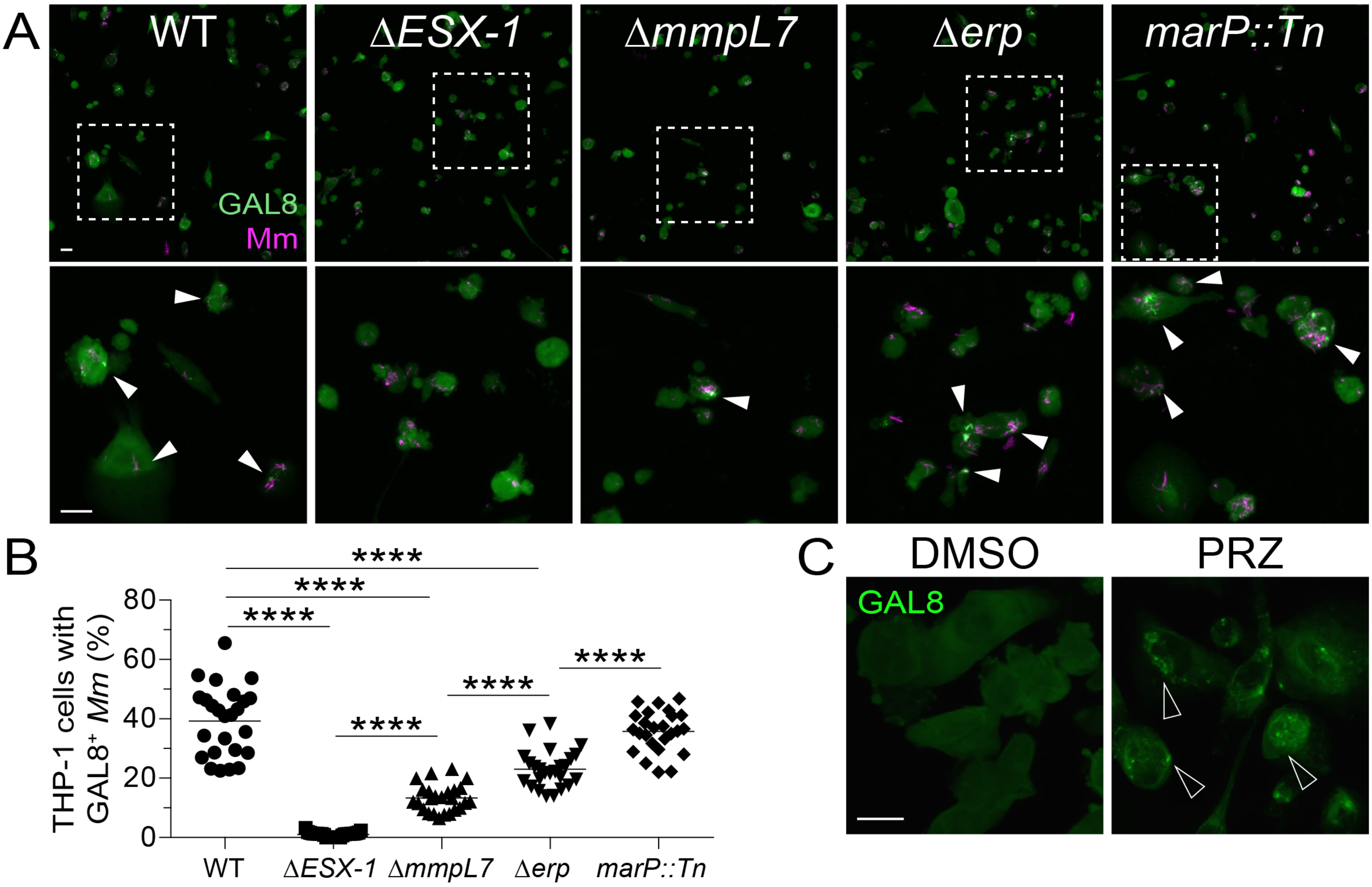
