## Supplementary material for "mTOR-regulated Mitochondrial Metabolism Limits Mycobacterium-induced Cytotoxicity": Table S1

**Table S1. Metabolic Profiles of Uninfected and Mm-infected THP-1 Macrophages Treated with Torin1, 2DG, or DMSO, Related to Figure S3.**

|  | **Normalized Counts Per Biological Replicate.** | | | | | | | | | | | | | | | | | | | | | | | |
| --- | --- | --- | --- | --- | --- | --- | --- | --- | --- | --- | --- | --- | --- | --- | --- | --- | --- | --- | --- | --- | --- | --- | --- | --- |
| **Metabolites** | **n01**  **UN DMSO** | **n02**  **UN DMSO** | **n03**  **UN DMSO** | **n04**  **UN DMSO** | **n05**  **UN Torin 1** | **n06**  **UN Torin 1** | **n07**  **UN Torin 1** | **n08**  **UN Torin 1** | **n09**  **UN 2DG** | **n10**  **UN 2DG** | **n11**  **UN 2DG** | **n12**  **UN 2DG** | **n13**  **Mm DMSO** | **n14**  **Mm DMSO** | **n15**  **Mm DMSO** | **n16**  **Mm DMSO** | **n17**  **Mm Torin1** | **n18**  **Mm Torin1** | **n19**  **Mm Torin1** | **n20**  **Mm Torin1** | **n21**  **Mm 2DG** | **n22**  **Mm 2DG** | **n23**  **Mm 2DG** | **n24**  **Mm 2DG** |
| **Acetyl CoA** | 52491 | 30385 | 38099 | 40894 | 22501 | 21678 | 27794 | 29051 | 24868 | 26499 | 27482 | 32089 | 29255 | 31323 | 36401 | 19537 | 21318 | 17097 | 15824 | 22298 | 33682 | 27606 | 24343 | 30240 |
| **Aconitic Acid** | 1365926 | 1066801 | 1168854 | 1121209 | 164720 | 198692 | 185748 | 223063 | 358255 | 467436 | 384625 | 446899 | 921138 | 1088699 | 1138256 | 759516 | 257359 | 302472 | 290579 | 371092 | 486800 | 412393 | 393118 | 564892 |
| **ADP** | 1345310 | 605373 | 842051 | 642504 | 597317 | 643032 | 725953 | 774759 | 518303 | 738896 | 739796 | 905147 | 661895 | 609531 | 833873 | 594090 | 325517 | 255169 | 287870 | 268836 | 599148 | 487781 | 510803 | 555942 |
| **AMP** | 641560 | 186613 | 318324 | 160147 | 347877 | 312646 | 473996 | 348814 | 209674 | 330418 | 332269 | 366911 | 278462 | 163785 | 301333 | 282974 | 133038 | 73133 | 86245 | 72840 | 225889 | 169845 | 193303 | 197510 |
| **Arginine** | 649868 | 755097 | 662466 | 739952 | 569815 | 672042 | 664626 | 688534 | 628624 | 697860 | 635388 | 599645 | 463597 | 454960 | 463458 | 347679 | 332130 | 385117 | 404983 | 420951 | 439693 | 507445 | 530867 | 727001 |
| **Argininosuccinic Acid** | 7093 | 4878 | 11259 | 6092 | 0 | 0 | 0 | 0 | 0 | 0 | 2598 | 1832 | 7847 | 11152 | 11402 | 2973 | 0 | 0 | 0 | 0 | 4320 | 0 | 0 | 6134 |
| **Asparagine** | 4067125 | 3321188 | 3229438 | 3261043 | 2972013 | 3092156 | 3142511 | 3346368 | 2334617 | 3164841 | 2603801 | 3092999 | 2923208 | 2878899 | 3438850 | 2272626 | 1487272 | 1276696 | 1363581 | 1336634 | 2574233 | 2390281 | 2430101 | 3105632 |
| **Aspartic Acid** | 6697457 | 4165832 | 5171314 | 4475326 | 879255 | 995256 | 1145266 | 1243230 | 1742045 | 2258211 | 1844763 | 2395504 | 4349900 | 5139298 | 6229693 | 3424719 | 841818 | 765794 | 890780 | 893047 | 1977825 | 1462858 | 1444019 | 1703390 |
| **ATP** | 1113271 | 335999 | 770413 | 875623 | 195911 | 226564 | 379108 | 319638 | 464633 | 256557 | 301168 | 611807 | 1249801 | 1134884 | 550969 | 318621 | 345311 | 235658 | 375576 | 569830 | 1015546 | 766772 | 310305 | 387429 |
| **Biotin** | 12897 | 13457 | 18982 | 8935 | 13261 | 9904 | 17898 | 6837 | 12184 | 5415 | 6692 | 9606 | 19250 | 13333 | 23633 | 14449 | 5769 | 5725 | 9493 | 7586 | 3301 | 3707 | 9700 | 0 |
| **cAMP** | 1288609 | 307072 | 559915 | 151356 | 363248 | 309212 | 504514 | 352184 | 176748 | 466103 | 446456 | 504477 | 156184 | 70093 | 162686 | 158343 | 27793 | 28982 | 27722 | 24894 | 71266 | 55001 | 85425 | 47329 |
| **CDP** | 52442 | 22821 | 28830 | 24977 | 56504 | 55975 | 65839 | 65038 | 23194 | 28114 | 26508 | 37331 | 22703 | 21379 | 24785 | 23891 | 35424 | 25221 | 36500 | 26973 | 21602 | 17665 | 19471 | 21715 |
| **Citric Acid** | 35180074 | 27684255 | 30517541 | 31006344 | 7917489 | 9476379 | 8047761 | 10666761 | 13560053 | 16890978 | 14991753 | 16395142 | 25417778 | 31517297 | 30477625 | 21672612 | 8965036 | 10629868 | 11589476 | 13738032 | 19345409 | 16479324 | 14225520 | 20057446 |
| **CMP** | 80197 | 37994 | 55628 | 46689 | 48031 | 54827 | 72170 | 67093 | 41319 | 54743 | 51448 | 71638 | 36556 | 43297 | 43382 | 22870 | 25750 | 26026 | 25311 | 30393 | 67194 | 46941 | 38790 | 54681 |
| **CoenzymeA** | 15228 | 9161 | 13308 | 12524 | 7747 | 8889 | 8532 | 5565 | 11469 | 8668 | 10907 | 10924 | 11559 | 10723 | 11403 | 8748 | 7991 | 6209 | 8256 | 7273 | 12500 | 11176 | 10278 | 10335 |
| **CTP** | 69121 | 18244 | 49024 | 52875 | 24375 | 31236 | 50427 | 49999 | 31968 | 16566 | 19380 | 44640 | 80154 | 71312 | 34257 | 17398 | 73582 | 53809 | 96381 | 138308 | 69696 | 55267 | 17333 | 25849 |
| **Erythrose-4-phosphate** | 16439 | 2817 | 4550 | 0 | 0 | 0 | 0 | 0 | 39837 | 55307 | 43318 | 40547 | 0 | 0 | 0 | 0 | 0 | 0 | 0 | 0 | 58485 | 36430 | 41111 | 40311 |
| **FAD** | 97398 | 71821 | 85351 | 71567 | 44391 | 43652 | 59789 | 58753 | 60186 | 72569 | 70443 | 76248 | 68611 | 66837 | 92946 | 62977 | 39941 | 40619 | 40043 | 38230 | 74567 | 61763 | 71699 | 72621 |
| **Folic Acid** | 7497 | 7345 | 7972 | 10466 | 424 | 8521 | 6870 | 15370 | 6681 | 9122 | 7564 | 6461 | 0 | 8271 | 3233 | 1279 | 4672 | 1855 | 8807 | 12008 | 11652 | 8762 | 8838 | 29178 |
| **Fructose-1,6-bisphosphate** | 31326 | 13554 | 29479 | 28274 | 14461 | 11799 | 25540 | 16572 | 12482 | 11168 | 9177 | 16449 | 23142 | 25621 | 21165 | 8950 | 23457 | 22178 | 28225 | 37609 | 18947 | 18652 | 10499 | 14925 |
| **Fumaric Acid** | 1605901 | 1246802 | 1396122 | 1312802 | 768150 | 751692 | 808149 | 848563 | 937674 | 1081708 | 1000311 | 997476 | 1361699 | 1531572 | 1515359 | 1137145 | 692795 | 743090 | 712828 | 854976 | 1065411 | 917056 | 895473 | 1060763 |
| **GDP** | 59734 | 15094 | 33650 | 28789 | 22459 | 20686 | 34480 | 24844 | 28463 | 28295 | 29562 | 50614 | 32892 | 25275 | 29700 | 22744 | 18261 | 12373 | 13844 | 12889 | 36900 | 30804 | 21545 | 24116 |
| **Glucose** | 184720 | 135539 | 144804 | 151158 | 94533 | 81441 | 91451 | 93740 | 39010824 | 42692289 | 39338172 | 41989983 | 139813 | 146726 | 141761 | 126635 | 69398 | 68598 | 71921 | 101814 | 35966614 | 30928260 | 33624793 | 34442668 |
| **Glucose-6-phosphate** | 3071821 | 2081789 | 2457768 | 2390771 | 894608 | 965799 | 1258652 | 1144202 | 1121845 | 1335857 | 1257922 | 1419597 | 1707834 | 1776537 | 2118072 | 1257570 | 510083 | 561794 | 618915 | 666001 | 1508856 | 1262499 | 1204491 | 1426724 |
| **Glucose-6-phosphate Peak2** | 123119 | 124350 | 98656 | 99205 | 67462 | 90931 | 66515 | 100064 | 5532689 | 6629624 | 6341225 | 6811482 | 84672 | 111946 | 89902 | 68670 | 39482 | 44307 | 77571 | 47646 | 6928282 | 6181670 | 6452547 | 6401583 |
| **Glutamic Acid** | 30105918 | 23098356 | 26127293 | 23674515 | 13497861 | 16508037 | 18957695 | 19867578 | 12316755 | 15350090 | 13889308 | 14441351 | 20816778 | 20973679 | 24017283 | 17309295 | 9125680 | 9305947 | 10570804 | 10925876 | 14533800 | 10880933 | 9167533 | 13416859 |
| **Glutamine** | 6910801 | 6675434 | 6390529 | 7181325 | 4033138 | 4572300 | 5003811 | 4612674 | 4160699 | 5560552 | 4458007 | 5180935 | 5268127 | 5559010 | 5318208 | 3067465 | 1555223 | 1724759 | 1815836 | 2153905 | 4419534 | 4363130 | 3948767 | 6187370 |
| **Glycerol phosphate** | 4489539 | 3870426 | 3969160 | 3591272 | 2880003 | 3123120 | 3194484 | 3447684 | 3485460 | 3946086 | 3740399 | 3921140 | 3484465 | 3281221 | 3946163 | 3206830 | 3211940 | 2784212 | 2895870 | 2764461 | 3732257 | 3385108 | 3480522 | 3695836 |
| **GMP** | 12717 | 6561 | 5632 | 8500 | 10978 | 6716 | 14392 | 8252 | 11062 | 10139 | 11960 | 12336 | 8230 | 4007 | 6993 | 6936 | 5986 | 3644 | 5067 | 4618 | 9107 | 10189 | 10287 | 11873 |
| **GSH** | 51021305 | 39195607 | 42318292 | 38168290 | 18070110 | 21189790 | 23801826 | 26039607 | 27548050 | 35157557 | 31814268 | 34954517 | 35238978 | 38493945 | 45655005 | 27993357 | 15300343 | 13387809 | 15464908 | 15742196 | 38481047 | 28664774 | 30087268 | 36620555 |
| **GSSG** | 3796772 | 2958720 | 3669246 | 3536803 | 2220788 | 1744730 | 2249447 | 2064274 | 2645831 | 2876438 | 2486541 | 2946976 | 3835303 | 3366855 | 4228071 | 2904111 | 2046274 | 1689338 | 1668389 | 1557847 | 3015814 | 2916438 | 3137044 | 2873379 |
| **GTP** | 14638 | 8902 | 9171 | 10879 | 4997 | 7262 | 6682 | 4752 | 9398 | 9953 | 9803 | 9785 | 14442 | 11567 | 11941 | 10263 | 6186 | 6479 | 5791 | 5887 | 16158 | 13693 | 14931 | 8735 |
| **Histidine** | 1082979 | 898075 | 897599 | 1001855 | 755017 | 925615 | 910887 | 1050898 | 904113 | 947875 | 925535 | 1103998 | 417371 | 466782 | 447177 | 286854 | 385826 | 299283 | 376652 | 413967 | 706912 | 674447 | 549693 | 761079 |
| **Hydroxyglutaric Acid** | 1605146 | 1053512 | 1356231 | 1185585 | 528484 | 547980 | 640492 | 632798 | 772712 | 883387 | 733714 | 831545 | 1069880 | 1167980 | 1348838 | 850680 | 457943 | 453728 | 496082 | 540522 | 866779 | 643726 | 751541 | 808587 |
| **Hypoxanthine** | 593300 | 548682 | 783568 | 471133 | 525270 | 366046 | 624698 | 418723 | 393186 | 453195 | 346842 | 325205 | 618113 | 390793 | 632754 | 451169 | 456638 | 379307 | 294840 | 265797 | 208028 | 322075 | 382416 | 263989 |
| **IMP** | 65524 | 48530 | 89636 | 56487 | 139350 | 68172 | 171692 | 59690 | 96559 | 129557 | 99057 | 97157 | 147262 | 79266 | 146764 | 116747 | 82884 | 74321 | 69911 | 47965 | 122114 | 135861 | 176136 | 92068 |
| **Itaconic Acid** | 433880 | 415357 | 486241 | 443682 | 401569 | 387045 | 434185 | 428319 | 450267 | 426129 | 426188 | 402279 | 500134 | 598792 | 395362 | 415112 | 326586 | 439538 | 474982 | 520563 | 472106 | 442492 | 433036 | 420801 |
| **Lysine** | 18030 | 25144 | 20124 | 16273 | 20587 | 19821 | 16854 | 12022 | 20264 | 26024 | 20959 | 18469 | 10960 | 4049 | 12579 | 8372 | 2249 | 10904 | 10117 | 6356 | 6858 | 11742 | 18648 | 16742 |
| **Malic Acid** | 13900229 | 9754925 | 11142769 | 10442481 | 3598439 | 3846706 | 4235548 | 4650461 | 5977579 | 7625603 | 6058228 | 6994597 | 10556580 | 12199093 | 12290554 | 8146952 | 3157056 | 3355548 | 3537423 | 4203536 | 7222512 | 5688630 | 5505901 | 7363775 |
| **Methionine** | 6767 | 3559 | 14770 | 6084 | 4449 | 2878 | 11604 | 4631 | 9465 | 3771 | 3854 | 0 | 10623 | 0 | 0 | 6226 | 4235 | 4544 | 509 | 2439 | 0 | 3197 | 4430 | 3366 |
| **NAD** | 19991 | 12426 | 18661 | 19633 | 10328 | 9180 | 13950 | 12026 | 16295 | 17680 | 15176 | 13827 | 11280 | 11991 | 12265 | 10451 | 9399 | 6671 | 5890 | 6674 | 13921 | 12342 | 11835 | 12721 |
| **NADH** | 132393 | 81257 | 83175 | 85010 | 72838 | 69619 | 89628 | 100057 | 72381 | 93738 | 85686 | 93702 | 69058 | 78574 | 83421 | 58646 | 48693 | 45112 | 51692 | 46970 | 99535 | 76733 | 84673 | 119763 |
| **NADP** | 26551 | 20491 | 23257 | 19172 | 10496 | 10671 | 13369 | 15398 | 9822 | 16220 | 17326 | 13976 | 19206 | 19041 | 25666 | 17840 | 10232 | 8949 | 8969 | 9945 | 15890 | 13171 | 13314 | 14852 |
| **NADPH** | 22887 | 11478 | 17474 | 20764 | 7551 | 9169 | 14930 | 11824 | 8752 | 6317 | 6269 | 11672 | 20451 | 24480 | 23745 | 11577 | 13594 | 10417 | 13994 | 13757 | 16851 | 13093 | 9936 | 15564 |
| **Nicotinic Acid** | 1059335 | 837720 | 893783 | 886559 | 1359764 | 1602453 | 1498501 | 1630017 | 596001 | 595156 | 492109 | 656583 | 1717812 | 1719728 | 1573784 | 1358121 | 1662734 | 1848390 | 1921691 | 2057986 | 836291 | 801940 | 692215 | 787793 |
| **O-phosphoserine** | 33020 | 42491 | 24322 | 25571 | 29758 | 31951 | 27529 | 30723 | 16213 | 31642 | 27431 | 20582 | 20050 | 27475 | 24960 | 35480 | 16652 | 18929 | 22978 | 17372 | 28587 | 11403 | 23923 | 27528 |
| **Oxaloacetic Acid** | 21929 | 17702 | 17208 | 22503 | 2473 | 7699 | 1958 | 7213 | 9349 | 10010 | 10754 | 9420 | 17505 | 18293 | 18531 | 13793 | 6806 | 7229 | 5799 | 9750 | 4047 | 10403 | 7522 | 14165 |
| **Palmitic Acid** | 693772 | 592521 | 555557 | 553255 | 553367 | 638377 | 531160 | 569843 | 604838 | 519230 | 593015 | 679691 | 739144 | 610178 | 649487 | 561751 | 532477 | 597405 | 550163 | 713668 | 527961 | 593520 | 528277 | 531487 |
| **Pentose** | 523804 | 417040 | 420550 | 387884 | 160460 | 190407 | 201582 | 200262 | 149486 | 150416 | 163898 | 166482 | 269739 | 269855 | 300938 | 225242 | 142518 | 156318 | 151765 | 149873 | 159457 | 129041 | 134553 | 147443 |
| **Pentose phosphate** | 176491 | 82557 | 110100 | 76496 | 92898 | 98208 | 122636 | 131241 | 98018 | 139253 | 133113 | 179009 | 60368 | 60594 | 67753 | 51948 | 34774 | 28160 | 27842 | 37132 | 140422 | 113063 | 132004 | 159519 |
| **PEP** | 82464 | 35468 | 54669 | 62311 | 29999 | 41233 | 46204 | 89023 | 47154 | 78272 | 68827 | 90755 | 16730 | 35487 | 50384 | 17904 | 14850 | 25602 | 31603 | 42487 | 72231 | 49423 | 51708 | 84056 |
| **Phenol Red** | 529964 | 481331 | 462027 | 545507 | 375305 | 472891 | 519406 | 545645 | 584670 | 550815 | 504187 | 589773 | 381040 | 468661 | 506426 | 385034 | 462101 | 528367 | 496847 | 595194 | 655802 | 633099 | 590500 | 843299 |
| **Phosphogluconate** | 83002 | 59899 | 60247 | 52009 | 22584 | 26421 | 22835 | 37265 | 22879 | 38630 | 32195 | 40502 | 35672 | 46313 | 55544 | 35267 | 9493 | 16558 | 21988 | 23916 | 37090 | 24642 | 23606 | 40376 |
| **Phosphogluconic Acid** | 85509 | 59899 | 60247 | 52968 | 22584 | 26421 | 22835 | 40912 | 22879 | 38630 | 30094 | 42807 | 40266 | 48838 | 56765 | 35267 | 9493 | 13236 | 22428 | 22784 | 37090 | 24642 | 27038 | 43441 |
| **Phosphoglyceric Acid** | 237235 | 119992 | 144246 | 165152 | 100900 | 163530 | 132871 | 298083 | 107985 | 190407 | 183114 | 217416 | 69082 | 139524 | 140932 | 75403 | 64574 | 78748 | 112582 | 158803 | 189851 | 138828 | 137848 | 243973 |
| **Proline** | 132103 | 112773 | 126946 | 104186 | 114102 | 121149 | 123076 | 116812 | 120867 | 124549 | 110608 | 123934 | 97007 | 78832 | 99654 | 67807 | 56817 | 41149 | 51891 | 41862 | 91864 | 85321 | 88286 | 103658 |
| **Quinic Acid** | 59462 | 44013 | 51659 | 86526 | 82748 | 48998 | 58302 | 45596 | 59103 | 179556 | 60454 | 52628 | 76722 | 74469 | 51349 | 99674 | 69774 | 77826 | 65005 | 72268 | 71859 | 84162 | 130009 | 47531 |
| **Seduheptulose-7-phosphate** | 427080 | 309921 | 350122 | 335498 | 98004 | 114544 | 146627 | 135417 | 150750 | 200177 | 163486 | 184467 | 244025 | 259922 | 312737 | 175512 | 76840 | 76760 | 81413 | 89369 | 213969 | 159678 | 172762 | 217993 |
| **Serine** | 659562 | 476112 | 558466 | 533995 | 444865 | 486965 | 531124 | 512024 | 306269 | 333343 | 312675 | 385924 | 401894 | 364687 | 366540 | 256648 | 285477 | 255836 | 264909 | 292394 | 294380 | 279992 | 247840 | 288540 |
| **Stearic Acid** | 615609 | 534184 | 525364 | 510662 | 570520 | 543606 | 560203 | 472450 | 498587 | 489123 | 566653 | 638767 | 627790 | 511651 | 501387 | 455010 | 433460 | 526776 | 540717 | 582336 | 458520 | 493533 | 490515 | 487530 |
| **Succinic Acid** | 416422 | 208378 | 310564 | 342333 | 111632 | 129745 | 117927 | 152512 | 219282 | 175909 | 213601 | 268921 | 333158 | 344058 | 297641 | 239831 | 104563 | 113232 | 97711 | 170799 | 182043 | 171980 | 195331 | 243360 |
| **Succinic-GSH** | 930773 | 754928 | 797913 | 780453 | 389944 | 671647 | 537138 | 811070 | 373674 | 566016 | 504351 | 545941 | 388818 | 675235 | 884358 | 484443 | 278145 | 282625 | 401219 | 431956 | 523879 | 410940 | 474459 | 566522 |
| **Taurine** | 15152334 | 12000021 | 14489756 | 12068878 | 4469507 | 3640031 | 5858496 | 4264058 | 6786172 | 6782866 | 5640582 | 6197108 | 13551039 | 11454646 | 12306518 | 9648989 | 4665512 | 4746916 | 4232683 | 3892172 | 5986053 | 5264281 | 6773268 | 5868020 |
| **Threonine** | 1602544 | 1380665 | 1483618 | 1468588 | 1436864 | 1285015 | 1589696 | 1245852 | 1334732 | 1505119 | 1266829 | 1473697 | 1222045 | 1007838 | 1138648 | 732251 | 652960 | 603967 | 617567 | 640876 | 1109156 | 1096809 | 1130459 | 1154041 |
| **Tyrosine** | 434714 | 441640 | 422528 | 435761 | 547592 | 527401 | 614358 | 537684 | 504622 | 567687 | 467279 | 526425 | 157431 | 156181 | 154452 | 103435 | 226547 | 191719 | 214060 | 223654 | 291974 | 284339 | 286490 | 361330 |
| **UDP** | 379397 | 212521 | 274043 | 254504 | 186523 | 240813 | 225634 | 287066 | 213004 | 210540 | 244595 | 259349 | 201865 | 259062 | 248196 | 216295 | 165552 | 173386 | 200317 | 201620 | 231151 | 221379 | 166425 | 223395 |
| **UDP-GlcNAc** | 19740574 | 13426879 | 16369004 | 14930371 | 3874731 | 4659947 | 6364619 | 6847348 | 5107493 | 7416184 | 6367045 | 7293974 | 8904025 | 11481825 | 15004340 | 6534458 | 2581519 | 2672325 | 3130267 | 3256257 | 7778300 | 5505639 | 5910920 | 7322553 |
| **UDP-glucose** | 2139876 | 1395633 | 1714553 | 1435666 | 474998 | 530674 | 632854 | 678729 | 2582143 | 3669065 | 3164476 | 3625370 | 1300718 | 1485346 | 2047951 | 1158124 | 372623 | 355646 | 391890 | 406178 | 4109012 | 2963298 | 3463207 | 4087035 |
| **UMP** | 111902 | 48815 | 73804 | 69134 | 45179 | 45463 | 55359 | 54303 | 54391 | 84099 | 71723 | 83081 | 58938 | 51893 | 80675 | 43909 | 19121 | 18861 | 13044 | 19090 | 94516 | 65108 | 58906 | 72306 |
| **Uracil** | 1827528 | 1346053 | 1478939 | 1355221 | 674321 | 712712 | 842613 | 838644 | 1464605 | 1452547 | 1499051 | 1580265 | 1342420 | 1403604 | 1466960 | 1051011 | 525438 | 529830 | 539444 | 586638 | 1774102 | 1476522 | 1425117 | 1626315 |
| **Uridine** | 308070 | 281713 | 268641 | 297133 | 2349693 | 2761514 | 2920161 | 2977613 | 177944 | 215422 | 178552 | 216786 | 222642 | 275776 | 248491 | 178118 | 1957841 | 2330106 | 2167868 | 2323961 | 193186 | 188321 | 163558 | 285673 |
| **UTP** | 121599 | 33843 | 83938 | 98841 | 21079 | 19488 | 29872 | 25903 | 74007 | 43414 | 51766 | 57870 | 154566 | 136183 | 68850 | 38225 | 28747 | 16531 | 24865 | 60757 | 107171 | 80696 | 54154 | 54138 |
