## Supplementary material for "mTOR-regulated Mitochondrial Metabolism Limits Mycobacterium-induced Cytotoxicity": Table S2

**Table S2. KEY RESOURCES TABLE**

| REAGENT or RESOURCE | SOURCE | IDENTIFIER |
| --- | --- | --- |
| Antibodies | | |
| AlexaFluor 488 anti-cytochrome c (clone 6H2.B4) | BioLegend | Cat# 612308;  RRID:AB_2565240 |
| Phospho-S6 Ribosomal Protein (S235/S236) XP rabbit (clone D57.2.2E) (Alexa Fluor 647 Conjugate) | Cell Signaling Technology | Cat# 4851;  RRID:AB_10695457 |
| S6 Ribosomal Protein (clone 54D2) Mouse mAb (Alexa Fluor 647 Conjugate) | Cell Signaling Technology | Cat# 5548;  RRID:AB_10707322 |
| Goat IgG anti-human Galectin-8 | R&D Systems | Cat# AF1305;  RRID:AB_2137229 |
| Donkey anti-goat IgG (H+L), Alexa Fluor 488 | Thermo Fisher Scientific | Cat# A-11055; RRID:AB_2534102 |
| Optical-bottom tissue culture plates | | |
| 96-well (half area) black plate with transparent bottom | Greiner Bio-One | Cat# 675090 |
| VisiPlate 24-well black plate with clear bottom | Perkin Elmer | Cat# 1450–606 |
| 6-well No. 1.5 coverslip, 20 mm glass diameter, uncoated plate | MatTek | Cat# P06G-1.5-20-F |
| 24-well No. 1.5 coverslip, 13 mm glass diameter, uncoated plate | MatTek | Cat# P24G-1.5-13-F |
| Bacterial and virus strains | | |
| *M. marinum* M strain transformed with *pmsp12::BFP2* | (Takaki et al., 2013) | Derivative of ATCC # BAA-535 |
| *M. marinum* M strain transformed with *pmsp12::mWasabi* | (Takaki *et al.*, 2013) | Derivative of ATCC # BAA-535 |
| *M. marinum* M strain transformed with *pmsp12::tdTomato* | (Takaki *et al.*, 2013) | Derivative of ATCC # BAA-535 |
| *M. marinum* M strain transformed with *pmsp12::tdKatushka2* | (Takaki *et al.*, 2013) | Derivative of ATCC # BAA-535 |
| *ΔESX1 M. marinum* M strain transformed with *pmsp12::tdTomato* | (Pagan et al., 2015) | Derivative of ATCC # BAA-535 |
| *ΔmmpL7 M.marinum* M strain transformed with *pmsp12::tdTomato* | (Cambier et al., 2014) | Derivative of ATCC # BAA-535 |
| *Δerp M. marinum* M strain transformed with *pmsp12::tdTomato* | (Takaki *et al.*, 2013) | Derivative of ATCC # BAA-535 |
| *marP::Tn M. marinum* M strain transformed with *pmsp12::tdTomato* | (Levitte et al., 2016) | Derivative of ATCC # BAA-535 |
| *ΔesxA M. marinum* M strain transformed with *pmsp12::tdTomato* | (Osman et al., 2022) | Derivative of ATCC # BAA-535 |
| *ΔesxA M. marinum* M strain transformed with *::esxA*^WT^ | (Osman *et al.*, 2022) | Derivative of ATCC # BAA-535 |
| *ΔesxA M. marinum* M strain transformed with *::esxA*^M83I^ | (Osman *et al.*, 2022) | Derivative of ATCC # BAA-535 |
| *ΔesxA M. marinum* M strain transformed with *::esxA*^M93T^ | (Osman *et al.*, 2022) | Derivative of ATCC # BAA-535 |
| *M. tuberculosis ΔleuDΔpanCD* mc^2^ 6206 transformed with *pmsp12::tdTomato* | (Roca et al., 2019) | N/A |
| Chemicals, peptides, and recombinant proteins | | |
| BD Difco Middlebrook 7H9 broth (dehydrated) | Fisher Scientific | Cat# DF0713-17-9 |
| Middlebrook 7H10 agar base | Sigma-Aldrich | Cat# M0303 |
| L-leucine | Sigma-Aldrich | Cat# L8000;  CAS: 61-90-5 |
| Calcium pantothenate | Sigma-Aldrich | Cat# C8731;  CAS: 137-08-6 |
| Remel OADC Enrichment | Fisher Scientific | Cat# 11903262 |
| RPMI 1640 medium | Sigma-Aldrich | Cat# R7509 |
| Gibco Phenol-free RPMI | Thermo-Fisher | Cat# 11835030 |
| XF DMEM, pH 7.4 | Agilent | Cat# 103575-100 |
| Instant Ocean Salt | ZM Systems | N/A |
| PTU (1-phenyl-2-thiourea) | Sigma-Aldrich | Cat# P7629;  CAS: 103-85-5 |
| Tango Buffer (10x) | Thermo Fisher | Cat# BY5 |
| Phenol Red Sodium Salt | Sigma-Aldrich | Cat# P5530;  CAS: 34487-61-1 |
| Pronase | Sigma-Aldrich | Cat# P5147;  CAS:9036-06-0 |
| Tricaine (ethyl 3-amonobenzoate, methanesulfonic acid salt) | Fisher Scientific | Cat# 10743661;  CAS: 886-86-2 |
| TopVision Low Melt Agarose | Thermo Fisher | Cat# R0801 |
| Rapamycin | Sigma-Aldrich | Cat# R0395;  CAS: 53123-88-9 |
| Torin1 | Cambridge Bioscience | Cat# CAY10997;  CAS: 1222998-36-8 |
| Nifedipine | Cambridge Bioscience | Cat# N3228;  CAS: 21829-25-4 |
| Diltiazem HCl | Cambridge Bioscience | Cat# D3447;  CAS: 33286-22-5 |
| 2DG (2-deoxy-D-glucose) | Sigma-Aldrich | Cat# D8375;  CAS: 154-17-6 |
| UK5099 | Cambridge Bioscience | Cat# B1952;  CAS: 56396-35-1 |
| Digitonin | Acros | Cat# 407565000  CAS: 11024-24-1 |
| Prazosin HCl | Sigma-Aldrich | Cat#: P7791;  CAS: 19237-84-4 |
| PMA (Phorbol 12-myristate 13-acetate) | Sigma-Aldrich | Cat#: P1585;  CAS: 16561-29-8 |
| Accutase | Sigma-Aldrich | Cat#: A6964 |
| Paraformaldehyde, 16% w/v | Alfa Aesar | Cat# 11490570 |
| Acridine Orange (2% solution in H_2_0) | Sigma-Aldrich | Cat# A9231;  CAS: 65-61-2 |
| MitoTracker Red CM-H_2_Xros | Thermo Fisher | Cat# M7513 |
| eBioscience Fixable Viability Dye eFluor 660 | Thermo Fisher | Cat# 65-0864-14 |
| Tetramethylrhodamine, ethyl ester | Abcam | Cat# ab113852 |
| SYTOX Green Nucleic Acid Stain | Thermo Fisher | Cat# S7020 |
| AMPure XP beads for PCR Purification | Beckman Coulter | Cat# A63881 |
| Precision Melt Supermix | BioRad | Cat# 172-5112 |
| KASP V4.0 2X Master Mix | LGC Biosearch | Cat# KBS-1016-002 |
| Mammalian Cell Lysis Buffer 5x | Abcam | Cat# ab179835 |
| Critical commercial assays | | |
| Gibson Assembly Cloning Kit | New England BioLabs | Cat# E5510S |
| mMessage mMachine T7 Transcription Kit | Thermo Fisher | Cat# AM1344 |
| Seahorse XFp Glycolytic Rate Assay Kit | Agilent | Cat# 103346-100 |
| Seahorse XFp Mito Stress Test Kit | Agilent | Cat#103015-100 |
| ATP Assay Kit (Colorimetric/Fluorometric) | Abcam | Cat# ab83355 |
| Glucose-6-Phosphate Dehydrogenase Activity Assay Kit (Fluorometric) | Abcam | Cat# ab176722 |
| Experimental models: Cell lines | | |
| THP-1 | ATCC | Cat# TIB-202, RRID:CVCL_0006 |
| Experimental models: Organisms/strains | | |
| Zebrafish (*Danio rerio*): wild type AB strain | University of Cambridge | ZDB-GENO-960809-7 |
| Zebrafish: TL strain | University of Cambridge | ZDB-GENO-990623-2 |
| Zebrafish: WIK strain | University of Washington | ZDB-GENO-010531-2 |
| Zebrafish: *Tg(mpeg1:YFP)^w200^* | (Roca and Ramakrishnan, 2013) | ZDB-ALT-130130-3 |
| Zebrafish: *Tg(mpeg1:Brainbow)^w201^* | (Pagan *et al.*, 2015) | ZDB-ALT-150512-3 |
| Zebrafish: *Tg(mfap4:tdTomato-CAAX)^xt6^* | (Walton et al., 2015) | ZDB-ALT-160122-3 |
| Zebrafish: *Tg(ubib:secA5-YFP)^cu34^* | This work | N/A |
| Zebrafish: *Tg(CMV:EGFP-map1lc3b)^zf155^* | (He et al., 2009) | ZDB-ALT-091029-2 |
| Zebrafish: *Tg(lysC:EGFP)^nz117^* | (Hall et al., 2007) | ZDB-ALT-071109-2 |
| Zebrafish: *Tg(cd41:GFP)* | (Lin et al., 2005) | N/A |
| Zebrafish: *pycard^w216^* | (Matty et al., 2019) | ZDB-ALT-191009-1 |
| Zebrafish: *mtor^fh178^* | This work | N/A |
| Zebrafish: *mtor^sa16755^* | Wellcome Trust Sanger Institute | ZDB-ALT-131217-12934 |
| Zebrafish: *rptor^sa11537^* | Wellcome Trust Sanger Institute | ZDB-ALT-130530-177 |
| Zebrafish: *rictora^sa15967^* | Wellcome Trust Sanger Institute | ZDB-ALT-130411-4494 |
| Zebrafish: *rictorb^sa18403^* | Wellcome Trust Sanger Institute | ZDB-ALT-131217-14421 |
| Zebrafish: *atg12^sa42684^* | Wellcome Trust Sanger Institute | ZDB-ALT-160601-8392 |
| Zebrafish: *casp9^sa11164^* | Wellcome Trust Sanger Institute | ZDB-ALT-130411-1023 |
| Zebrafish: *sting1^sa35634^* | Wellcome Trust Sanger Institute | ZDB-ALT-160601-4021 |
| Zebrafish: *g6pd^sa24272^* | Wellcome Trust Sanger Institute | ZDB-ALT-161003-11894 |
| Oligonucleotides | | |
| *mtor^fh178^* forward primer for genotyping by HRMA  5’-TCACAGTATCAGATCTTCATTCCTATGGT-3’ | This paper | N/A |
| *mtor^fh178^* reverse primer for genotyping by HRMA  5’-ACATCATAGCGCTGGTGATTGAT-3’ | This work | N/A |
| *mtor^sa16755^* forward primer for genotyping by HRMA  5’-TGACTACAGCACCAGCGAGA-3’ | This work | N/A |
| *mtor^sa16755^* reverse primer for genotyping by HRMA  5’-ATGGTGTGGTGATTGGACAG-3’ | This work | N/A |
| Alt-R crRNA Dr.Cas9.NDUFAF1.1.AA  5’-TGGAACAGACCCGTGTCGTG-3’ | IDT | N/A |
| Alt-R crRNA Dr.Cas9.NDUFAF1.1.AB  5’-TATCGAGTCTCTCCATCACG-3’ | IDT | N/A |
| Alt-R crRNA Dr.Cas9.NDUFAF1.1.AC  5’-GGTCCCATACAGCAAACACG-3’ | IDT | N/A |
| Alt-R crRNA Dr.Cas9.NDUFAF1.1.AD  5’-CGTGTGTCTGGAGGCTGACC-3’ | IDT | N/A |
| Alt-R tracrRNA | IDT | N/A |
| Forward primer for genotyping *ndufaf1* AA mutagenesis by HRMA  5’-AGAGCACATGCTGGAACAGA-3’ | This work | N/A |
| Reverse primer for genotyping *ndufaf1* AA mutagenesis by HRMA  5’-TGTTTTTGCCCAGACTGACA-3’ | This work | N/A |
| Forward primer for genotyping *ndufaf1* AB, AC mutagenesis by HRMA  5’-CGCAGTGTGGCTTATGTCAG-3’ | This work | N/A |
| Reverse primer for genotyping *ndufaf1* AB, AC mutagenesis by HRMA  5’-TTGGAGCGCATAGAGCAGTA-3’ | This work | N/A |
| Forward primer for genotyping *ndufaf1* AD mutagenesis by HRMA  5’-AGGAGCAAGTTTGAGCGAGA-3’ | This work | N/A |
| Reverse primer for genotyping *ndufaf1* AD mutagenesis by HRMA  5’-AAGCCAAAGTGCTTCCTGAC-3’ | This work | N/A |
| pDestTol2pA2_ubi:EGFP forward primer for vector fragment amplification for Gibson Assembly  5’-GGCGGTGGAAGATCTGGG-3’ | This work | N/A |
| pDestTol2pA2_ubi:EGFP reverse primer for vector fragment amplification for Gibson Assembly  5’-GGTCCAGCCTGCTTTTTTG-3’ | This work | N/A |
| secA5-YFP forward primer for insert fragment amplification for Gibson Assembly  5’-aaagcaggctggaccATGCATAAGGTTTTGCTG-3’ | This work | N/A |
| secA5-YFP reverse primer for insert fragment amplification for Gibson Assembly  5’-agatcttccaccgccGATGAATTAATTCGAGCTCC-3’ | This work | N/A |
| ubb:secA5 joint sequence forward primer to validate assembled product  5’-TCGTTTAACATGGGAGAAGTGC-3’ | This work | N/A |
| ubb:secA5 joint sequence reverse primer to validate assembled product  5’-AGCCTTTCATAGCCTTCCGA-3’ | This work | N/A |
| YFP-SV40pA joint sequence forward primer to validate assembled product  5’-CTGTACAAGTAAAGCGGCCG-3’ | This work | N/A |
| YFP-SV40pA joint sequence reverse primer to validate assembled product  5’-GTAAAACGACGGCCAGTGAA-3’ | This work | N/A |
| Recombinant DNA and Proteins | | |
| T7-TPase | (Khattak et al., 2014) | RRID:Addgene_51818 |
| pDestTol2pA2_ubi:EGFP | (Mosimann et al., 2011) | RRID:Addgene_27323 |
| pBH-UAS-secA5-YFP | (van Ham et al., 2010) | RRID:Addgene_32359 |
| pTol2-ubb:secA5-YFP | This work | N/A |
| Alt-R Sp Cas9 Nuclease V3 | IDT | N/A |
| Software and algorithms |  |  |
| NIS Elements (5.21) | Nikon | N/A |
| IMARIS (8.2) and IMARIS for Cell Biologists (9.1) | Bitplane | N/A |
| FlowJo 10 | TreeStar | N/A |
| Prism (versions 7 and 9) | GraphPad | N/A |
| ImageJ | https://imagej.nih.gov/ij/ | N/A |
| Fluorescent Pixel Count Macro (Image J) | (Takaki *et al.*, 2013) | N/A |
| AssayR | (Wills et al., 2017) | N/A |
| MetaboAnalyst 5.0 | (Pang et al., 2021) | N/A |
| Photoshop CS6 | Adobe | N/A |
| Illustrator CS6 | Adobe | N/A |

REFERENCES

Cambier, C.J., Takaki, K.K., Larson, R.P., Hernandez, R.E., Tobin, D.M., Urdahl, K.B., Cosma, C.L., and Ramakrishnan, L. (2014). Mycobacteria manipulate macrophage recruitment through coordinated use of membrane lipids. Nature *505*, 218-222. 10.1038/nature12799.

Hall, C., Flores, M.V., Storm, T., Crosier, K., and Crosier, P. (2007). The zebrafish lysozyme C promoter drives myeloid-specific expression in transgenic fish. BMC Dev Biol *7*, 42. 10.1186/1471-213X-7-42.

He, C., Bartholomew, C.R., Zhou, W., and Klionsky, D.J. (2009). Assaying autophagic activity in transgenic GFP-Lc3 and GFP-Gabarap zebrafish embryos. Autophagy *5*, 520-526. 10.4161/auto.5.4.7768.

Khattak, S., Murawala, P., Andreas, H., Kappert, V., Schuez, M., Sandoval-Guzman, T., Crawford, K., and Tanaka, E.M. (2014). Optimized axolotl (Ambystoma mexicanum) husbandry, breeding, metamorphosis, transgenesis and tamoxifen-mediated recombination. Nature protocols *9*, 529-540. 10.1038/nprot.2014.040.

Levitte, S., Adams, K.N., Berg, R.D., Cosma, C.L., Urdahl, K.B., and Ramakrishnan, L. (2016). Mycobacterial Acid Tolerance Enables Phagolysosomal Survival and Establishment of Tuberculous Infection In Vivo. Cell Host Microbe *20*, 250-258. 10.1016/j.chom.2016.07.007.

Lin, H.F., Traver, D., Zhu, H., Dooley, K., Paw, B.H., Zon, L.I., and Handin, R.I. (2005). Analysis of thrombocyte development in CD41-GFP transgenic zebrafish. Blood *106*, 3803-3810. 10.1182/blood-2005-01-0179.

Matty, M.A., Knudsen, D.R., Walton, E.M., Beerman, R.W., Cronan, M.R., Pyle, C.J., Hernandez, R.E., and Tobin, D.M. (2019). Potentiation of P2RX7 as a host-directed strategy for control of mycobacterial infection. Elife *8*. 10.7554/eLife.39123.

Mosimann, C., Kaufman, C.K., Li, P., Pugach, E.K., Tamplin, O.J., and Zon, L.I. (2011). Ubiquitous transgene expression and Cre-based recombination driven by the ubiquitin promoter in zebrafish. Development *138*, 169-177. 10.1242/dev.059345.

Osman, M.M., Shanahan, J.K., Chu, F., Takaki, K.K., Pinckert, M.L., Pagan, A.J., Brosch, R., Conrad, W.H., and Ramakrishnan, L. (2022). The C terminus of the mycobacterium ESX-1 secretion system substrate ESAT-6 is required for phagosomal membrane damage and virulence. Proc Natl Acad Sci U S A *119*, e2122161119. 10.1073/pnas.2122161119.

Pagan, A.J., Yang, C.T., Cameron, J., Swaim, L.E., Ellett, F., Lieschke, G.J., and Ramakrishnan, L. (2015). Myeloid Growth Factors Promote Resistance to Mycobacterial Infection by Curtailing Granuloma Necrosis through Macrophage Replenishment. Cell Host Microbe *18*, 15-26. 10.1016/j.chom.2015.06.008.

Pang, Z., Chong, J., Zhou, G., de Lima Morais, D.A., Chang, L., Barrette, M., Gauthier, C., Jacques, P.E., Li, S., and Xia, J. (2021). MetaboAnalyst 5.0: narrowing the gap between raw spectra and functional insights. Nucleic Acids Res *49*, W388-W396. 10.1093/nar/gkab382.

Roca, F.J., and Ramakrishnan, L. (2013). TNF dually mediates resistance and susceptibility to mycobacteria via mitochondrial reactive oxygen species. Cell *153*, 521-534. 10.1016/j.cell.2013.03.022.

Roca, F.J., Whitworth, L.J., Redmond, S., Jones, A.A., and Ramakrishnan, L. (2019). TNF Induces Pathogenic Programmed Macrophage Necrosis in Tuberculosis through a Mitochondrial-Lysosomal-Endoplasmic Reticulum Circuit. Cell *178*, 1344-1361 e1311. 10.1016/j.cell.2019.08.004.

Takaki, K., Davis, J.M., Winglee, K., and Ramakrishnan, L. (2013). Evaluation of the pathogenesis and treatment of Mycobacterium marinum infection in zebrafish. Nature protocols *8*, 1114-1124. 10.1038/nprot.2013.068.

van Ham, T.J., Mapes, J., Kokel, D., and Peterson, R.T. (2010). Live imaging of apoptotic cells in zebrafish. FASEB J *24*, 4336-4342. 10.1096/fj.10-161018.

Walton, E.M., Cronan, M.R., Beerman, R.W., and Tobin, D.M. (2015). The Macrophage-Specific Promoter mfap4 Allows Live, Long-Term Analysis of Macrophage Behavior during Mycobacterial Infection in Zebrafish. PLoS One *10*, e0138949. 10.1371/journal.pone.0138949.

Wills, J., Edwards-Hicks, J., and Finch, A.J. (2017). AssayR: A Simple Mass Spectrometry Software Tool for Targeted Metabolic and Stable Isotope Tracer Analyses. Anal Chem *89*, 9616-9619. 10.1021/acs.analchem.7b02401.
